# Spatial Organization of Lipids Drives GPCR Conformational Equilibria

**DOI:** 10.64898/2026.06.11.731710

**Authors:** Anuradha V. Wijesekara, Jingjing Ji, Nessa P. Afsharian, Keen Zhang, Naveen Thakur, Arka P. Ray, Bar Elmaleh, Niloofar Gopal Pour, Ed Lyman, Matthew T. Eddy

## Abstract

Despite the widespread use of lipid nanodiscs for structure–function studies of membrane proteins, little is known about how lipids are spatially organized within nanodiscs and how this organization influences embedded proteins. The activity and conformational equilibria of the human A_2A_ adenosine receptor, a class A G protein-coupled receptor, are highly sensitive to anionic lipids that directly interact with the receptor. We leverage this lipid-dependent sensitivity to probe the accessibility of anionic lipids across nanodiscs of varying sizes. We identify a threshold concentration of anionic lipids required to fully populate active receptor conformations that is higher for POPS than for POPG. Computational simulations reveal that POPS and POPG each form lipid clusters reducing the effective availability of anionic lipids to interact with the receptor. This effect is stronger for POPS and scales with increasing nanodisc size, correlating with experimental biophysical and biochemical measurements. Simulations further identify positively charged residues within the membrane scaffold protein that coordinate anionic lipid headgroups. Targeted protein engineering reduces the threshold concentration of anionic lipids required for receptor activation, supporting strategies to control lipid accessibility within nanodiscs. Because membrane scaffold proteins are derived from apolipoprotein A-I, similar lipid–protein interactions may also influence lipid organization within biological systems such as HDL particles.

## Introduction

A longstanding challenge in structural and biochemical characterization of membrane proteins is the development of reconstitution systems that preserve native-like structure and activity while enabling experiments in aqueous solutions. Nanodiscs are soluble disc-like particles that have become one of the most widely used reconstitution systems for studies of integral membrane proteins and their complexes. Multiple types of amphipathic proteins and synthetic polymers have been developed for assembling lipids inside nanodiscs. This work focuses on nanodiscs formed from membrane scaffold proteins (MSPs), which were originally engineered from Apolipoprotein A1^1,2^ and subsequently optimized for different experimental applications.^3,4^ MSP-based nanodiscs are extensively employed in structural and biophysical studies of membrane proteins, including cryo-electron microscopy,^5–7^ nuclear magnetic resonance (NMR) spectroscopy,^8^ and single-molecule fluorescence^9–13^.

While nanodiscs are thought to provide a more physiologically relevant environment for membrane proteins than detergent micelles, increasing evidence indicates that nanodiscs can exert unintended influences on conformations and activities of embedded proteins. Cryo-EM structure determination of a ligand-gated ion channel demonstrated that the determined structure depended on the size of nanodisc used for reconstitution,^14^ and these structures differed from those determined in liposomes.^15^ Cryo-EM structures of an ABC transporter further showed that the protein could access a full conformational ensemble only in comparatively larger nanodiscs.^16^ Additional examples of nanodisc-dependent structural differences have been reported, including for the nicotinic acetylcholine receptor^17,18^ and GABA_A_ receptor.^19,20^ These observations have been largely attributed to direct noncovalent interactions between the scaffold proteins used for nanodisc assembly and the embedded membrane protein. This study focuses on an important yet less explored mechanism by which nanodiscs can modulate membrane protein structure and activity, specifically how lipid accessibility and organization within nanodiscs influences the conformational equilibria and biochemical activity of membrane proteins.

The conformational equilibria and signaling activity of G protein-coupled receptors (GPCRs) are well documented to be sensitive to the surrounding lipid environment.^21–24^ Lipid-dependent modulation of GPCR structure and function has been observed across an increasing number of receptors in both experimental^25–28^ and computational studies.^29–31^ The human A_2A_ adenosine receptor (A_2A_AR), a class A GPCR and important drug target for neurological diseases^32,33^ and potentially cancers^34,35^, has served as a model system for investigating GPCR signaling mechanisms^36–39^. In a previous study of A_2A_AR, NMR spectroscopic data, molecular simulations, and activity measurements demonstrated that receptor conformational equilibria were strongly influenced by the presence of anionic lipids, including POPS and POPG.^40^ In the present study, we leverage the sensitivity of A_2A_AR conformational equilibria to anionic lipids as an internal sensor to assess the accessibility of lipids to interact with the receptor in lipid nanodiscs.

Taking advantage of the precise control of lipid composition afforded by nanodiscs, we used ^19^F-NMR spectroscopy to systematically survey the conformational equilibria of A_2A_AR across binary mixtures of zwitterionic and anionic lipids at defined ratios in MSP nanodsics of varying sizes. We observe that a threshold concentration of anionic lipids is required to populate active A_2A_AR conformational ensembles, and that this threshold increases with increasing nanodisc size. NMR observations correlate with measurements of A_2A_AR-catalyzed Gα_S_ nucleotide exchange measured in nanodiscs of different sizes. Computational simulations, molecular modeling, and targeted mutagenesis of the scaffold proteins provided mechanistic insights into these observations and identified two major contributing factors: interactions between the scaffold protein and surrounding anionic lipids, and the formation of lipid nanoclusters within nanodiscs. Taken together, these findings indicate that the spatial organization and accessibility of lipids within nanodiscs can directly influence membrane protein conformational equilibria and receptor activity, and further suggest that these properties may be tunable through nanodisc engineering. Because membrane scaffold proteins share an identical sequence with the central region of Apo-A1, these findings may also have broader implications for lipid organization and protein-lipid interactions in larger, native lipid-protein assemblies.

## Results

### A_2A_AR can access its full conformational ensemble across nanodiscs of varying sizes

We first sought to determine whether A_2A_AR could access its full conformational ensemble in lipid nanodiscs of different sizes. We investigated human A_2A_AR in three different lipid nanodisc systems formed with the membrane scaffold proteins MSP1D1βH5,^4^ MSP1D1^1^ and MSP1E3D1, each containing defined binary mixtures of different molar ratios of the zwitterionic phospholipid POPC (1-palmitoyl-2-oleoyl-glycero-3-phosphocholine) and anionic lipids POPS (1-palmitoyl-2-oleoyl-sn-glycero-3-phospho-L-serine) or POPG (1-palmitoyl-2-oleoyl-sn-glycero-3-phospho-(1’-rac-glycerol)). These three membrane scaffold proteins were selected because they are widely used in biophysical and structural biology studies and because each produces nanodiscs of different average sizes, spanning from approximately 8 nm to 12 nm in diameter (Fig. 1). Samples of nanodiscs assembled from the three different scaffold proteins were highly pure and monodisperse, both without A_2A_AR and containing A_2A_AR (Supplementary Fig. 1 and Supplementary Fig. 2). The phospholipid composition of nanodisc samples was verified by ^31^P-NMR spectroscopy (Supplementary Fig. 3). Because the lipid headgroups exhibited unique chemical shifts, we quantified the relative amounts of lipids by integrating the relative signal intensities for each lipid species. This analysis showed that the relative amounts of each phospholipid species used in samples with defined binary mixtures quantitatively agreed with the intended molar ratios. Radioligand competition binding experiments confirmed the pharmacological activity of A_2A_AR[A289C] reconstituted into each of the three nanodisc systems (Supplementary Fig. 4), with determined K_i_ values for the agonist NECA that were within a factor of ∼3 for all systems and consistent with earlier measurements of A_2A_AR pharmacological activity in MSP1D1 nanodiscs^40^, with no apparent trend between nanodisc size and K_I_ value. This indicated that reconstituted A_2A_AR retained pharmacological activity in all three nanodisc systems.

**Fig. 1.**
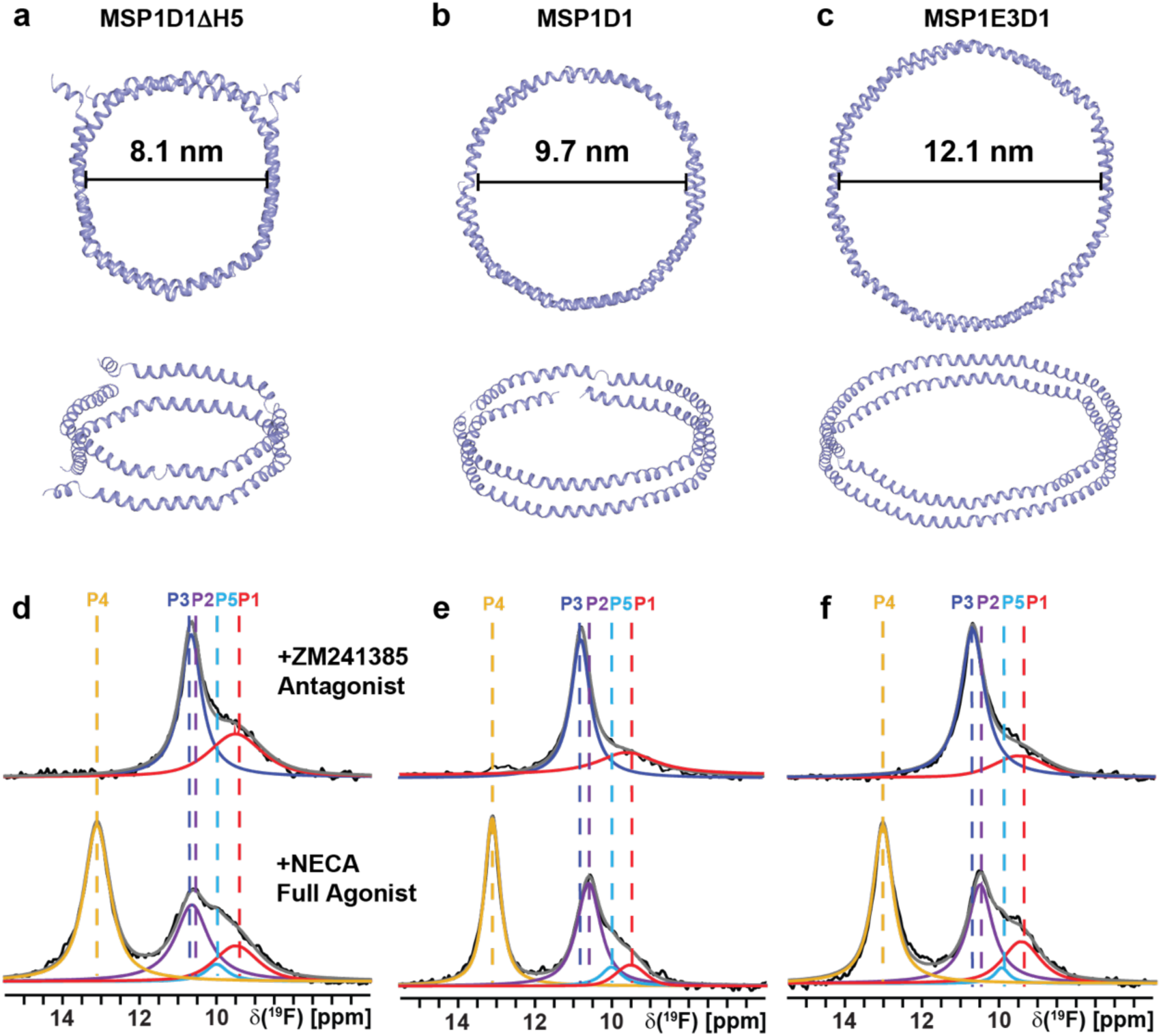
NMR-observed conformational equilibria of A_2A_AR compared across lipid nanodiscs of three distinct sizes. a–c. Models of the three different scaffold proteins taken from the initial configurations in simulations presented from top-down and side-on views for (a) MSP1D1βH5, (b) MSP1D1, and (c) MSP1E3D1. Lipids are not shown. The solid black lines show estimates of the diameter across each nanodisc at the initial simulation time. **d–f** The 1-dimensional ^19^F-NMR spectra of A_2A_AR[A289C^TET^] in complex with the antagonist ZM241385 or full agonist NECA reconstituted into nanodiscs containing the lipids POPC and POPS in a 70:30 molar ratio for nanodiscs composed of three different belt proteins. The NMR spectra shown as black lines are interpreted by Lorenztian deconvolutions with the minimal number of components that provided a good fit, labeled P1 to P5. The chemical shifts of P1 to P5 are indicated by the colored dashed vertical lines.

For ^19^F-NMR experiments, we expressed the human A_2A_AR containing a single extrinsic cysteine introduced into position 289, located near the intracellular-facing surface of transmembrane (TM) helix VII, producing A_2A_AR[A289C]. The resulting receptor retained pharmacological activity consistent with A_2A_AR expressed in mammalian cells, as previously validated.^9,38^ A ^19^F-NMR reporter group was introduced by reacting the solvent-accessible cysteine at position 289 with 2,2,2-trifluoroethanethiol (TET) using an in-membrane chemical modification approach^41^, yielding A_2A_AR[A289C^TET^]. Earlier studies employing both the same stable-isotope labeling methodology and A_2A_AR variant demonstrated no other cysteines were available for ^19^F labeling^38^. The ^19^F-NMR probe at position 289C was highly sensitive to both function-related changes in the efficacies of bound drugs and to the surrounding lipid bilayer environment in nanodiscs, providing ‘fingerprints’ for the corresponding functional states^38^.

We recorded 1-dimensional ^19^F-NMR spectra of A_2A_AR[A289C^TET^] for both complexes with antagonists and agonists in nanodiscs formed from the membrane scaffold proteins MSP1D1βH5, MSP1D1, and MSP1E3D1, each containing the same defined binary mixture of POPC and POPS at a molar ratio of 70 to 30. The ^19^F-NMR spectra of A_2A_AR[A289C^TET^] antagonist and agonist complexes in MSP1D1 nanodiscs were consistent with previously published data measured in nanodiscs formed from the same scaffold protein (Fig. 1 and Supplementary Fig. 5).^40,42^ For complexes with antagonists, ^19^F-NMR spectra manifested two components at δ ≈ 11.3 ppm (P3) and δ ≈ 9.5 ppm (P1) with the signal P3 being the largest signal in the spectrum (Fig. 1). ^19^F-NMR spectra for complexes with agonists contained a signal at the chemical shift of P1 and two new signals, P2 and P4, at δ ≈ 10.7 ppm and δ ≈ 13.1 ppm, respectively, and no signal intensity at the chemical shift for P3 (Fig. 1). State P4 was assigned previously to a fully active A_2A_AR conformation by experiments in which we observed a population of P4 upon complex formation of A_2A_AR[A289C^TET^] with mini-Gα_S_^40^, an engineered G protein.^43^

We also recorded 1-dimensional ^19^F-NMR spectra of A_2A_AR[A289C^TET^] for both complexes with antagonists and agonists in nanodiscs formed from the scaffold proteins MSP1D1βH5 (Fig. 1d) and MSP1E3D1 (Fig. 1f). ^19^F-NMR spectra of A_2A_AR[A289C^TET^] in nanodiscs formed from MSP1E3D1 were nearly identical to those observed with nanodiscs formed from MSP1D1 for both antagonist and agonist complexes, exhibiting signals with identical chemical shifts and at nearly the same relative peak intensities, with only marginal differences in observed line widths (Fig. 1f). NMR spectra of A_2A_AR[A289C^TET^] in nanodiscs formed from MSP1D1βH5 were also highly similar to spectra in nanodiscs formed from MSP1D1, with nearly identical chemical shifts and relative peak intensities and slightly increased line broadening for component P4 (Fig. 1d). The highly similar chemical shifts for all three nanodisc systems suggests that there is not a direct influence on the probe environment by the different scaffold proteins. Most importantly, comparison of these data indicate that the A_2A_AR function-related conformational equilibrium is largely preserved across all three nanodisc systems for complexes with antagonists and agonists.

## A threshold of POPS is required to populate the fully active A_2A_AR conformation in nanodiscs

We previously demonstrated that ^19^F-NMR spectra measured for agonist-bound A_2A_AR[A289C^TET^] in MSP1D1 nanodiscs without anionic lipids closely resembled spectra for antagonist-bound A_2A_AR, most notably by the absence of the P4 resonance assigned to a fully active conformation of A_2A_AR.^40^ Given that the receptor remained pharmacologically active and bound both antagonists and agonists with native affinity in this condition^40^, the NMR data were interpreted as evidence that activation of agonist-bound A_2A_AR required association with anionic lipids to populate a fully active conformation. This conclusion was further supported by mutagenesis experiments, additional NMR data and molecular simulations that identified direct interactions between anionic lipid headgroups and a region at the intracellular surface of A_2A_AR between TM VI and TMVII involved in complex formation with G proteins.^40^ Collectively, these data demonstrated that the conformational equilibrium of A_2A_AR was very sensitive to interactions with anionic lipids in the surrounding membrane environment. We therefore reasoned that the conformational equilibrium of A_2A_AR would serve as a sensitive probe of lipid accessibility within nanodiscs. Specifically, when agonist-bound A_2A_AR has access to anionic lipids, we expect to observe an active conformational ensemble, whereas if A_2A_AR is unable to access anionic lipids we would expect to observe a conformational ensemble that more closely resembles that observed for an inactive receptor. We could then leverage the clearcut differences in the ^19^F-NMR spectra between inactive and active conformational ensembles to detect the receptor’s access to anionic lipids in nanodiscs.

We leveraged the ability to precisely control lipid composition in nanodiscs to systematically compare the ^19^F-NMR observed conformational equilibria of A_2A_AR in complex with the full agonist NECA across a range of defined ratios of zwitterionic and anionic lipids for nanodiscs prepared from all three membrane scaffold proteins. ^19^F-NMR spectra of the A_2A_AR–NECA complex measured in MSP1D1 nanodiscs containing a 70:30 molar ratio of POPC and POPS showed a conformational equilibria consistent with previous reports,^40,44^ including observation of the component P4 assigned to the fully active conformation (Fig. 1, Fig. 2 and Supplementary Fig. 6). At a 78:22 molar ratio of POPC and POPS, we observed a nearly identical conformational equilibrium for agonist-bound A_2A_AR in MSP1D1 nanodiscs (Fig. 2b). For samples containing less POPS than the 78 to 22 molar ratio, we observed a clear loss of the component P4, and these spectra appeared highly similar to ^19^F-NMR data of agonist-bound A_2A_AR measured in nanodiscs completely lacking anionic lipids (Fig. 2b, top row). These observations suggest that below this threshold ratio, agonist-bound A_2A_AR no longer has sufficient access to anionic lipids to populate the fully active conformational ensemble.

**Fig. 2.**
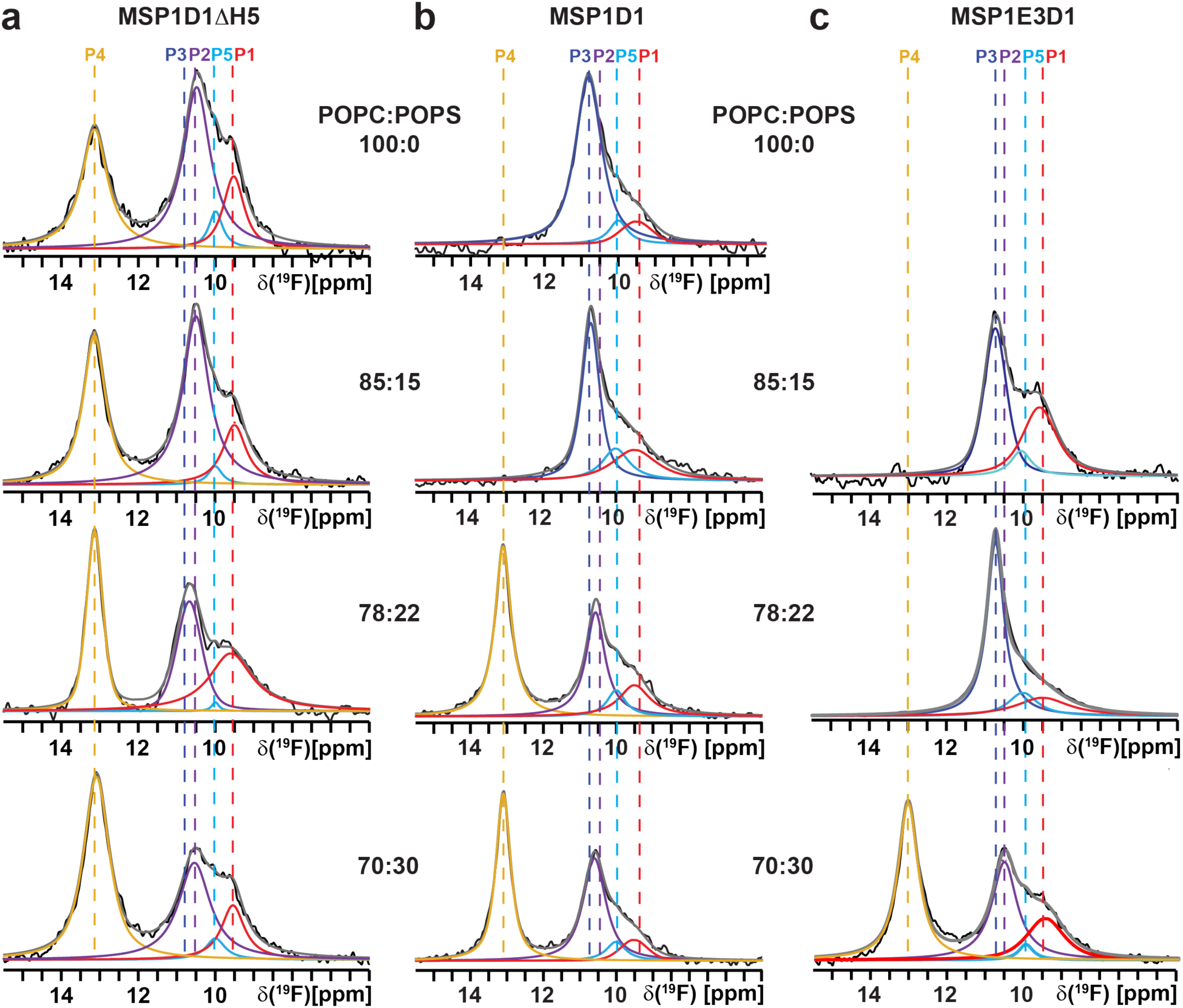
Conformational response of agonist-bound A_2A_AR in three different nanodisc sizes with varying amounts of the anionic lipid POPS. The 1D ^19^F-NMR spectra of A_2A_AR[A289C^TET^] in complex with the full agonist NECA in nanodiscs containing different defined ratios of POPC and POPS composed of the scaffold protein (a) MSP1D1βH5, (b) MSP1D1, and (c) MSP1E3D1. Other figure presentation details are the same as in Fig. 1.

^19^F-NMR spectra of the A_2A_AR–NECA complex measured in MSP1E3D1 nanodiscs, the largest nanodiscs studied, containing a 70:30 molar ratio of POPC:POPS showed a conformational equilibria very similar to that observed for MSP1D1 nanodiscs containing the same lipid composition, including the presence of component P4 (Fig. 1 and Fig. 2). We were also able to prepare samples of the A_2A_AR–NECA complex in MSP1E3D1 nanodiscs with POPC:POPS lipid molar ratios of 78:22 and 85:15. Multiple attempts were made to prepare samples of A_2A_AR in MSP1E3D1 nanodiscs containing only POPC lipids, which resulted in unstable and heterogeneous samples that were excluded from NMR analysis. These additional samples established that in MSP1E3D1 nanodiscs, component P4 was completely absent in nanodiscs below a 70:30 molar ratio of POPC:POPS lipids, yielding spectra that resembled those of agonist-bound A_2A_AR in nanodiscs lacking anionic lipids (Fig. 2c). These data indicate that below this threshold, agonist-bound A_2A_AR is unable to access anionic lipids to populate a fully active conformational ensemble. This threshold was notably higher than that observed for MSP1D1 nanodiscs, and thus the amount of POPS required to populate the fully active conformational ensemble increased with the size of the nanodisc system.

We next examined the conformational equilibrium of agonist-bound A_2A_AR in the smallest nanodiscs formed from the membrane scaffold protein MSP1D1βH5. In contrast to both MSP1D1 and MSP1E3D1 nanodisc systems, agonist-bound A_2A_AR in MSP1D1βH5 nanodiscs populated active conformational ensembles across all tested ratios of POPC and POPS (Fig. 2). Remarkably, active conformational ensembles were also observed in MSP1D1βH5 nanodiscs containing no anionic lipids. This behavior substantially differed from that observed in MSP1D1 and MSP1E3D1 nanodiscs, as reported previously^40,44^ and in the present study. Considered together with the additional measurements of A_2A_AR-catalyzed nucleotide exchange in MSP1D1 nanodiscs (see below), these observations suggest that the conformational equilibrium of A_2A_AR is influenced by properties of MSP1D1βH5 nanodiscs, either indirectly through differences in lipid-protein behavior or potentially directly through interactions between the receptor and scaffold protein (see Discussion).

## POPG populates active A_2A_AR conformational ensembles at lower threshold concentrations than POPS

To determine whether the threshold concentration of anionic lipids required to populate an active A_2A_AR conformational ensemble was specific to POPS or extended to other anionic lipids, we systematically compared the ^19^F-observed conformational ensemble of agonist-bound A_2A_AR in the three nanodisc systems containing defined binary mixtures of POPC with the anionic lipid 1-palmitoyl-2-oleoyl-sn-glycero-3-phospho-(1’-rac-glycerol) (POPG). For MSP1D1 nanodiscs prepared with a 70:30 molar ratio of POPC:POPG, the ^19^F-NMR spectra of the A_2A_AR-NECA complex were very similar to those measured with the same ratio of POPC and POPS in MSP1D1 nanodiscs, including the same chemical shift values for all components in the spectrum and similar peak intensities (Fig. 3 and Supplementary Fig. 7). This observation indicated that the A_2A_AR active conformational ensemble was not significantly influenced by the specific chemical structure of the anionic lipid headgroup, consistent with previous NMR measurements in additional binary lipid mixtures.^40^ For spectra measured with a lower concentration of POPG at a molar ratio of 85:15 POPC:POPG, we observed an overall active conformational ensemble for agonist-bound A_2A_AR (Fig. 3b). This contrasts with NMR measurements with MSP1D1 nanodiscs containing a POPC:POPS molar ratio of 85:15, which showed a conformational equilibrium resembling an inactive conformational ensemble for agonist-bound A_2A_AR (Fig. 2b). This indicated that in the same nanodiscs and at the same molar ratio of anionic lipids, POPG remained sufficiently accessible to interact with A_2A_AR and populate an active conformational ensemble, in contrast to POPS at the same molar ratio.

**Fig. 3.**
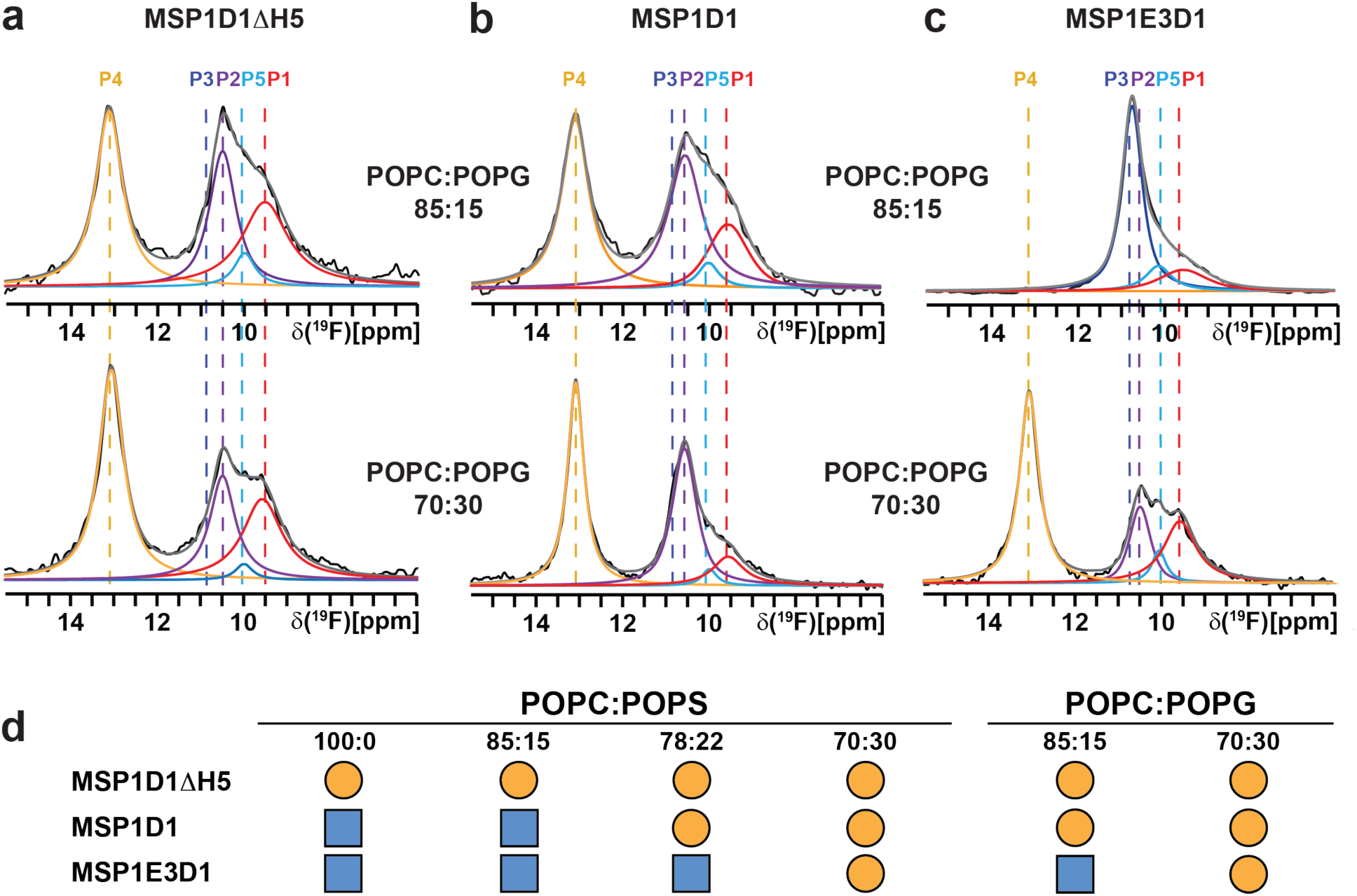
Comparison of the conformational equilibria of agonist-bound A_2A_AR in nanodiscs of varying sizes containing defined mixtures of POPC and POPG and summary of NMR data. a–c. The 1D ^19^F-NMR spectra of A_2A_AR[A289C^TET^] in complex with the full agonist NECA in nanodiscs containing different defined ratios of POPC and POPG composed of the scaffold protein **a** MSP1D1βH5, **b** MSP1D1, and **c** MSP1E3D1. Other figure presentation details are the same as in Fig. 1. **d** Summary of ^19^F-observed conformational equilibria of A_2A_AR[A289C^TET^] in the three nanodisc systems studied across different defined binary lipid mixtures. Orange circles indicate conditions in which an active conformational ensemble is observed for agonist-bound A_2A_AR, and blue squares indicate conditions in which an active conformational ensemble is not observed for agonist-bound A_2A_AR.

We next compared the conformational equilibria of agonist-bound A_2A_AR in MSP1E3D1 and MSP1D1βH5 nanodiscs containing defined binary mixtures of POPC and POPG (Fig. 3a and 3c). For agonist-bound A_2A_AR reconstituted into MSP1E3D1 nanodiscs containing a 70:30 molar ratio of POPC:POPG, the observed conformational equilibrium was very similar to that measured for MSP1D1 nanodiscs containing the same lipid composition. In contrast, agonist-bound A_2A_AR reconstituted into MSP1E3D1 nanodiscs containing a POPC:POPG molar ratio of 85:15 exhibited a conformational equilibrium resembling inactive A_2A_AR, including loss of the P4 resonance assigned to the fully active state (Fig. 3c). This indicated that we also observed a threshold of POPG required to populate an active conformational ensemble for A_2A_AR, though this threshold was lower than that observed for POPS. For agonist-bound A_2A_AR reconstituted into MSP1D1βH5 nanodiscs, active conformational ensembles were observed across all tested ratios of POPC and POPG (Fig. 3a).

## Nucleotide exchange measurements validate NMR observations of nanodisc-dependent A_2A_AR activation

We investigated the impact of nanodisc size and lipid composition on A_2A_AR[A289C] activity by measuring A_2A_AR-catalyzed nucleotide exchange in the three nanodisc systems containing defined binary lipid mixtures using a GTP hydrolysis assay^45^ (Fig. 4). This assay monitors the exchange of GTP for GDP bound to the stimulatory G protein Gα_S_ following complex formation with A_2A_AR[A289C]. We first compared GTPase activity for A_2A_AR[A289C] reconstituted in nanodiscs formed from the membrane scaffold proteins MSP1D1βH5, MSP1D1, and MSP1E3D1, each prepared with the same molar ratio of POPC and POPS of 78:22 (Fig. 4a). This lipid composition was selected based on the ^19^F-NMR observations that showed significant differences in the relative populations of active A_2A_AR between MSP1D1 and MSP1E3D1 nanodiscs (Fig. 2).

**Fig. 4.**
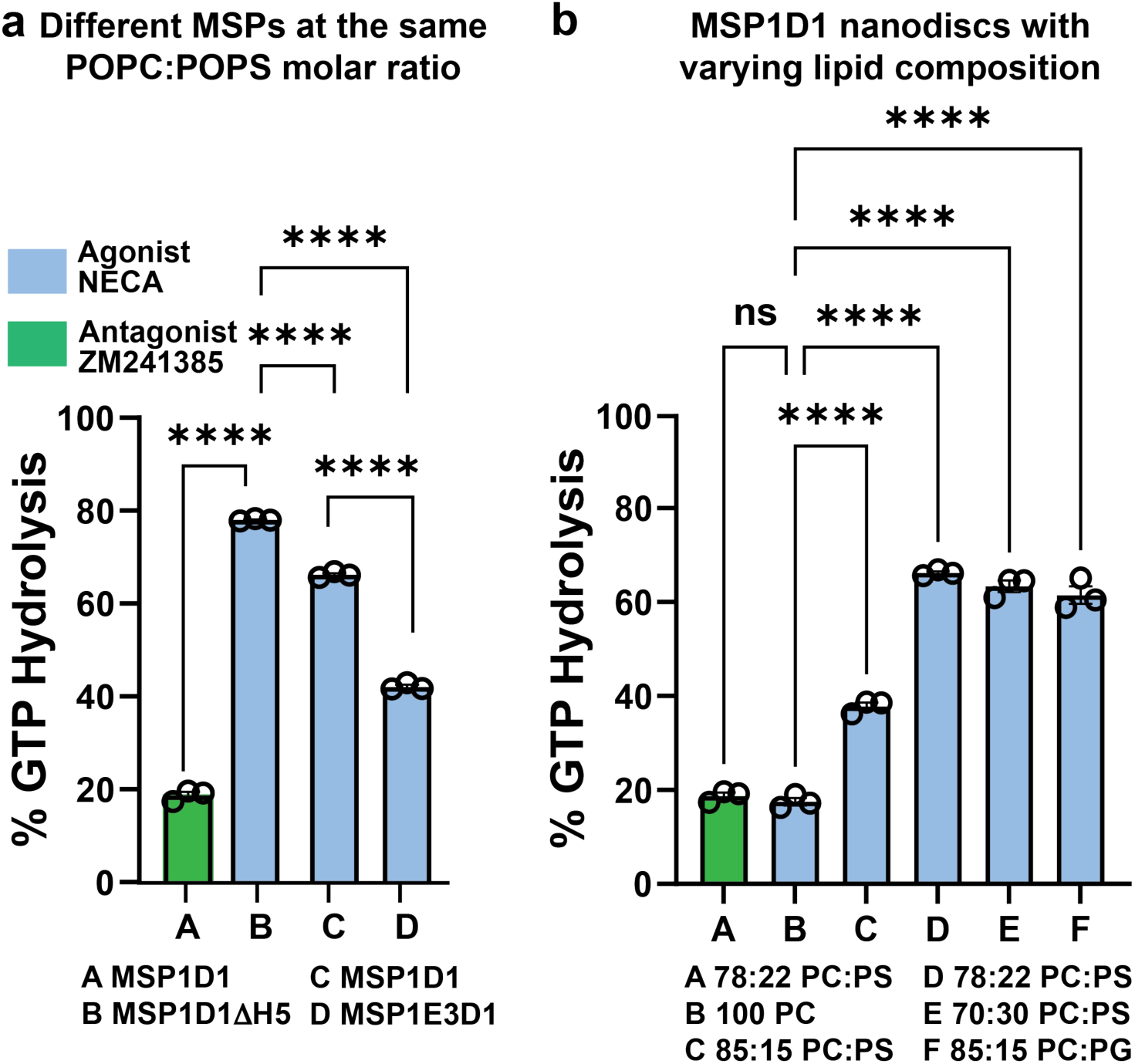
A_2A_AR-catalyzed GTP hydrolysis in different nanodisc systems. (a and. **b)** The percent GTP hydrolysis by Gα_S_ measured using a GTPase-Glo assay in the presence of A_2A_AR complexes with ligands reconstituted in different nanodisc systems. **a** Percent GTP hydrolysis measured with A_2A_AR reconstituted into nanodiscs formed from MSP1D1, MSP1βH5, and MSP1E3D1, each containing POPC and POPS at a molar ratio of 78:22. The green bar indicates measurements for A_2A_AR in complex with the antagonist ZM241385, included as an internal control, and blue bars indicate measurements made with A_2A_AR in complex with the full agonist NECA. **b** Percent GTP hydrolysis measured with A_2A_AR in MSP1D1 nanodiscs containing different molar ratios of POPC and POPS or with a binary mixture of POPC and POPG, as indicated in the plot. The color scheme is the same as in **a**.

At this lipid ratio, A_2A_AR reconstituted in MSP1D1βH5 nanodiscs showed the highest level of GTPase activity (Fig. 4a), qualitatively consistent with observations of a relatively high population of the active conformation of A_2A_AR in ^19^F-NMR data. Relative to this condition, A_2A_AR in MSP1D1 nanodisc showed a ∼15% decrease in GTPase activity, although activity remained substantially greater than that observed for the antagonist-bound control sample of A_2A_AR in complex with ZM241385 (Fig. 4a). In contrast, A_2A_AR[A289C] reconstituted in MSP1E3D1 nanodiscs showed a marked decrease in nucleotide exchange activity at the same lipid ratio (Fig. 4a). This observation was qualitatively consistent with the ^19^F-NMR measurements, which showed a significant reduction in the active-state conformational population for A_2A_AR in MSP1E3D1 nanodiscs containing the same lipid composition (Fig. 2).

We next compared GTPase activity for A_2A_AR[A289C] reconstituted in MSP1D1 nanodiscs across a range of increasing POPS concentrations and in MSP1D1 nanodiscs containing a binary mixture of POPC and POPG. For A_2A_AR in MSP1D1 nanodiscs composed entirely from POPC, GTPase activity was comparable to the low level of activity observed for the control sample of A_2A_AR bound to the antagonist ZM241385 (Fig. 4b), consistent with ^19^F-NMR observations (Fig. 1 and Fig. 2). Increasing the proportion of anionic lipid to a POPC:POPS molar ratio of 85:15 resulted in increased GTPase activity relative to samples prepared with only POPC. This level of activity was comparable to the somewhat reduced signaling activity observed for A_2A_AR in MSP1E3D1 nanodiscs containing a ratio of POPC:POPS molar ratio of 78 to 22 (Fig. 4, a and b). Further increasing the proportion of POPS led to additional increases in GTPase activity up to a POPC:POPS ratio of 78:22, beyond which no further significant increases in activity were observed (Fig. 4b). A_2A_AR reconstituted in MSP1D1 nanodiscs containing POPC and POPG at a molar ratio of 85:15 showed significantly higher GTPase activity as compared to nanodiscs containing POPC and POPS at the same molar ratio (Fig. 4b). This observation was consistent with the ^19^F-NMR observations, which showed a significantly higher population of the active A_2A_AR conformation in MSP1D1 nanodiscs containing POPG relative to POPS at the same molar ratio (Fig. 3).

### Anionic lipid clustering reduces effective lipid availability to A_2A_AR in nanodiscs

^19^F-NMR measurements of the conformational equilibria of agonist-bound A_2A_AR in the three nanodisc systems studied showed that the amount of anionic lipids required to populate an active A_2A_AR conformational ensemble increased with increasing nanodisc size (Fig. 2, Fig. 3d). Measurements of nucleotide exchange activity showed the same trend (Fig. 4). These effects were substantially more pronounced for POPS than for POPG. These observations were unexpected when considered in the context of geometric properties of nanodiscs. Approximating nanodiscs as circular particles, the number of lipids scales approximately with the square of the nanodisc radius, whereas the nanodisc circumference increases linearly with the radius. Consequently, if interactions between the scaffold protein and lipids at the nanodisc perimeter were the primary mechanism responsible for sequestering anionic lipids from A_2A_AR, the relative influence of these interactions would be expected to decrease with increasing nanodisc size. Experimentally, however, we observe the opposite trend.

To explain these observations, we hypothesized that interactions scaling with the total number of lipids within nanodiscs underlie the observed behavior. Previous solid-state NMR studies of phosphatidylserine-containing membranes showed that calcium can induce the formation of distinct PS lipid populations^46^, including populations assigned to PS–PS interactions mediated by calcium ions. Simulations have obtained similar results for other anionic lipids, particularly inositol-containing lipids^47^ (see Discussion). Because PS headgroups can participate in intermolecular hydrogen-bonding interactions, whereas PG headgroups cannot, we considered whether POPS molecules could form transient hydrogen-bonded nanoclusters within nanodiscs, reducing the effective chemical activity and accessibility of POPS, thereby limiting its availability to interact with A_2A_AR as compared with POPG.

To test this hypothesis, we performed molecular simulations of all three nanodisc systems containing mixtures of either POPC and POPS or POPC and POPG lipids at molar ratios of 80:20 POPC:anionic lipid. Simulations were also performed for the two largest nanodisc systems containing A_2A_AR and the same lipid compositions (see Supplementary Table 1 for simulation conditions and durations). A stable initial configuration of the smallest nanodiscs formed from MSP1D1βH5 containing A_2A_AR could not be obtained for production simulations and was therefore excluded from further analysis.

The final 4.5 μs of each 5 μs long trajectory were analyzed to quantify the frequency of pairwise interactions between anionic lipid headgroups and the formation of higher-order lipid clusters (see Methods). POPS headgroups participated substantially more frequently than POPG headgroups in both pairwise interactions and in the formation of higher-order lipid clusters (Fig. 5a and Fig. 5b). This observation was consistent for nanodisc systems both without and with A_2A_AR. Additionally, the fraction of POPS molecules that participated in clusters, particularly clusters containing three or four lipid molecules, increased with nanodisc size (Fig. 5a and Fig. 5b). This trend was also consistent for nanodisc systems both without A_2A_AR and containing A_2A_AR. Observations from the simulations indicate that POPS clustering correlates with an increased threshold of POPS required to populate the active A_2A_AR conformational ensemble, suggesting that POPS lipids in clusters are not available to directly interact with A_2A_AR. Because Na^+^ was present in the NMR samples and present in simulations, we investigated whether Na^+^ contributed to the formation of POPS clusters. Simulations showed that Na^+^ did not contribute significantly to cluster formation (Supplementary Fig. 8).

**Fig. 5.**
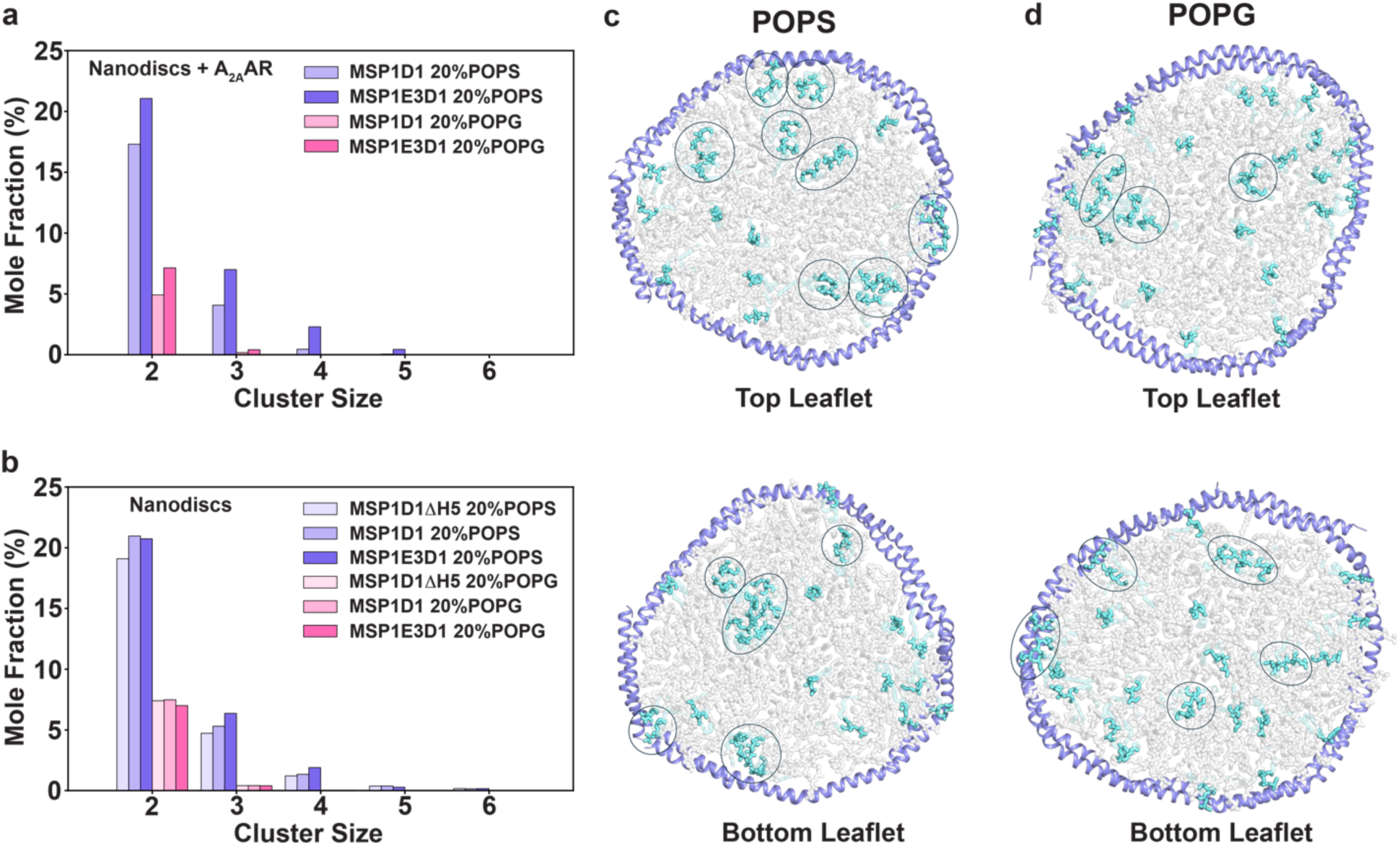
Clustering of POPS and POPG lipids within nanodiscs. a and. **b** Mole fractions of POPS or POPG lipids in nanoclusters of varying sizes for (a) nanodiscs formed from two different scaffold proteins containing lipids and A_2A_AR or (b) nanodiscs formed from three different scaffold proteins containing defined binary lipid mixtures. MSP1D1βH5 was not studied with A_2A_AR embedded, as that system could not be equilibrated in simulations (see text). **c and d** Representative snapshots from MD simulations with nanodiscs formed from the scaffold protein MSP1E3D1 with POPC and (c) POPS or (d) POPG lipids. A nanocluster is defined as the formation of at least two POPS or POPG lipids in close proximity, indicated by circles. The snapshots were taken from the frames exhibiting the highest mole fraction of (c) POPS or (d) POPG in nanoclusters.

We then analyzed the hydrogen-bond dynamics from MD simulations (See Methods). The hydrogen-bond lifetimes of anionic lipids were calculated to be 0.45-0.70 ps for POPS and 0.09-0.13 ps for POPG in the simulated nanodisc systems (Supplementary Fig. 9 and Supplementary Table 2). Although these lifetimes were relatively short overall, the markedly longer lifetime observed for POPS hydrogen bonds indicates a greater persistence of POPS pairs/clusters compared with POPG.

## Basic residues in membrane scaffold proteins influence anionic lipid organization in nanodiscs

Previous biophysical studies have reported that anionic lipid molecules within nanodiscs could interact at the nanodisc periphery with positively charged residues of membrane scaffold proteins^48^. Thus, although the observed clustering of POPS correlated with the NMR data and nucleotide exchange measurements, we sought to determine whether interactions between anionic lipid headgroups and side chains of basic residues of MSPs could additionally contribute to limiting the availability of anionic lipids for interacting with A_2A_AR embedded in nanodiscs. For each positively charged scaffold protein side chain, we quantified the fraction of simulation time during which the residue interacted with an anionic lipid headgroup. These interactions were then classified into three categories based on interaction frequency: strong (β40% of simulation time), medium (β5%), and weak (< 5%).

Across all three nanodisc systems, we observed strong, medium, and weak interactions between basic MSP residues and POPS or POPG headgroups in both nanodiscs without A_2A_AR and nanodiscs containing A_2A_AR (Fig. 6a). Although the total mole fraction of strong, medium, and weak interactions remained about the same for all systems, the relative distribution of these interactions differed substantially. We observed a significantly larger mole fraction of strong interactions for all three nanodisc systems containing POPS than for nanodiscs containing POPG, both for nanodiscs without A_2A_AR and nanodiscs containing A_2A_AR (Fig. 6a). Comparison across nanodisc sizes revealed a clear increase in the mole fraction of strong POPS interactions with increasing nanodisc diameter, observed for both nanodiscs without A_2A_AR and nanodiscs containing A_2A_AR.

**Fig. 6.**
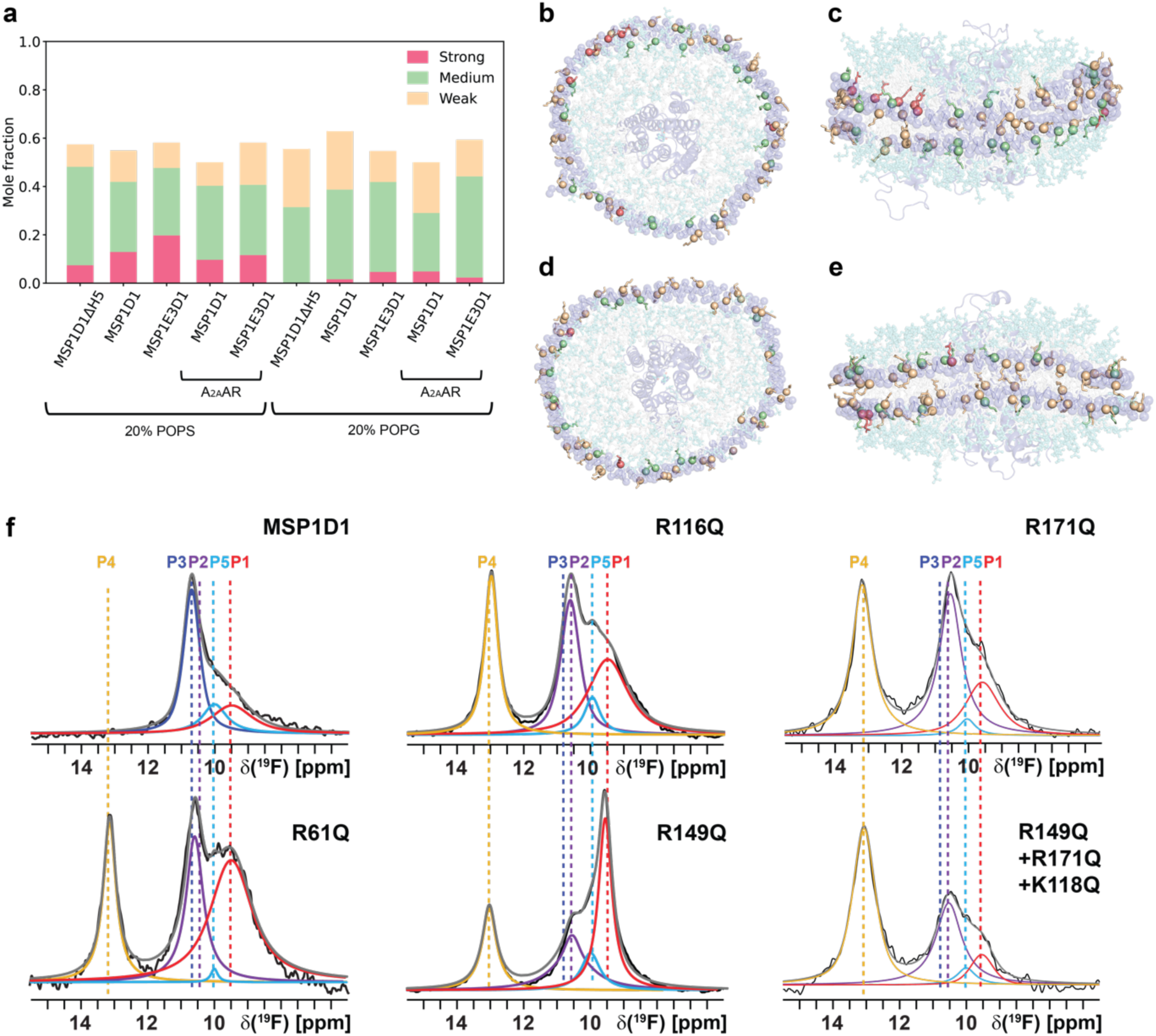
Interactions of POPS and POPG lipids with positively charged residues Arg/Lys in nanodiscs. **a** Mole fractions of positively charged residues in nanodiscs interacting with POPS/POPG lipids at three interaction levels (<5%, ≥5%, and ≥40% of simulation time). The annotation “A_2A_AR” indicates nanodisc systems containing embedded A_2A_AR. The annotations “20% POPS” and “20% POPG” denote the two membrane compositions, POPS:POPC (1:4) and POPG:POPC (1:4), respectively. **b** and **c** Top and side views of MSP1D1 containing A_2A_AR embedded in a POPS:POPC (1:4) membrane. **d** and **e** Top and side views of MSP1D1 containing A_2A_AR embedded in a POPG:POPC (1:4) membrane. The Cα atoms of MSP1D1 residues are shown as spheres: red indicates residues exhibiting strong interactions with POPS/POPG lipids (≥40% of the simulation time), green indicates medium interactions (≥5%), and wheat indicates weak interactions (<5%), including zero interactions for Arg/Lys residues that do not contact lipids. Definitions of strong, medium, and weak interactions are provided in the Methods section. Non-positively charged residues are shown in purple with transparency. Lipid headgroups are shown as aquamarine transparent sticks, while lipid tails are shown as gray transparent sticks. A_2A_AR is shown as a transparent purple cartoon representation, and the ligand is shown as transparent sticks. **f** 1D ^19^F-NMR spectra of A_2A_AR[A289C^TET^] in complex with the full agonist NECA in nanodiscs containing POPC and POPS at a molar ratio of 85:15, formed from MSP1D1 or variant MSP1D1 proteins containing one or more amino acid replacements, as annotated. Other figure presentation details are the same as in Fig. 1.

**Fig. 7.**
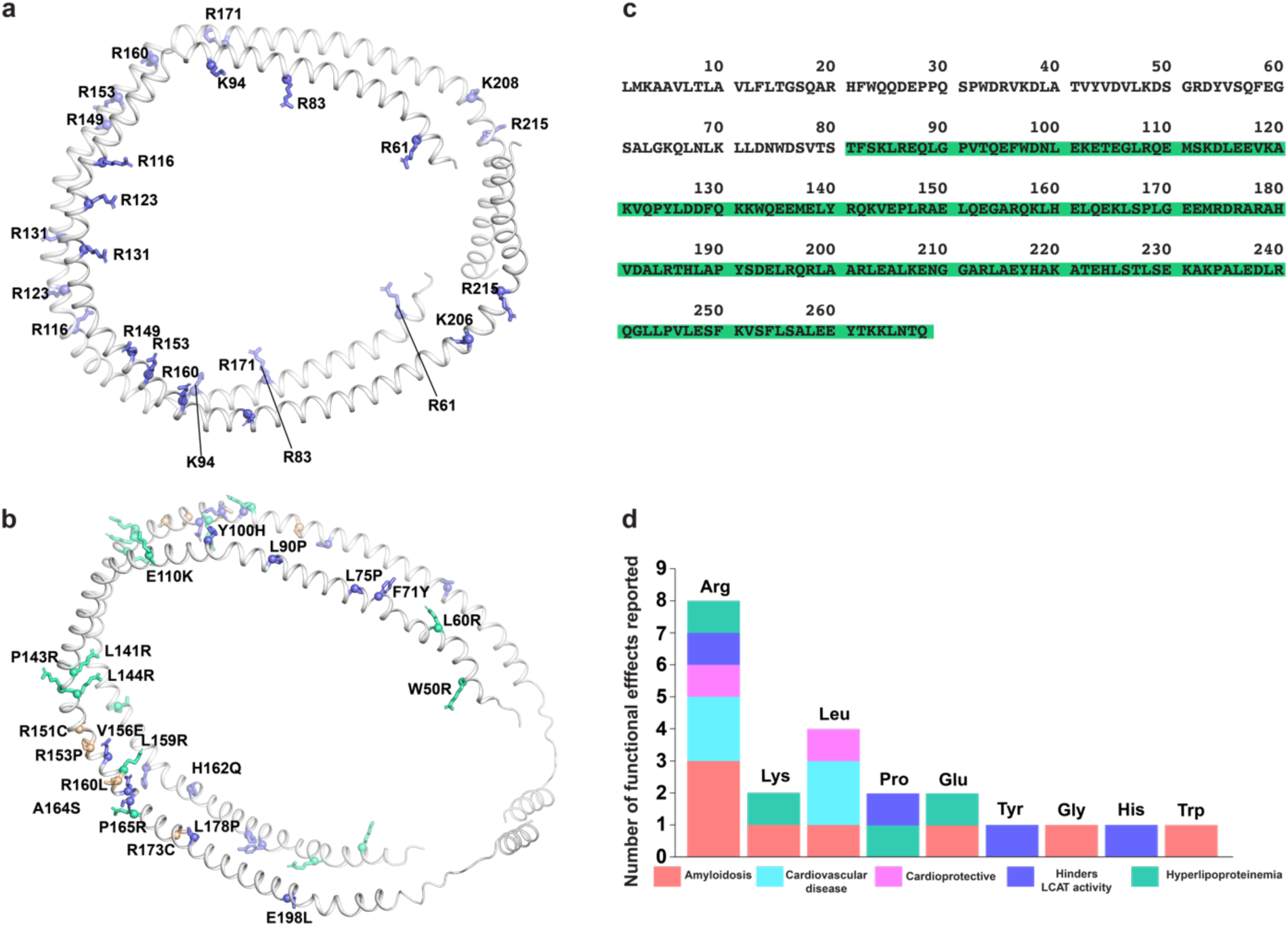
Comparison of membrane scaffold protein and Apolipoprotein A1 sequences and mapping of disease-associated Apo-A1 mutations. **a** Model of MSP1D1 with positively charged residues in colors and annotated on the model. **b** Model of Apo-A1 with positively charged residues associated with several diseases in colors and annotated on the model. **c** Sequence comparison of MSP1D1 and Apo-A1. The full amino acid sequence of Apo-A1 is shown, and residues from Apo-A1 that form MSP1D1 are highlighted with a green background. **d** Visualization of disease-associated single point mutations in Apo-A1 arranged by amino acid type. The height of each vertical bar represents the number of reported mutations associated with Apo-A1 functional changes, and each disease category is colored.

Nanodiscs containing POPG showed only a marginal increase in strong interactions with increasing nanodisc size, with a similar to slightly lower proportion of strong interactions in the largest nanodiscs containing A_2A_AR (Fig. 6a).

To visualize site-specific interactions between MSP basic residues and anionic lipids, we mapped those interactions onto models of MSP1D1 nanodiscs generated from molecular dynamics simulations of nanodiscs containing A_2A_AR and an 80:20 mole fraction of POPC and POPS or POPC and POPG (Fig. 6b-e). Interactions with other nanodisc systems are shown in Supplementary Figs. 10-11. The interaction frequences for every residue are listed in Supplementary Tables 3-16. Differences in the number of strongly interacting residues were clearly visible between nanodiscs containing a binary mixture of POPC and POPS over nanodiscs containing POPC and POPG. The side chains of basic residues involved in strong and medium interactions were oriented toward the lipid bilayer headgroups (Fig. 6). Conversely, those participating in weak interactions and failing to engage with lipids entirely were directed either toward the interface between the two MSP chains or outward into the solvent (Fig. 6).

The binding equilibrium between anionic lipids and A_2A_AR is perturbed by both lipid self-clustering and competing interactions with MSP basic residues. Together, these effects reduce the availability of receptor-bound anionic lipids and shift the equilibrium toward lipid release, thereby favoring the inactive conformational ensemble of the receptor. Accordingly, the H230^6.32^-R291^7.56^ distance distribution, serving as a reporter for distinguishing between the active and inactive conformations of A_2A_AR in prior work^40^, shifts toward smaller values in MSP1E3D1 nanodiscs containing a mixture of POPC:POPS (80:20 molar ratio) when compared to mixtures of POPC:POPG at the same molar ratio (Supplementary Fig. 14).

We next used ^19^F-NMR spectroscopy to test whether replacement of MSP1D1 basic residues identified in molecular simulations as interacting with anionic lipids impacted the conformational equilibrium of agonist-bound A_2A_AR. Individual amino acid substitutions were introduced into the MSP1D1 scaffold protein, and agonist-bound A_2A_AR[A289C^TET^] was reconstituted into variant MSP1D1 nanodiscs containing a POPC:POPS molar ratio of 85:15. This lipid composition was specifically selected because ^19^F-NMR spectra of agonist-bound A_2A_AR reconstituted in nanodiscs formed from the original scaffold protein exhibited an inactive conformational equilibrium in these conditions due to the lack of accessible POPS (Fig. 2).

We prepared samples of agonist-bound A_2A_AR[A289C^TET^] into MSP1D1 nanodiscs containing amino acid substitutions selected from the simulations that showed the greatest frequencies of interactions from simulations. These included the single substitutions MSP1D1[R61Q], MSP1D1[R116Q], MSP1D1[R149Q], and MSP1D1[R171Q], as well as several variants containing combinations of these substitutions. All MSP1D1 variants could be expressed and purified to homogeneity and showed similar levels of purity and size as assessed by SDS-PAGE and analytical SEC characterization (Supplementary Fig. 12). We next recorded ^19^F-NMR spectra of agonist-bound A_2A_AR[A289C^TET^] reconstituted into nanodiscs formed from the MSP1D1 variants with a POPC:POPS molar ratio of 85:15. For all variants tested, we observed a significant increase in the population P4 of the active state and decrease in the population P3 of the inactive state, and the resulting spectra resembled that obtained with the higher POPC:POPS lipid molar ratio of 70:30 (Fig. 6f and Supplementary Fig. 13). We did observe variation in the relative population of P4 among the nanodiscs formed from MSP1D1 variants. The largest difference was observed for nanodiscs formed from the variant R149Q, which showed a significant increase in the active state P4 population, though approximately half of the signal intensity as compared with the other MSP1D1 variants or with MSP1D1 and a higher POPC:POPS lipid molar ratio (Fig. 6f). These data suggest that the R149Q substitution partially increases the accessibility of POPS to A_2A_AR but is insufficient to fully restore access to levels observed under conditions supporting a fully active conformational ensemble. Consistent with this interpretation, combining R149Q with additional substitutions, including R171Q, resulted in conformational equilibria characterized by complete restoration of the active-state population and the appearance of a fully active conformational ensemble (Fig. 6f).

## Sequence conservation between MSPs and apolipoprotein A1 suggests shared principles of lipid organization in biological protein-lipid assemblies

Membrane scaffold proteins were originally developed from apolipoprotein A1 (Apo-A1)^1,2^, the primary structural protein of high-density lipoprotein (HDL) particles. Like nanodiscs, Apo-A1 consists of a series of amphipathic α-helices that form a belt-like structure surrounding lipid molecules^49^. Although a crystal structure of human Apo-A1 has been determined^49^, this structure was solved in the absence of lipids. To visualize the spatial distribution of basic residues in Apo-A1 and their location with respect to the contained lipids, we therefore generated a model of Apo-A1 containing an 80:20 molar ratio of POPC and POPS and compared this to our model of MSP1D1 containing the same defined lipid composition (Fig. 8a, b). MSP1D1 was generated by truncating the N-terminal region of Apo-A1, and thus all 160 amino acid residues of MSP1D1 are contained within a central region of Apo-A1 that is essential for the formation of the disc-like architecture (Fig. 8c).

We next surveyed the available literature on point mutations in Apo-A1 that are associated with various diseases, including cardiovascular diseases and amyloid disorders, as reviewed^50^, and categorized these mutations according to amino acid type (Fig. 8d). From 22 identified disease-associated point mutations in Apo-A1, nearly half of these mutations (10 of 22) involve either the replacement of a positively charged residue with a neutral residue or the replacement of a neutral residue with a positively charged residue (Fig. 8d). Although the exact relationship between many of these mutations and disease phenotypes are not well understood mechanistically, these observations raise the possibility that at least some mutations may alter interactions between Apo-A1 and lipids within HDL particles. More broadly, these findings suggest that modulation of lipid organization by protein-lipid interactions may represent a general principle extending beyond nanodiscs to larger biological lipid-protein assemblies.

## Discussion

The systematic ^19^F-NMR measurements of A_2A_AR conformational equilibria across the three nanodisc systems and multiple lipid compositions reveal a complex relationship between nanodisc size, lipid organization, and their collective influence on GPCR energy landscapes. These observations suggest that the question of which nanodisc system provides the most physiologically relevant environment for GPCRs does not have a simple or universal answer. One consistent trend that emerged from this systematic analysis was that increasing nanodisc size required progressively larger amounts of anionic lipids to populate an active A_2A_AR conformational ensemble for the agonist bound receptor. Thus, this suggests that increasing nanodisc size while maintaining the same relative molar ratio of lipids within nanodiscs may result in a deviation from receptor physiological activity.

In the smallest nanodiscs formed from MSP1D1βH5, the conformational equilibria observed for antagonist-bound A_2A_AR and for agonist-bound A_2A_AR in nanodiscs containing a molar ratio of 70:30 POPC:POPS were highly similar to those measured in the two larger nanodisc systems containing the same lipid composition (Fig. 1). However, unlike the two larger nanodisc systems, for agonist-bound A_2A_AR in MSP1D1βH5 nanodiscs, we consistently observed an active conformational ensemble across all lipid compositions tested, including conditions in which both ^19^F-NMR measurements and nucleotide exchange assays in the larger nanodisc systems showed substantially reduced receptor activity or predominantly inactive receptor conformational equilibria. Together, these observations indicate that the scaffold protein can directly or indirectly influence the conformational equilibria of A_2A_AR in the smallest nanodiscs. At the same time, the magnitude of this effect does not appear to be sufficient to overcome the stronger influence of antagonist binding, as highly similar inactive conformational equilibria were observed across all three nanodisc systems for antagonist-bound A_2A_AR. The nearly identical conformational equilibria observed for A_2A_AR in MSP1D1 and MSP1E3D1 nanodiscs containing a 70:30 molar ratio of POPC and POPS argue against possible influences from direct MSP–A_2A_AR interactions in determining the receptor’s conformational equilibrium in these conditions (Fig. 1).

Our systematic analysis of the effects of nanodisc size and lipid composition on A_2A_AR conformational equilibria also provides a valuable experimental benchmark for the development and validation of computational approaches aimed at understanding how membrane environments influence GPCR energy landscapes. The present results, together with previously published experimental studies^40,42,44^, highlight several challenges in accurately capturing the effects of lipid composition and nanodisc architecture in molecular simulations. For example, simulations reported by Vattulainen and colleagues concluded that unliganded (apo) A_2A_AR in nanodiscs composed entirely of POPC and formed from MSP1E1, which has an ∼11 nm diameter comparable to nanodiscs formed from MSP1D1, adopted a more active conformational ensemble.^51^ Increasing the nanodisc diameter to ∼19 nm in simulations shifted the receptor conformational equilibria toward a predominantly inactive conformational ensemble.^51^ These observations differ from both experimental measurements reported previously^40,42,44^ and those reported here, which show that A_2A_AR exhibits substantially reduced activity in MSP1D1 nanodiscs containing only POPC lipids (Figs. 1, 2 and 4), conditions that are highly similar to those used in the reported simulations. Moreover, we do not observe a simple, clear relationship between nanodisc size and receptor conformational equilibria, underscoring that even nanodiscs containing binary lipid mixtures are complex. Further, previous studies also demonstrated differences between the conformational ensembles of A_2A_AR in lipid nanodiscs and vesicles, suggesting that larger nanodiscs may not entirely replicate the conformational equilibria observed in vesicles.^52^ Although the mechanism of these differences remains incompletely understood, the collective body of experimental evidence demonstrates that accurately reproducing GPCR conformational equilibria in lipid environments remains an important challenge for current computational methodologies.

A major mechanism contributing to the reduced availability of anionic lipids for interactions with A_2A_AR was identified from the molecular simulations as the formation of small, transient clusters of anionic lipids (Fig. 5). POPS formed substantially more clusters containing two, three, or more anionic lipids than POPG, and the number of these clusters increased with increasing nanodisc size. This trend correlated with the experimental observation that progressively more amounts of POPS were required to populate an active A_2A_AR conformational ensemble in larger nanodiscs. Heterogeneous spatial organization of anionic lipids within membranes has been reported previously and investigated using both experimental^46^ and computational^47,53,54^ approaches. Prior studies primarily focused on the role of cations, particularly divalent cations such as Ca^2+^, in promoting the formation of anionic lipid clusters^46,47,53,54^ and altering the timescale of molecular dynamics at the membrane–water interface^55^. In the present study, we observed very dynamic yet thermodynamically significant PS–PS interactions in the absence of calcium ions. Monovalent cations such as Na^+^ also did not contribute significantly to cluster formation (Supplementary Fig. 8). These observations suggest a potential mechanistic basis for the longer-lived anionic lipid clusters previously observed in the presence of Ca^2+^. Specifically, rapidly forming and dissolving POPS nanoclusters may serve as transient nucleation sites that facilitate formation of larger and more stable lipid assemblies upon the introduction of Ca^2+^. Even in the absence of Ca^2+^, the clustering of PS lipids likely lowers their chemical activity relative to PG.

A second mechanism contributing to reduced anionic lipid availability was identified as direct interactions between basic residues of the membrane scaffold proteins and anionic lipid headgroups (Fig. 6). Previous studies combining molecular dynamics simulations and EPR spectroscopy demonstrated that cholesteryl hemisuccinate (CHS), which contains a carboxylic group that is negatively charged near physiological pH, preferentially associated with the MSP belt in nanodiscs formed from MSP1D1, reducing its accessibility to the nanodisc interior relative to cholesterol^48^. Thus our observations with POPS and POPG may also extend to additional lipids or membrane-embedded molecules containing negatively charged chemical groups.

The combined contributions of both lipid clustering and lipid–MSP interactions may additionally provide a framework for understanding previous observations regarding the lipid dependence of receptor thermal stability in nanodiscs. We previously reported a systematic analysis of A_2A_AR thermal stability in MSP1D1 nanodiscs containing defined binary mixtures of zwitterionic and anionic lipids, including comparing compositions containing POPS, POPG, and POPA.^40^ Among these mixtures, nanodiscs containing POPS produced the greatest thermal stabilization of A_2A_AR, and receptor melting temperatures increased as the proportion of POPS was increased.^40^ The ability of POPS to engage in both receptor interactions and lipid clustering suggests that lipid organization and MSP-lipid interactions may contribute to the stabilization of membrane proteins embedded in nanodiscs.

While the present study focused on lipid–lipid and lipid–protein interactions within MSP-based nanodiscs, our findings may likely extend to additional membrane-mimetic systems. Since the introduction of MSP nanodiscs, numerous alternative lipid nanoparticles have been developed for structural and biophysical studies of membrane proteins. Among these are the Salipro nanoparticles, which are assembled from the saposin-A (SapA) scaffold protein^56,57^. Analogous to MSPs, SapA forms a belt-like structure around a lipid bilayer. Each SapA monomer contains multiple Lys and Arg residues, and two SapA molecules assemble to form the nanoparticle^58^, thus there are numerous sites for potential interactions between the SapA scaffold and lipids within the nanoparticles. Polymer-based systems such as styrene-maleic acid lipid particles (SMALPs)^59–62^ provide another interesting comparison. The more widely used poly(styrene) co-maleic acid molecule and many of its derivatives contain chemical groups that are negatively charged at physiological pH, raising the possibility that the belt encompassing SMALPs may also influence internal lipid organization. Future studies will be needed to determine the extent to which lipid clustering and scaffold-mediated lipid organization are general features of membrane-mimetic systems.

Replacement of MSP1D1 basic residues identified in simulations as interacting with anionic lipids altered the conformational equilibria of A_2A_AR, indicating that lipid accessibility within nanodiscs can be controlled through rational modification of the scaffold protein. These findings suggest that membrane scaffold proteins can be engineered not only to control nanodisc size, but also to tune lipid organization and lipid availability within nanodiscs. Such an approach could provide a strategy for designing membrane mimetics with tailored biophysical properties, for example, the insertion of additional basic residues into scaffold proteins to intentionally increase the amounts of anionic lipids within nanodiscs.

Finally, the sequence of MSP1D1 is derived entirely from the central region of the polypeptide chain for Apo-A1 (Fig. 8). The extensive sequence overlap between MSPs and Apo-A1 raises the possibility that the principles identified in this study may extend beyond nanodiscs to native biological lipid–protein assemblies. Exploring these possibilities is an intriguing direction for future investigation.

## Acknowledgements

This work is supported by the National Institutes of Health, NIGMS MIRA grants R35GM138291 (M.T.E.) and R35GM153273 (E.L). A portion of this work was supported by the McKnight Brain Institute at the National High Magnetic Field Laboratory’s AMRIS Facility, which is funded by National Science Foundation Cooperative Agreement No. DMR-1644779 and the State of Florida. A portion of this work was also funded by an NIH award, S10 OD028753, for magnetic resonance instrumentation. The simulation work used Delta at the National Center for Supercomputing Applications through allocation BIO240227 from the Advanced Cyberinfrastructure Coordination Ecosystem: Services & Support (ACCESS) program, which is supported by U.S. National Science Foundation grants #2138259, #2138286, #2138307, #2137603, and #2138296.

## Author Contributions

A.V.W., N.P.A., N.T., and A.P.R. expressed A_2A_AR, prepared samples, recorded NMR data and additional biophysical data and analyzed NMR and biophysical data with input from M.T.E. K.Z. expressed and purified scaffold proteins and assisted in preparing lipid nanodisc samples. B.E. expressed and purified scaffold proteins and contributed to preparing samples of A_2A_AR variants in lipid nanodiscs. J.J. and E.L performed computational modeling, simulations, and analysis of simulation data. M.T.E. designed the study, analyzed NMR and other biophysical data and wrote the manuscript with input from all authors.

## Methods

### Molecular cloning of membrane scaffold proteins

Several variants of the membrane scaffold protein MSP1D1 were obtained from Genscript, specifically MSP1D1[XX], MSP1D1[YY], and MSP1D1[ZZ]. Additional MSP1D1 variants were generated using PCR-based site-directed mutagenesis with the Accuprime *Pfx* SuperMix (ThermoFisher Scientific, Catalog Number: 12344040).

### Expression of membrane scaffold proteins

Plasmids containing genes for the MSP1D1, MSP1E3D1, and MSP1D1βH5, and all MSP variant proteins were cloned into a pET28a vector and transformed into the E. *coli* BL21(DE3) strain. Transformed E. *coli* colonies were selected and used to inoculate 10 ml of Luria-Bertani (LB) broth containing 50 mg/ml Kanamycin in an incubator shaker overnight at 37 ^°^C and 250 rpm. Cultures were transferred to 1L terrific broth (TB) media and grown at 37 ^°^C until the optical density (OD) measured at 600nm reached 0.6-0.8, after which protein production was initiated by adding IPTG to a final concentration of 1 mM. The cells were grown for an additional 3 hours and then harvested by centrifugation. SDS-PAGE was performed to compare samples before and after induction and confirm protein production.

### Purification of membrane scaffold proteins

Expressed membrane scaffold proteins were purified following previously described protocols^2,4,63^. Cells were lysed by using a cell disruptor operating at 27 KPSI with lysis buffer (50 mM Tris-HCl, pH 8.0, 500 mM NaCl, 1% Triton, 1 mM EDTA, and in house prepared protease inhibitor cocktail). 5 mM MgCl2 was added to the lysate and incubated at 4 ^°^C for 30 min. The soluble fraction was collected by centrifugation at 2000 rpm and 4 ^°^C for 30 min. The supernatant was incubated with Ni-NTA resin that was prewashed with buffer A (50 mM Tris-HCl, pH 8.00, 500 mM NaCl) and 1% Triton X-100 for 3 h at 4 ^°^C. Next, the slurry was passed through a gravity column and the resin bed was sequentially washed with 10 column volumes (CV) of buffer A and 1% Triton, 10 CV of buffer A and 50 mM sodium cholate, 10 CV of buffer A, and 10 CV of buffer B (50 mM Tris-HCl, pH 8.0, 500 mM NaCl, and 20 mM imidazole). The protein was eluted with 3-5 CV of elution buffer (50 mM Tris-HCl, pH 8.0, 500 mM NaCl, and 500 mM imidazole). The pooled elution fractions were exchanged into buffer (50 mM Tris-HCl, pH 8.0, 20 mM NaCl, and 0.5 mM EDTA) via an AKTA Start FPLC (Cytiva) equipped with a desalting column. The protein concentration was measured by monitoring the UV absorbance at 280 nm using a NanoDrop. Tobacco etch virus (TEV) was added to the protein and incubated overnight at 4 ^°^C. Cleaved MSP was isolated by performing a reverse IMAC step with an AKTA Start FPLC equipped with a Histrap affinity column (Cytiva) and collecting the flow-through. Purified MSP was dialyzed against buffer (20 mM Tris-HCl, pH 8.0, 100 mM NaCl, and 0.5 mM EDTA) using snakeskin dialysis tubing with a 10 kDa MWCO for 5 h. The protein was concentrated to 1 mM and frozen in liquid nitrogen for future use.

For purification of all MSP variants, a similar protocol was used with minor modifications. Prior to the addition of TEV, the NaCl concentration was increased to 300 mM to minimize protein aggregation. The buffer compositions of all other reagents were same.

### A_2A_AR[A289C] expression and purification

High expressing colonies were selected from a microscale expression assay^64,65^ and used for production of A_2A_AR[A289C] in *P. pastoris*. Production and purification of A_2A_AR[A289C] followed previously published protocols^63–65^. Glycerol stocks were used to inoculate 4 ml cultures of buffered minimal glycerol (BMGY) for 48 h in an incubator shaker operating at 30 ^°^C and 200 rpm. Each of these cultures were then used to inoculate 50 ml of BMGY medium and grown for an additional 60 h at 30 °C and 200 rpm. Subsequently, each 50 ml culture was used to inoculate 500 ml of BMGY medium and incubated for 48 h at 30 °C and 200 rpm. Cultures were centrifuged at 6000 rpm for 15 min, and the cell pellets were resuspended in 500 ml of buffered minimal methanol (BMMY) medium without methanol. After 6 h of cell growth to remove trace amounts of glycerol, the temperature was reduced to 28 ^°^C and protein production was initiated by adding 25 ml of 10% w/v of methanol (final methanol concentration of 0.5% w/v per 500 ml culture). Two successive methanol additions were made at 12 h intervals for a total expression time of 36 hrs. Cells were harvested by centrifugation at 6000 rpm and 4 ^°^C for 15 min and stored at -80 ^°^C or lysed immediately. Cell lysis was performed using a cell disruptor operating at 40 kpsi, and the cells were resuspended in lysis buffer (50 mM sodium phosphate, pH 7.0, 100 mM NaCl, 5% w/v glycerol, and an in-house-prepared protease inhibitor cocktail). The membranes were pelleted using an ultracentrifuge operating at 35000 rpm and 4 °C for 20 min. Isolated membrane fractions were frozen and stored at -80 °C or used for immediate extraction of A_2A_AR[A289C].

The isolated membranes were resuspended in high-salt buffer (10 mM HEPES, pH 7.0, 10 mM KCl, 20 mM MgCl2, 1 M NaCl, 4 mM theophylline and 2 mg/mL iodoacetamide) and an in-house-prepared protease inhibitor cocktail solution for 1 h at 4°C. The suspension was centrifuged at 35000 rpm 4 ^°^C for 20 mins to separate the solid membranes. The membranes were then resuspended in high salt buffer in the presence of 1 mM theophylline and in-house prepared protease inhibitor cocktail and incubated for 30 mins at 4 ^°^C followed by the addition of solubilization buffer (50 mM HEPES pH 7.0, 500 mM NaCl, 0.5% w/v n-Dodecyl-b-D-Moltopyranoside (DDM), and 0.05 % cholesteryl hemi succinate (CHS). The solution was allowed to mix gently for 6 h at 4 ^°^C. The soluble fraction was separated by centrifugation, and the supernatant was incubated with Co^+2^-charged affinity resin and 30 mM imidazole at 4 ^°^C for 12 h with gentle rotation.

After overnight incubation, the Co^+2^ resin was isolated by centrifugation at 2000 rpm and 4 ^°^C for 15 min. The isolated resin slurry was subsequently washed with 25 CV of wash buffer 1 (50 mM HEPES pH 7.0, 500 mM NaCl, 10 mM MgCl_2_, 30 mM imidazole, 8 mM ATP, 0.05% DDM, and 0.005% CHS), and washed 2 subsequent times with 25 CV each of wash buffer 2 (25 mM HEPES pH 7.0, 250 mM NaCl, 5% glycerol, 30 mM imidazole, 0.05% DDM, 0.005% CHS, and an excess of ligand). A_2A_AR was eluted with buffer (50 mM HEPES pH 7.0, 250 mM NaCl, 5% glycerol, 300 mM imidazole, 0.05% DDM, 0.005% CHS, and ligand). Pooled eluted fractions were combined and exchanged into buffer (25 mM HEPES pH 7.0, 75 mM NaCl, 0.05% DDM, 0.005% CHS, and ligand) using a PD-10 desalting column. All buffers were prepared with a saturating concentration of the required ligand. For ^19^F-NMR experiments, the final sample buffer contained 100 µM trifluoroacetic acid (TFA).

### ^19^F-NMR labeling using an in-membrane chemical modification approach

Preparation of [A289C^TET^] followed the protocol described above for purification of [A289C] using an in-membrane chemical modification approach^38,41^ to incorporate a ^19^F-NMR probe. Prior to solubilization, the isolated membranes were resuspended in high-salt buffer (10 mM HEPES, pH 7.0, 10 mM KCl, 20 mM MgCl2, 1 M NaCl, 4 mM theophylline) and incubated with 1 mM 4,4’-dithiodipyridine (aldrithiol-4) and an in-house-prepared protease inhibitor cocktail solution for 1 h at 4 °C. Note that iodoacetamide was not added. The suspension was centrifuged at 35000 rpm and 4 ^°^C for 20 mins to separate the solid membranes. The membranes were then resuspended in high salt buffer to remove any excess aldrithiol and incubated with 1 mM of 2,2,2-trifluoroethaethiol (TET) for 1 h at 4 ^°^C. The mixture was centrifugated using the same conditions in the previous step. Extraction and purification of [A289C^TET^] followed the steps as described above.

## Assembly of lipid nanodiscs containing A_2A_AR

Lipid nanodiscs were assembled by adapting protocols from earlier studies.^40,44,63^ Lipid stocks in chloroform were used to make a lipid film and then vacuum dried for 16 h. The dried lipid film was resuspended in cholate buffer (25 mM Tris-HCL, Ph 8.0, 200 mM sodium cholate and 150 mM NaCl) to a final phospholipid concentration of 100 mM. For nanodiscs without A_2A_AR, purified MSP and lipids were incubated at ratios optimized for each differently sized nanodisc. For MSP1D1 and all MSP1D1 variants, the molar ratio of MSP to lipids was 1:50; for MSP1E3D1, the molar ratio of MSP to lipids was 1:120; and for MSP1D1βH5, the molar ratio of MSP to lipids was 1:40. Lipid stocks and purified membrane scaffold proteins were incubated together for 1-2 h at 4 ^°^C followed by overnight incubation of pre-washed biobeads at 4 ^0^C. The nanodisc solution was then separated from biobeads and buffer exchanged using nanodisc desalting buffer (25 mM HEPES pH 7.0, 75 mM NaCl) prior to any experiments.

For A_2A_AR reconstituted nanodiscs, purified A_2A_AR in DDM/CHS detergent micelles was mixed with purified MSP and detergent-solubilized lipids of choice at optimized receptor: MSP: lipid ratios. For MSP1D1, (A_2A_AR: MSP: lipids = 1:5:250), MSP1E3D1 (A_2A_AR: MSP: lipids = 1:5:600), and MSP1D1βH5 (A_2A_AR: MSP: lipids = 1:5:200) molar ratios were incubated for 2-3 h at 4 ^°^C followed by addition of pre-washed biobeads and overnight incubation at 4 ^°^C. Next, the solution was separated from biobeads and incubated with Ni-NTA resin overnight 4 ^°^C to separate receptor reconstituted nanodiscs from empty nanodiscs. The resin was collected using a gravity column and washed with 2CV of nanodisc wash buffer (50 mM HEPES, pH 7.0, 150 mM NaCl, and 10 mM Imidazole). A_2A_AR reconstituted nanodisc were eluted using nanodisc elution buffer (50 mM HEPES pH 7.0, 150 mM NaCl, 300 mM Imidazole, and saturating amounts of the ligand of interest) and buffer exchanged to desalting buffer (25 mM HEPES pH 7.0, 75 mM NaCl, and ligand) using PD-10 columns. For all A_2A_AR [A289C^7.54^] nanodisc samples 100 µM TFA was added to the desalting buffer as an internal standard.

## ^19^F-NMR and ^31^P-NMR in aqueous solutions

Assembled A_2A_AR [A289C^TET^] lipid nanodisc samples were concentrated to ∼ 200 uM in 300 µL using a Vivaspin-6 concentrator with a molecular weight cut off of 30 kDa. 10% v/v D2O was added to the sample and gently mixed before loading into a 5 mm Shigemi NMR tube. All ^19^F and ^31^P NMR data were collected on a Bruker Avance III HD spectrometer operating at ^1^H Larmor frequency of 600 MHz equipped with a Bruker 5 mm BBFO probe. The temperatures at which the NMR spectra were recorded varied from 280-300 K and mentioned under each figure label. A standard sample of 4 % methanol in D4-MeOH was used for temperature calibration. All 1D ^19^F NMR experiments were recorded with a data size of 32k complex points and an acquisition period of 360 ms, and 16k scans with a 0.3 s recycle delay. All ^31^P NMR spectra were recorded with 900 ms acquisition time, 2k scans, and 0.3 s recycle delay.

### Gα_S_ expression and purification

The human Gα_S_ plasmid containing an N-terminal polyhistidine tag and an N-terminal TEV cleavage site was transformed into the BL21(DE3)-RIL *E. coli* strain. A selected colony was grown in 5 mL of LB with 0.2% glucose, 34 ug/ml chloramphenicol and 100 ug/ml carbenicillin for 8 h at 200 rpm. The cultures were used to inoculate 75 mL of LB and were grown at 30 °C overnight at 200 rpm. The cells were centrifuged at 3900 rpm for 5 min at 4 ^°^C and subsequently resuspended in 1 L of LB containing 34 ug/ml chloramphenicol and 100 ug/ml carbenicillin. The cultures were grown at 30 °C until the OD_600_ reached 0.8, after which protein expression was initiated by adding IPTG to a final concentration of 50 µM. The cells were grown overnight at 25 ^°^C at 200 rpm. After 16 h, cells were harvested by centrifugation at 6000 rpm and 4 °C. Post-induction and pre-induction aliquots of cell culture were analyzed by SDS-PAGE to confirm successful protein expression.

The cells were resuspended in buffer (25 mM Tris-HCl, pH 8.0, 150 mM NaCl, 1 mM MgCl_2_, 5 mM GDP, in-house protease inhibitor cocktail) and lysed via sonication. The suspension was centrifuged at 25,000 rpm and 4 ^°^C for 30 min, and the lysate was incubated with Ni-NTA resin overnight at 4 ^°^C. The Ni-NTA resin was washed with 20 CV of wash buffer (25 mM Tris-HCl pH 8.0, 500 mM NaCl, 1 mM MgCl_2_, 5 mM imidazole, 5 mM GDP, and 2 mg/mL iodoacetamide) and eluted with elution buffer (25 mM Tris-HCl pH 8.0, 250 mM NaCl, 1 mM MgCl_2_, 250 mM imidazole, 10% (v/v) glycerol, and 5 mM GDP). The pooled elution fractions were exchanged into desalting buffer (25 mM Tris-HCl, pH 8.0, 100 mM NaCl, 1 mM MgCl_2_, 10% (v/v) glycerol, 5 mM GDP), followed by incubation overnight with TEV protease at a Gα_S_:TEV molar ratio of 1:50. Gα_S_ was incubated with Ni-NTA resin for 30 min at 4 ^°^C. Gα_S_ with a cleaved HIS tag was then collected using a gravity column. Gα_S_ was further purified by size exclusion chromatograhy using a Superdex 200 Increase column (Cytiva) equilibrated with buffer (20 mM HEPES, pH 8.0, 150 mM NaCl, 5 mM MgCl_2_, 1 mM EDTA, 10% (v/v) glycerol, and 5 mM GDP). Purified fractions were collected, analyzed by SDS-PAGE and either used immediately or frozen in liquid nitrogen for future use.

### Mini-Gα_S_ expression and purification

Mini-Gαs was expressed and purified based on previously published protocols.^66^ The mini-Gαs plasmid was transformed into the BL21(DE3)-RIL *E.coli* strain and expressed in TB medium with 0.2% glucose, 34 ug/ml Chloramphenicol, 100 ug/ml Carbenicillin, and 5 mM MgSO_4_. Protein expression was initiated by adding 50 uM IPTG. The cells were then incubated overnight at 25 ^°^C and 250 rpm. The cells were isolated by centrifugation at 6000 rpm and 4 ^°^C for 15 mins, resuspended in buffer (40 mM HEPES, pH 7.5, 100 mM NaCl, 10 mM imidazole, 10% v/v glycerol, 5 mM MgCl_2_, 50 μM GDP, and in-house protease inhibitor cocktail) and lysed by sonication. The lysate was centrifuged at 2000 rpm for 30 min at 4 ^°^C, and the supernatant was loaded on a His-Trap Ni-NTA column (Cytiva) pre-equilibrated with binding buffer (40 mM HEPES, pH 7.5, 100 mM NaCl, 10 mM Imidazole, 10% v/v Glycerol, 5 mM MgCl_2_, and 50 uM GDP). Fractions of purified protein were eluted using buffer (20 mM HEPES, pH 7.5, 500 mM NaCl, 500 mM Imidazole, 10% v/v Glycerol, 5 mM MgCl_2_, and 50 uM GDP). The pooled elution fractions were buffer exchanged using a HiPrep 26/10 column (Cytiva) equilibrated with buffer (20 mM HEPES, pH 7.5, 100 mM NaCl, 10% v/v glycerol, 1 mM MgCl_2_, and 10 µM GDP). Mini-Gα_S_ was incubated overnight with TEV at a molar ratio of 1:100 mini-Gα_S_:TEV. The digested mini-Gα_S_ was removed via reverse IMAC by incubating the sample with Ni-NTA resin in the presence of 30 mM imidazole for 1 h at 4 °C. Fractions of cleaved mini-Gα_S_ were collected using a gravity column and further purified via gel filtration with a HiLoad Superdex 75 column (Cytiva) equilibrated with 10 mM HEPES (pH 7.5), 100 mM NaCl, 10% v/v glycerol, 1 mM MgCl_2_, 1 µM GDP, and 0.1 mM TCEP. Samples were frozen in liquid nitrogen and stored at -80 °C. Prior to using the samples, they were buffer exchanged using a PD-10 desalting column equilibrated with 25 mM HEPES, pH 7.0, 75 mM NaCl, 5 mM MgCl_2_, and 1 µM GDP. Purified mini-Gα_S_ was concentrated to 1 mM using a centrifugal concentrator with a 10 kDa MWCO.

## GTP Hydrolysis Assay

The GTPase-Glo sensor assay (Promega) was used to quantify rates of the GTP hydrolysis for agonist and antagonist-bound A_2A_AR in lipid nanodiscs formed from different membrane scaffold proteins, following the manufacturer’s protocol. A_2A_AR reconstituted in lipid nanodiscs was mixed with purified full-length Gα_S_ at a molar ratio of 4:1 in the presence of 5 uM GTP in buffer (25 mM HEPES, pH 7.0, and 75 mM NaCl). The reaction mixtures were incubated in a 384-well plate at room temperature for 1h. The final GTP turnover luminescence rates were determined using CLARIOstar (BMG Labtech) plate-reader operating at 25 ^°^C.

## Dynamic Light Scattering (DLS) Measurements

The particle sizes of empty lipid nanodiscs assembled with MSP1D1 and MSP1D1 variants were analyzed via DLS by diluting the samples to 10 uM with buffer (25 mM HEPES pH 7.0, 75 mM NaCl). DLS measurements were collected using a Malvern Zetasizer Nano-ZS instrument operating at 25 ^°^C. The volume distribution of particles was determined by Zetasizer software 7.3 (Malvern Panalytical).

## Radioligand Binding Assays

Membrane evaluation and heterologous competition binding assays were measured for A_2A_AR reconstituted in nanodiscs assembled with MSP1D1, MSP1D1βH5, or MSP1E3D1 scaffold proteins and defined mixtures of POPC and POPS lipids. Nanodisc samples were incubated in buffer containing 25 mM HEPES pH 7.0, 75 mM NaCl, [^3^H] CGS21680 (American radiolabeled chemicals, SKU: ART 0884-250 µCi) and increasing amounts of NECA for 60 min at 25 °C. The binding reaction was terminated by filtration with a Microbeta filtermat-96 cell harvester (PerkinElmer). Radioactivity for each nanodisc sample was counted using a MicroBeta2 microplate counter (PerkinElmer). NECA binding affinities (K_i_) were determined using heterologous competition binding experiments. Radioligand experiments were conducted in triplicate and IC_50_ values determined using a nonlinear, least-square regression analysis (Prism 11; GraphPad Software, Inc.). The IC_50_ values were converted to K_i_ values using the Cheng−Prusoff equation^67^. Error bars for each sample were calculated as the standard error of mean (s.e.m) for *n* = 3 independent experiments.

## Simulation Methods System Setup

The initial structure for the receptor was based on the coordinates of the crystal structure of A_2A_AR in a ternary complex with the full agonist NECA and an intracellular engineered G protein (PDB: 5G53)^43^. The starting configuration was created by deleting the G protein *in silico* in order to simulate the fully active receptor, but with the intracellular face accessible to anionic lipid headgroups, as reported in previous work^40^. Modeller 10.5^68^ was used to rebuild residues 147 to 158 and residues 212 to 223 that were not resolved in the receptor PDB structure. Two membrane compositions, POPC:POPS (80:20 molar ratio) and POPC:POPG (80:20 molar ratio), were combined with three Membrane Scaffold Protein (MSP) types: MSP1D1ΔH5, MSP1D1, and MSP1E3D1 (Supplementary Table 1), yielding six nanodisc systems in total. A_2A_AR was incorporated into the lipid bilayer enclosed by the MSP belt in all cases except MSP1D1ΔH5, whose limited size was not able to accommodate the receptor using the membrane assembly tools described below. Ten systems in total were simulated: six nanodisc systems without protein (three different belt proteins and two different lipid compositions each) and four nanodiscs with the receptor (two different belt proteins and two different lipid compositions each).

Each system was prepared individually using the membrane builder (nanodisc builder) in CHARMM-GUI^69,70^. The membrane nanodisc systems in the presence and absence of A_2A_AR were solvated with TIP3P^71^ water molecules. Na^+^ ions were added to neutralize the system, and additional Na^+^ and Cl^–^ ions were added to maintain 0.15 M ionic concentration. The MSP1D1ΔH5 systems contained approximately 197,000 atoms and measured around 12.6 × 12.6 × 12.6 nm^3^. The MSP1D1 systems contained approximately 264,000 atoms and measured around 14.0 × 14.0 × 14.0 nm^3^. The MSP1E3D1 systems contained approximately 472,000 atoms and measured around 17.0 × 17.0 × 17.0 nm^3^.

## Molecular Dynamics (MD) Simulation Details

The systems in Supplementary Table 1 were run with GROMACS version 2025^72,73^. These systems were subjected to a series of energy minimization and equilibration stages with the input files generated from the CHARMM-GUI membrane builder^69^. The CHARMM36 force field parameters were used for protein, lipids, salt (0.15 M NaCl), and explicit TIP3P water^74^. The ligand NECA was modeled with the CHARMM general force field^75^. The initial configuration was relaxed by 5000 steps of steepest descent. The next two simulation stages used a 1 fs timestep for a total of 500 ps (250 ps × 2) of simulation time in isothermal−isochoric (NVT), the third stage used 1 fs timestep for 250 ps in isothermal−isobaric (NPT) ensemble, and the last three stages used a 2 fs timestep for a total of 1500 ps (500 ps × 3) simulation length in the NPT ensemble. During these relaxation stages position restraints were applied to the heavy atoms of the protein backbones and the ligand with a force constant of 4000 kJ mol^−1^ nm^−2^. The restraints were reduced to 1000 kJ mol^−1^ nm^−2^ during the third step (NPT) and further to 50 kJ mol^−1^ nm^−2^ during subsequent NPT equilibrations to facilitate gradual structural relaxation. The heavy atoms of the protein sidechains were positionally restrained with a force constant of 2000 kJ mol^−1^ nm^−2^, which was gradually reduced to 0 in a stepwise manner. Position restraints along the z direction (parallel to the membrane normal) were applied to the phosphorus atoms of the lipids with a force constant of 1000 kJ mol^−1^ nm^−2^ and gradually reduced to 0 in the last NPT equilibration. The temperature was maintained at 298.15 K using the v-rescale thermostat^76^ with a coupling constant of τ = 1.0 ps. During the NPT stages the semi-isotropic pressure was maintained at 1 bar using the C-rescale^77^ barostat, with a pressure coupling constant of τ = 5.0 ps and a compressibility of 4.5 × 10^−5^ bar^−1^. Orthorhombic periodic boundary conditions were applied to each system. The nonbonded interaction neighbor list was updated every 20 steps. Short-range electrostatic and van der Waals interactions were computed with a 1.2 nm cutoff. The particle-mesh Ewald (PME) method^78,79^ was employed to handle the long-range electrostatics every timestep with a Fourier grid spacing of 1.2 Å and fourth-order interpolation. The long-range dispersion of the vdW interactions were smoothly switched to zero between 10 and 12 Å using a force-switch scheme. The bonds involving hydrogen atoms were constrained using the linear constraint solver (LINCS) algorithm^80^. Finally, the production simulations were performed under NPT conditions for 5.0 μs per system, with all restraints removed. During the initial stages of the production simulation, MSP nanodiscs underwent significant structural relaxation, deviating markedly from their initial circular geometry, as observed previously in simulations of apolipoprotein A1^81^. The three-dimensional shape anisotropy (or global ellipticity) was quantitatively evaluated (Supplementary Table 17), and data were analyzed after this structural relaxation was complete.

## Modeling Apo A1

The Apo A1 system was first energy-minimized using 5000 steps of the steepest descent algorithm. This was followed by NVT equilibration with a 1 fs time step for 250 ps. During equilibration, position restraints were applied to the protein backbone heavy atoms with a force constant of 400 kJ mol^-1^ nm^-2^, and to the side chains with a force constant of 40 kJ mol^-1^ nm^-2^. Subsequently, a 1 ns production simulation was carried out under NPT conditions without position restraints, with coordinates written every 10 ps. All other simulation parameters were consistent with those described above. The frame at 10 ps was extracted for image rendering.

## Visualization and analysis

Visual Molecular Dynamics (VMD)^82^ was used to visualize the MD trajectories. PyMOL^83^ was used for visualization and image renderings. To quantify the three-dimensional anisotropic shape fluctuations of MSP nanodiscs, a principal component-based geometric analysis was performed on the Cα atomic coordinates extracted from trajectory frames. For each frame, Cartesian coordinates were first centered by subtracting the instantaneous center of geometry of the selected atoms. A covariance matrix of the centered coordinates was then constructed and subjected to eigen decomposition to obtain eigenvalues and eigenvectors, which define the principal axes of the structural distribution. The eigenvectors corresponding to the largest, second largest, and third largest eigenvalues were defined as the major, (in-plane) minor, and third (membrane-normal, “thickness”) axes, respectively. Atomic positions were projected onto these orthogonal directions, and the spatial extent along each axis was quantified as the difference between the maximum and minimum projected values. The aspect ratio was calculated as the ratio of the major- to minor-axis length. To exclude initial relaxation effects, the first 500 ns of the production trajectories were omitted from analysis (the same procedure was applied in all analyses below). Using this procedure, we evaluated the overall ellipticity of the MSP nanodiscs (Supplementary Table 17).

Hydrogen bond analysis was performed using the Hydrogen Bond Analysis Function implemented in MDAnalysis package^84^. The geometric criteria for hydrogen bonds are a donor-acceptor distance cutoff of 3.0 Å, a distance cutoff of 1.2 Å used for finding donor-hydrogen pairs, and a donor-hydrogen-acceptor angle cutoff of 150°. Hydrogen bonds between POPS (POPG) lipids excluded intramolecular interactions.

In addition, after completing the 5.0 μs production simulation, the final frame of each system was extracted and used to perform an additional 500 ps short production simulation to analyze hydrogen-bond dynamics among POPS or POPG lipids. For POPS lipids, the protonated amino group serves as the hydrogen-bond donor, whereas phosphate oxygens and carboxylate oxygens, and ester oxygens in the headgroup act as hydrogen-bond acceptors. For POPG lipids, the headgroup hydroxyl group serves as the hydrogen-bond donor, while the phosphate oxygens, hydroxyl oxygens, and ester oxygens act as hydrogen-bond acceptors. Trajectories were saved every 10 fs. All other simulation parameters were the same as those described above. The hydrogen bond lifetime was calculated via the time autocorrelation function of the presence of a hydrogen bond in MDAnalysis:

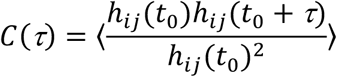

A biexponential was used to fit the time autocorrelation curve to get the hydrogen-bond lifetime:

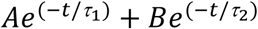

The lifetime was calculated as Aτ_1_ + Bτ_2_. We also estimated the hydrogen-bond lifetime using the trapezoidal rule (Supplementary Table 2).

To characterize the interactions between protein positively charged residues (Arg and Lys) and negatively charged lipids (POPS and POPG), contacts were defined between the positively charged nitrogen atoms of Arg/Lys and the negatively charged phosphate (–PO4^–^) and/or carboxylate (–COO^–^) atoms of the lipids, using a distance cutoff of 4.0 Å. The interaction frequency of Arg and Lys residues in the MSP belt with POPS/POPG was then categorized into three classes: ≥40% of the simulation time (strong), β5% (medium), and < 5% (weak). Non-interacting residues denote those Arg or Lys side chains that are physically shielded from the lipid bilayer due to their specific structural rearrangements in the MSPs.

In prior work^40^, we showed that the H230^6.32^-R291^7.56^ distance served as a reporter for distinguishing between the active and inactive conformations of A_2A_AR. Building upon this foundation, we monitored the H230^6.32^-R291^7.56^ distance across different nanodisc environments in the current simulations, defining it as the minimum distance between their closest side-chain nitrogen atoms. A rightward shift in this distance distribution toward larger values denotes the population of the active receptor conformation. Data analysis and plotting were performed using in-house Python scripts based on publicly hosted Python packages: NumPy^85^, Pandas^86,87^, MDAnalysis^84^, and Matplotlib^88^.

## Data Availability

Source data will be provided with this paper upon its acceptance. NMR source data will be made available through the Open Science Framework or similar repository upon acceptance of this manuscript.

## Code Availability

No new code was generated in this study. Simulation input parameters, initial coordinates and final coordinates will be made available through the Open Science Framework or Zenodo repository upon acceptance of this article.

## References

1 Denisov, I. G., Grinkova, Y. V., Lazarides, A. A. & Sligar, S. G. Directed self-assembly of monodisperse phospholipid bilayer nanodiscs with controlled size. J. Am. Chem. Soc. 126, 3477–3487 (2004). 10.1021/ja0393574

2 Bayburt, T. H., Grinkova, Y. V. & Sligar, S. G. Self-assembly of discoidal phospholipid bilayer nanoparticles with membrane scaWold proteins. Nano Lett. 2, 853–856 (2002). 10.1021/nl025623k

3 Hagn, F., Nasr, M. L. & Wagner, G. Assembly of phospholipid nanodiscs of controlled size for structural studies of membrane proteins by NMR. Nat. Protoc. 13, 79–98 (2018). 10.1038/nprot.2017.094

4 Hagn, F., Etzkorn, M., Raschle, T. & Wagner, G. Optimized phospholipid bilayer nanodiscs facilitate high-resolution structure determination of membrane proteins. J. Am. Chem. Soc. 135, 1919–1925 (2013). 10.1021/ja310901f

5 Zhang, M. et al. Cryo-EM structure of an activated GPCR–G protein complex in lipid nanodiscs. Nat. Struct. Mol. Biol. 28, 258–267 (2021). 10.1038/s41594-020-00554-6

6 Staus, D. P. et al. Structure of the M2 muscarinic receptor–β-arrestin complex in a lipid nanodisc. Nature 579, 297–302 (2020). 10.1038/s41586-020-1954-0

7 Köck, Z. et al. Cryo-EM structure of cell-free synthesized human histamine 2 receptor/Gs complex in nanodisc environment. Nat. Comm. 15, 1831 (2024).

8 Raschle, T. et al. Structural and Functional Characterization of the Integral Membrane Protein VDAC-1 in Lipid Bilayer Nanodiscs. J. Am. Chem. Soc. 131, 17777–17779 (2009). 10.1021/ja907918r

9 Wei, S. et al. Slow conformational dynamics of the human A2A adenosine receptor are temporally ordered. Structure 30, 329–337 (2022). 10.1016/j.str.2021.11.005

10 Lamichhane, R. et al. Single-molecule view of basal activity and activation mechanisms of the G protein-coupled receptor β _2_ AR. Proc. Natl. Acad. Sci. 112, 14254–14259 (2015). 10.1073/pnas.1519626112

11 Krishna Kumar, K., et al. Stepwise activation of a metabotropic glutamate receptor. Nature 629, 951–956 (2024).

12 Huang, S. K. et al. Allosteric modulation of the adenosine A2A receptor by cholesterol. eLife 11, e73901 (2022). 10.7554/eLife.73901

13 Guo, C., Yang, L., Liu, Z., Liu, D. & Wüthrich, K. Two-Dimensional NMR Spectroscopy of the G Protein-Coupled Receptor A2AAR in Lipid Nanodiscs. Molecules 28, 5419 (2023). 10.3390/molecules28145419

14 Dalal, V. et al. Lipid nanodisc scaWold and size alter the structure of a pentameric ligand-gated ion channel. Nat. Comm. 15, 25 (2024). 10.1038/s41467-023-44366-w

15 Dalal, V., Tan, B. K., Xu, H. & Cheng, W. W. Cryo-EM structures of a pentameric ligand-gated ion channel in liposomes. Elife 14, RP106728 (2025).

16 HoWmann, L. et al. The ABC transporter MsbA in a dozen environments. Structure 33, 916–923. e914 (2025).

17 Rahman, M. M. et al. Structural mechanism of muscle nicotinic receptor desensitization and block by curare. Nat. Struct. Mol. Biol. 29, 386–394 (2022).

18 Zarkadas, E. et al. Conformational transitions and ligand-binding to a muscle-type nicotinic acetylcholine receptor. Neuron 110, 1358–1370. e1355 (2022).

19 Laverty, D. et al. Cryo-EM structure of the human α_1_β_3_γ_2_ GABAA receptor in a lipid bilayer. Nature 565, 516–520 (2019).

20 Kim, J. J. et al. Shared structural mechanisms of general anaesthetics and benzodiazepines. Nature 585, 303–308 (2020).

21 Jain, G. & Eddy, M. T. The influence of lipids and biological membranes on the conformational equilibria of GPCRs: Insights from NMR spectroscopy. Curr. Opin. Struct. Biol. 94, 103103 (2025).

22 Jones, A. J. Y., Gabriel, F., Tandale, A. & Nietlispach, D. Structure and dynamics of GPCRs in lipid membranes: physical principles and experimental approaches. Molecules 25, 4729 (2020). 10.3390/molecules25204729

23 Damian, M. et al. Allosteric modulation of ghrelin receptor signaling by lipids. Nature Communications 2021 12:*1* 12, 1-15 (2021). 10.1038/s41467-021-23756-y

24 Gomes, A. A. et al. Lipids modulate the dynamics of GPCR: β-arrestin interaction. Nat. Comm. 16, 4982 (2025).

25 Strohman, M. J. et al. Local membrane charge regulates β_2_ adrenergic receptor coupling to G_I3_. Nat. Comm. 10, 2234 (2019). 10.1038/s41467-019-10108-0

26 Dawaliby, R. et al. Allosteric regulation of G protein–coupled receptor activity by phospholipids. Nat. Chem. Biol. 12, 35–39 (2015).

27 Kimura, T. et al. Recombinant cannabinoid type 2 receptor in liposome model activates g protein in response to anionic lipid constituents. J. Biol. Chem. 287, 4076–4087 (2012).

28 Yen, H.-Y. et al. PtdIns(4,5)P2 stabilizes active states of GPCRs and enhances selectivity of G-protein coupling. Nature 559, 423–427 (2018). 10.1038/s41586-018-0325-6

29 Duncan, A. L., Song, W. & Sansom, M. S. P. Lipid-Dependent Regulation of Ion Channels and G Protein–Coupled Receptors: Insights from Structures and Simulations. Annu. Rev. Pharmacol. Toxicol. 60, 31–50 (2020). 10.1146/ANNUREV-PHARMTOX-010919-023411

30 Song, W., Yen, H.-Y., Robinson, C. V. & Sansom, M. S. P. State-dependent lipid interactions with the A2A receptor revealed by MD simulations using in vivo-mimetic membranes. Structure 27, 392–403 (2019). 10.1016/j.str.2018.10.024

31 Dijkman, P. M. et al. Conformational dynamics of a G protein–coupled receptor helix 8 in lipid membranes. Sci. Adv. 6, eaav8207 (2020).

32 Jacobson, K. A., Suresh, R. R. & Oliva, P. A2A adenosine receptor agonists, antagonists, inverse agonists and partial agonists. Int. Rev. Neurobiol. 170, 1–27 (2023).

33 Zheng, J., Zhang, X. & Zhen, X. Development of Adenosine A _2A_ Receptor Antagonists for the Treatment of Parkinson’s Disease: A Recent Update and Challenge. ACS Chem. Neurosci. 10, 783–791 (2019). 10.1021/acschemneuro.8b00313

34 Young, A. et al. Co-inhibition of CD73 and A2AR Adenosine Signaling Improves Anti-tumor Immune Responses. Cancer Cell 30, 391–403 (2016). 10.1016/J.CCELL.2016.06.025

35 Cekic, C. & Linden, J. Adenosine A2A receptors intrinsically regulate CD8+ T cells in the tumor microenvironment. Cancer Res. 74, 7239–7249 (2014). 10.1158/0008-5472.CAN-13-3581

36 Huang, S. K. et al. Mapping the conformational landscape of the stimulatory heterotrimeric G protein. Nat. Struct. Mol. Biol. 30, 502–511 (2023). 10.1038/s41594-023-00957-1

37 Ye, L., Van Eps, N., Zimmer, M., Ernst, O. P. & Scott Prosser, R. Activation of the A2A adenosine G-protein-coupled receptor by conformational selection. Nature 533, 265–268 (2016). 10.1038/nature17668

38 Sušac, L., Eddy, M. T., Didenko, T., Stevens, R. C. & Wüthrich, K. A2A adenosine receptor functional states characterized by 19F-NMR. Proc. Natl. Acad. Sci. 115, 12733–12738 (2018). 10.1073/pnas.1813649115

39 Clark, L. D. et al. Ligand modulation of sidechain dynamics in a wild-type human GPCR. eLife 6, e28505 (2017). 10.7554/eLife.28505

40 Thakur, N. et al. Anionic phospholipids control mechanisms of GPCR-G protein recognition. Nat. Comm. 14, 794 (2023).

41 Sušac, L., O’Connor, C., Stevens, R. C. & Wüthrich, K. In-membrane chemical modification (IMCM) for site-specific chromophore labeling of GPCRs. Angew. Chem. 127, 15461–15464 (2015). 10.1002/ange.201508506

42 Thakur, N. et al. Membrane mimetic-dependence of GPCR energy landscapes. Structure, S0969212624000376 (2024). 10.1016/j.str.2024.01.013

43 Carpenter, B., Nehmé, R., Warne, T., Leslie, A. G. W. & Tate, C. G. Structure of the adenosine A_2A_ receptor bound to an engineered G protein. Nature 536, 104–107 (2016). 10.1038/nature18966

44 Ray, A. P., Thakur, N., Pour, N. G. & Eddy, M. T. Dual mechanisms of cholesterol-GPCR interactions that depend on membrane phospholipid composition. Structure 31, 836–847 (2023). 10.1016/j.str.2023.05.001

45 Mondal, S., Hsiao, K. & Goueli, S. A. A Homogenous bioluminescent system for measuring GTPase, GTPase activating protein, and guanine nucleotide exchange factor activities. Assay Drug Dev. Technol. 13, 444–455 (2015). 10.1089/adt.2015.643

46 Kijac, A. et al. Lipid−Protein Correlations in Nanoscale Phospholipid Bilayers Determined by Solid-State Nuclear Magnetic Resonance. Biochemistry 49, 9190–9198 (2010). 10.1021/bi1013722

47 Han, K., Kim, S. H., Venable, R. M. & Pastor, R. W. Design principles of PI (4, 5) P2 clustering under protein-free conditions: Specific cation eWects and calcium-potassium synergy. Proc. Natl. Acad. Sci. 119, e2202647119 (2022).

48 Augustyn, B. et al. Cholesteryl Hemisuccinate Is Not a Good Replacement for Cholesterol in Lipid Nanodiscs. J. Phys. Chem. B, 7 (2019).

49 Borhani, D. W., Rogers, D. P., Engler, J. A. & Brouillette, C. G. Crystal structure of truncated human apolipoprotein AI suggests a lipid-bound conformation. Proc. Natl. Acad. Sci. 94, 12291–12296 (1997).

50 Bhale, A. S. & Venkataraman, K. Leveraging knowledge of HDLs major protein ApoA1: Structure, function, mutations, and potential therapeutics. Biomed. Pharmacother. 154, 113634 (2022).

51 Hägg, V., Levental, I., Kaptan, S. & Vattulainen, I. How receptor conformation depends on lipid nanodisc size: Adenosine A2A receptor and implications for class-A GPCR proteins. Biochimica et Biophysica Acta (BBA)-Biomembranes, 184535 (2026).

52 Ray, A. P., Jin, B. & Eddy, M. T. The conformational equilibria of a human GPCR compared between lipid vesicles and aqueous solutions by integrative 19 F-NMR. J. Am. Chem. Soc. 147, 17612–17625 (2025).

53 Jarin, Z., Venable, R. M., Han, K. & Pastor, R. W. Ion-induced PIP2 clustering with Martini3: modification of phosphate–ion interactions and comparison with CHARMM36. J. Phys. Chem. B 128, 2134–2143 (2024).

54 Han, K., Gericke, A. & Pastor, R. W. Characterization of specific ion eWects on PI (4, 5) P2 clustering: molecular dynamics simulations and graph-theoretic analysis. J. Phys. Chem. B 124, 1183–1196 (2020).

55 Valentine, M. L., Cardenas, A. E., Elber, R. & Baiz, C. R. Physiological calcium concentrations slow dynamics at the lipid-water interface. Biophys. J. 115, 1541–1551 (2018).

56 Frauenfeld, J. et al. A saposin-lipoprotein nanoparticle system for membrane proteins. Nat. Methods 13, 345–351 (2016).

57 Chien, C.-T. H. et al. An adaptable phospholipid membrane mimetic system for solution NMR studies of membrane proteins. J. Am. Chem. Soc. 139, 14829–14832 (2017).

58 Popovic, K., Holyoake, J., Pomès, R. & Privé, G. G. Structure of saposin A lipoprotein discs. Proc. Natl. Acad. Sci. 109, 2908–2912 (2012).

59 Dörr, J. M. et al. The styrene–maleic acid copolymer: a versatile tool in membrane research. Eur. Biophys. J. 45, 3–21 (2016).

60 Lee, S. C. et al. A method for detergent-free isolation of membrane proteins in their local lipid environment. Nat. Protoc. 11, 1149–1162 (2016).

61 Knowles, T. J. et al. Membrane proteins solubilized intact in lipid containing nanoparticles bounded by styrene maleic acid copolymer. J. Am. Chem. Soc. 131, 7484–7485 (2009).

62 Craig, A. F. et al. Tuning the size of styrene-maleic acid copolymer-lipid nanoparticles (SMALPs) using RAFT polymerization for biophysical studies. Biochimica et Biophysica Acta (BBA)-Biomembranes 1858, 2931–2939 (2016).

63 Thakur, N., Wei, S., Ray, A. P., Lamichhane, R. & Eddy, M. T. Production of human A_2A_AR in lipid nanodiscs for ^19^F-NMR and single-molecule fluorescence spectroscopy. STAR Protocols 3, 101535 (2022). 10.1016/j.xpro.2022.101535

64 Eddy, M. T. et al. Allosteric coupling of drug binding and intracellular signaling in the A_2A_ adenosine receptor. Cell 172, 68–80 (2018). 10.1016/j.cell.2017.12.004

65 Eddy, M. T. et al. Extrinsic tryptophans as NMR probes of allosteric coupling in membrane proteins: application to the A_2A_ adenosine receptor. J. Am. Chem. Soc. 140, 8228–8235 (2018). 10.1021/jacs.8b03805

66 Lebon, G. et al. Agonist-bound adenosine A_2A_ receptor structures reveal common features of GPCR activation. Nature 2011 474:7352 474, 521-525 (2011). 10.1038/nature10136

67 Cheng, Y. & PrusoW, W. H. Relationship between the inhibition constant (K1) and the concentration of inhibitor which causes 50 per cent inhibition (I50) of an enzymatic reaction. Biochem. Pharmacol. 22, 3099–3108 (1973). 10.1016/0006-2952(73)90196-2

68 Webb, B. & Sali, A. Comparative protein structure modeling using MODELLER. Current protocols in bioinformatics 54, 5.6. 1-5.6. 37 (2016).

69 Lee, J. et al. CHARMM-GUI *Membrane Builder* for complex biological membrane simulations with glycolipids and lipoglycans. J. Chem. Theory Comput. 15, 775–786 (2019). 10.1021/acs.jctc.8b01066

70 Park, S., Choi, Y. K., Kim, S., Lee, J. & Im, W. CHARMM-GUI membrane builder for lipid nanoparticles with ionizable cationic lipids and PEGylated lipids. J. Chem. Inf. Model. 61, 5192–5202 (2021).

71 Jorgensen, W. L., Chandrasekhar, J., Madura, J. D., Impey, R. W. & Klein, M. L. Comparison of simple potential functions for simulating liquid water. J. Chem. Phys. 79, 926–935 (1983). 10.1063/1.445869

72 Abraham, M. J. et al. GROMACS: High performance molecular simulations through multi-level parallelism from laptops to supercomputers. SoftwareX 1, 19–25 (2015).

73 Kutzner, C. et al. More bang for your buck: Improved use of GPU nodes for GROMACS 2018. J. Comput. Chem. 40, 2418–2431 (2019).

74 Huang, J. et al. CHARMM36m: an improved force field for folded and intrinsically disordered proteins. Nat. Methods 14, 71–73 (2017). 10.1038/nmeth.4067

75 Vanommeslaeghe, K. et al. CHARMM general force field: A force field for drug-like molecules compatible with the CHARMM all-atom additive biological force fields. J. Comput. Chem. 31, 671–690 (2009). 10.1002/jcc.21367

76 Bussi, G., Donadio, D. & Parrinello, M. Canonical sampling through velocity rescaling. The Journal of chemical physics 126 (2007).

77 Bernetti, M. & Bussi, G. Pressure control using stochastic cell rescaling. The Journal of Chemical Physics 153 (2020).

78 Darden, T., York, D. & Pedersen, L. Particle mesh Ewald: An *N* ⋅log(*N*) method for Ewald sums in large systems. J. Chem. Phys. 98, 10089–10092 (1993). 10.1063/1.464397

79 Essmann, U. et al. A smooth particle mesh Ewald method. The Journal of chemical physics 103, 8577–8593 (1995).

80. Hess, B., Bekker, H., Berendsen, H. J. C. & Fraaije, J. G. E. M. LINCS: A linear constraint solver for molecular simulations. J. Comput. Chem. 18, 1463–1472 (1997). 10.1002/(SICI)1096-987X(199709)18:12<1463::AID-JCC4>3.0.CO;2-H

81 Pourmousa, M. et al. Tertiary structure of apolipoprotein AI in nascent high-density lipoproteins. Proc. Natl. Acad. Sci. 115, 5163–5168 (2018).

82 Humphrey, W., Dalke, A. & Schulten, K. VMD: visual molecular dynamics. J. Mol. Graph. 14, 33–38 (1996).

83 Schrodinger, LLC. The PyMOL Molecular Graphics System, Version 1.*8* (2015).

84 Michaud-Agrawal, N., Denning, E. J., Woolf, T. B. & Beckstein, O. MDAnalysis: a toolkit for the analysis of molecular dynamics simulations. J. Comput. Chem. 32, 2319–2327 (2011).

85 Harris, C. R. et al. Array programming with NumPy. Nature 585, 357–362 (2020).

86 McKinney, W. Data structures for statistical computing in Python. *scipy* **445**, 51–56 (2010).

87 Team, T. P. D. pandas-dev/pandas: Pandas. Zenodo, February (2020).

88 Hunter, J. D. Matplotlib: A 2D graphics environment. Computing in science & engineering 9, 90–95 (2007).

