## Supplemental Information for "Spatial Organization of Lipids Drives GPCR Conformational Equilibria"

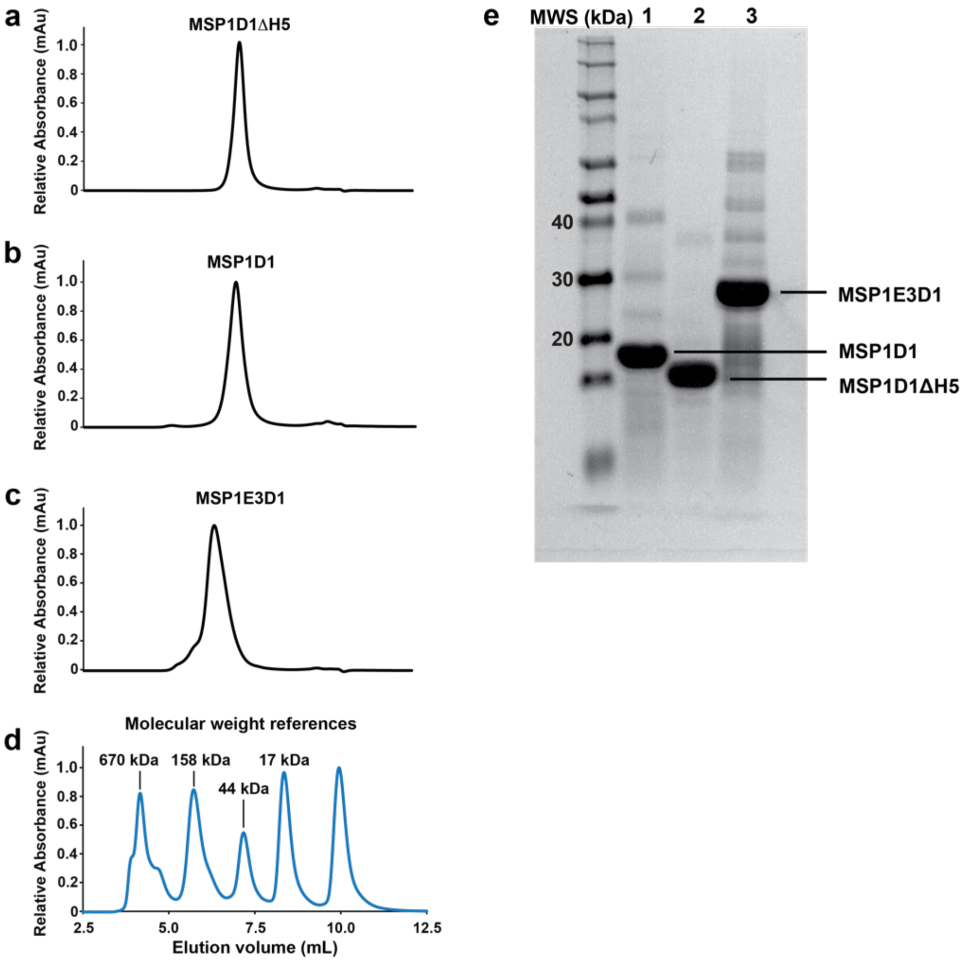


**Supplementary Fig. 1. Biochemical characterization of lipid nanodiscs without $A_{2A}$AR embedded. a-c** Representative analytical size exclusion (aSEC) chromatograms of lipid nanodiscs containing a defined binary mixture of POPC and POPS at a molar ratio of 70:30 formed from the scaffold proteins **a** MSP1D1ΔH5, **b** MSP1D1, and **c** MSP1E3D1. **d** Chromatogram of the molecular standards used to estimate the molecular weight of the nanodisc systems with annotated molecular weights shown. **e** Representative SDS-PAGE gel showing the same nanodisc samples as characterized by aSEC formed from the scaffold proteins MSP1D1 (lane 1), MSP1D1ΔH5 (lane 2) and MSP1E3D1 (lane 3). The lane labeled “MWS” corresponds to molecular weight standards.

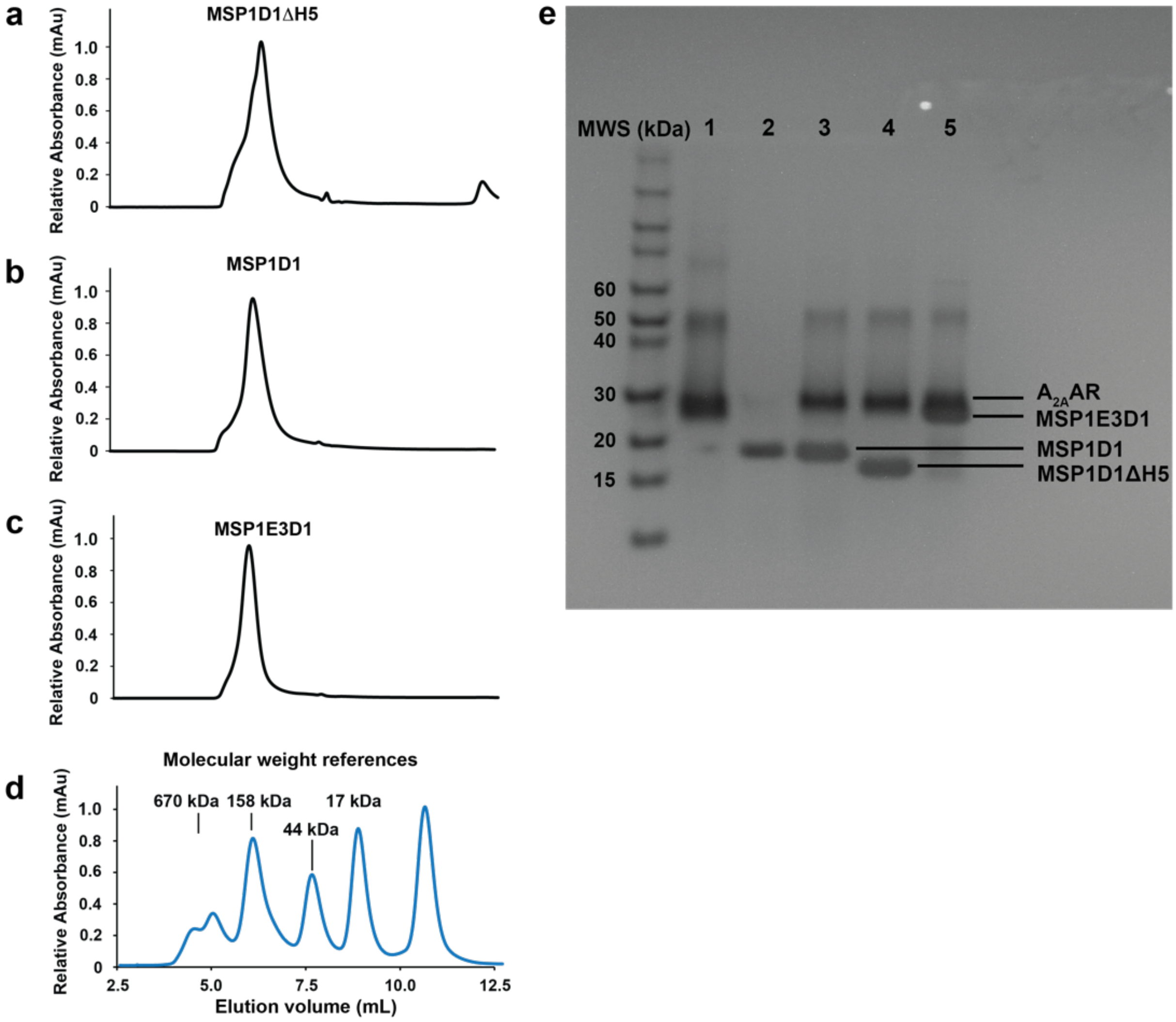


**Supplementary Fig. 2. Biochemical characterization of lipid nanodiscs containing $A_{2A}$AR[A289C]. a-c** Representative analytical size exclusion (aSEC) chromatograms of human $A_{2A}$AR[A289C] in lipid nanodiscs containing a defined binary mixture of POPC and POPS at a molar ratio of 70:30 formed from the scaffold proteins **a** MSP1D1ΔH5, **b** MSP1D1, and **c** MSP1E3D1. **d** Chromatogram of the molecular standards used to estimate the molecular weight of the nanodisc systems with annotated molecular weights shown. **e** Representative SDS-PAGE gel showing purified $A_{2A}$AR[A289C] in DDM/CHS detergent micelles (lane 1), purified MSP1D1 (lane 2), and the same nanodisc samples containing $A_{2A}AR$[A289C] as characterized by aSEC formed from the scaffold proteins MSP1D1 (lane 3), MSP1D1ΔH5 (lane 4) and MSP1E3D1 (lane 5). The lane labeled "MWS" corresponds to molecular weight standards.

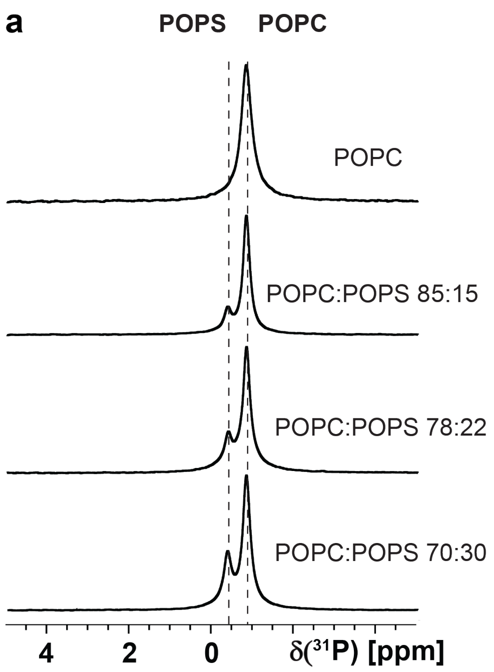


**Supplementary Fig. 3. Validation of nanodisc lipid composition with $^{31}$P-NMR in aqueous solutions.** **a** The 1D $^{31}$P-NMR spectra of $A_{2A}$AR[A289C$^{TET}$] in MSP1D1 nanodiscs containing mixtures of POPC and POPS at different defined molar ratios, as indicated. The vertical dashed lines indicate the chemical shifts for each lipid headgroup, which were consistent with the manufacturer's reported values. The relative intensities of the two observed signals were quantitatively consistent with the expected lipid ratios. **b** The 1D $^{31}$P-NMR spectra of $A_{2A}$AR[A289C$^{TET}$] in MSP1D1 nanodiscs containing mixtures of POPC and POPG at different defined molar ratios, as indicated. Same presentation details as in **a**.

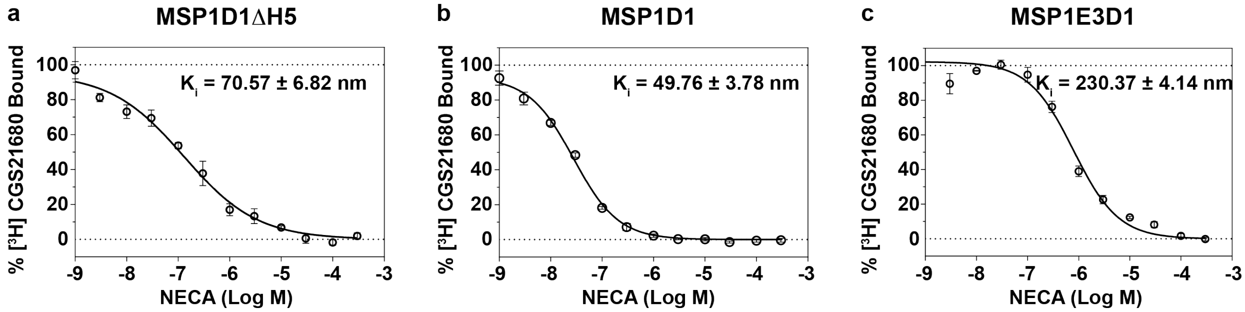


**Supplementary Fig. 4. Pharmacological activity of $A_{2A}$AR[A289C] in three lipid nanodisc systems. a-c** Competitive binding experiments with the agonist NECA are shown for $A_{2A}$AR[A289C] in nanodiscs formed from the scaffold proteins **a** MSP1D1ΔH5, **b** MSP1D1, and **c** MSP1E3D1. Each nanodisc system was formed from a binary lipid mixture of POPC and POPS at a molar ratio of 70:30. The measured $K_i$ values are shown in each panel. Error bars indicate the s.e.m for 3 independent trials done in triplicate.

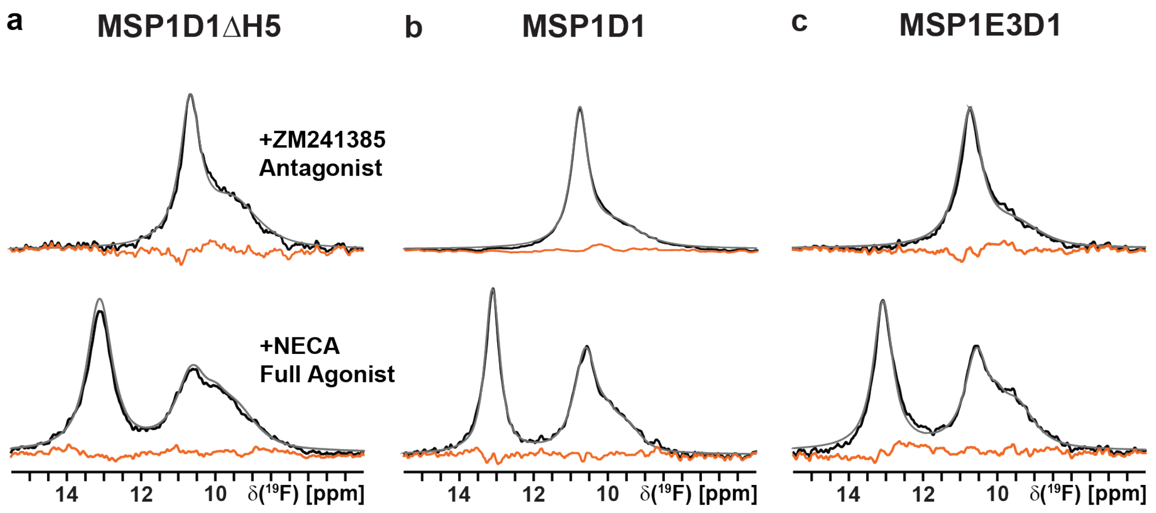


**Supplementary Fig. 5. Residual differences between the raw NMR data and summation of the individually fit components from Figure 1.** Experimental NMR data are shown in black, grey lines superimposed on the spectra are the total sums of the individual deconvolutions, and the orange lines are the calculated differences between the grey and black lines for $^{19}$F-NMR spectra with $A_{2A}$AR[A289C$^{TET}$] in nanodiscs formed from **a** MSP1D1ΔH5, **b** MSP1D1, and **c** MSP1E3D1 scaffold proteins.

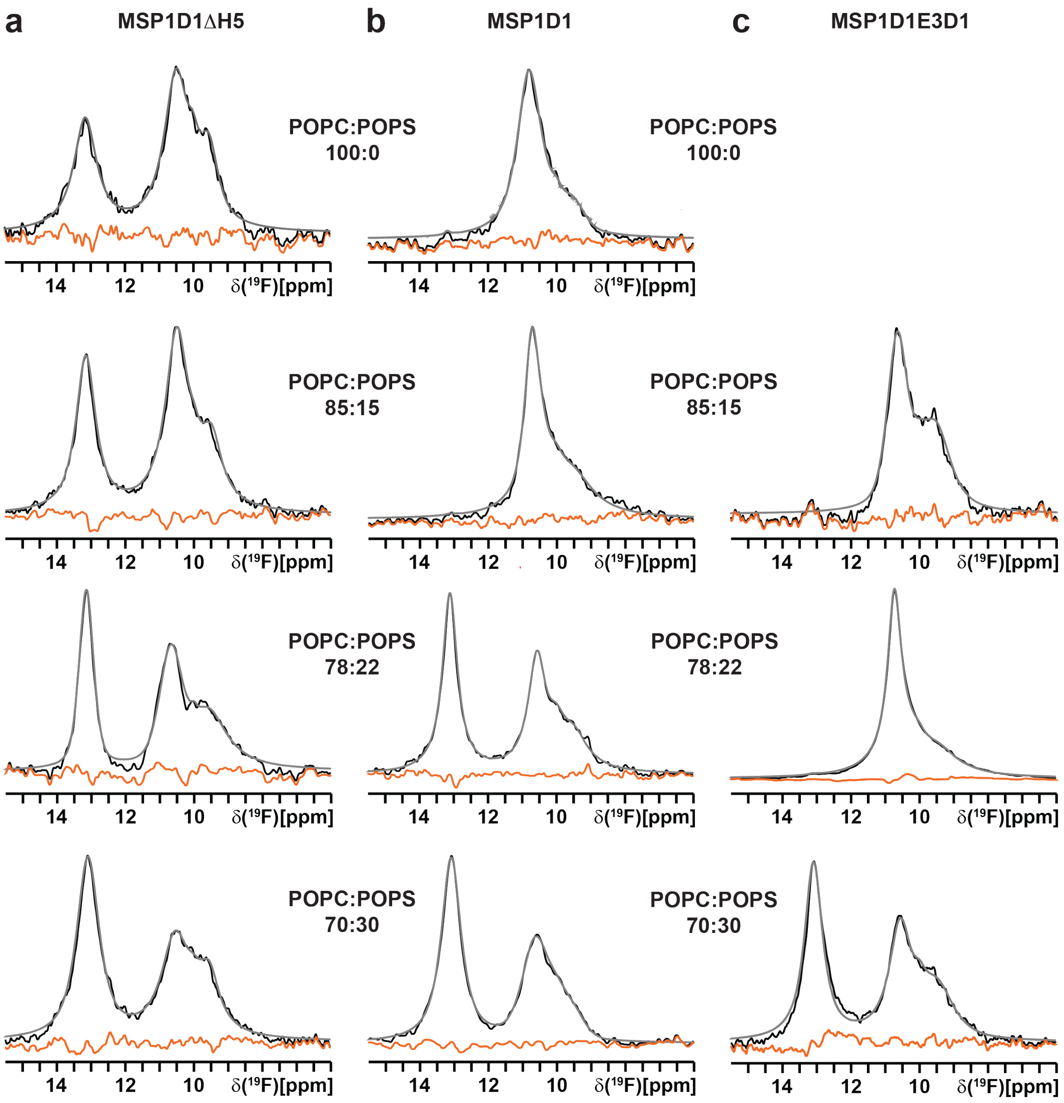


**Supplementary Fig. 6. Residual differences between the raw NMR data and summation of the individually fit components from Figure 2.** Experimental NMR data are shown in black, grey lines superimposed on the spectra are the total sums of the individual deconvolutions, and the orange lines are the calculated differences between the grey and black lines for $^{19}$F-NMR spectra with $A_{2A}$AR[A289C$^{TET}$] in nanodiscs formed from **a** MSP1D1ΔH5, **b** MSP1D1, and **c** MSP1E3D1 scaffold proteins.

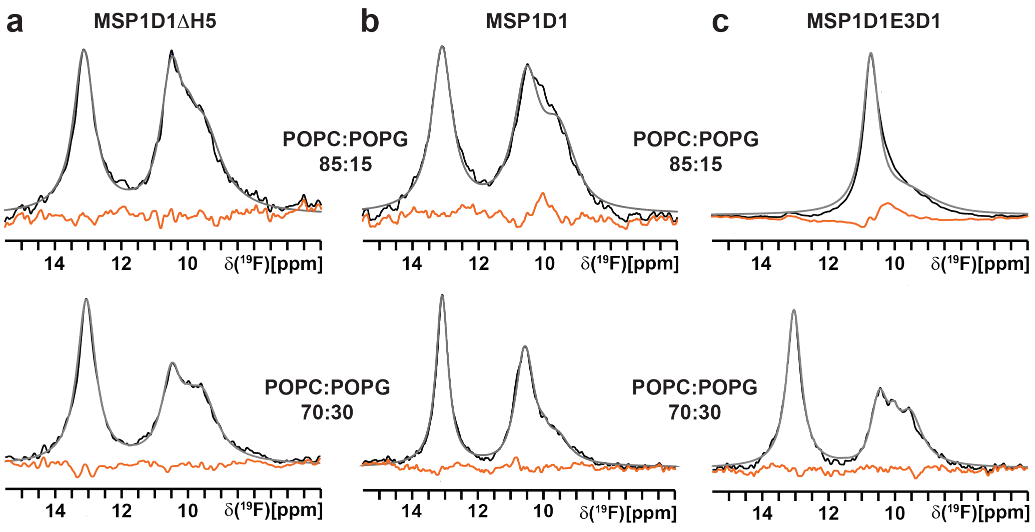


**Supplementary Fig. 7. Residual differences between the raw NMR data and summation of the individually fit components from Figure 3.** Experimental NMR data are shown in black, grey lines superimposed on the spectra are the total sums of the individual deconvolutions, and the orange lines are the calculated differences between the grey and black lines for $^{19}$F-NMR spectra with $A_{2A}$AR[A289C$^{TET}$] in nanodiscs formed from **a** MSP1D1ΔH5, **b** MSP1D1, and **c** MSP1E3D1 scaffold proteins.

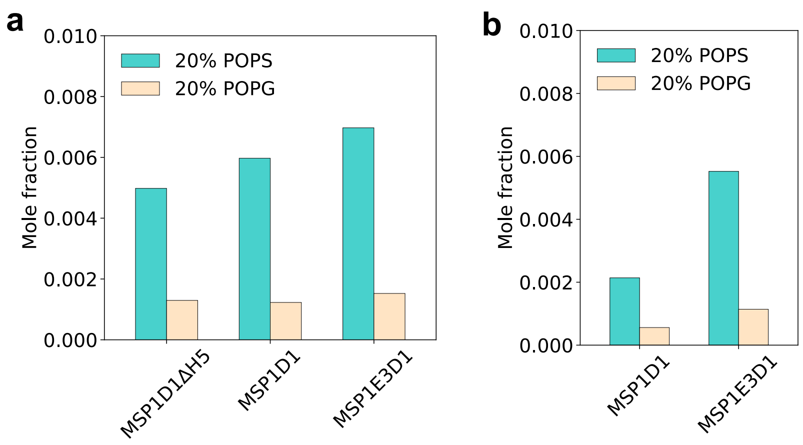


**Supplementary Fig. 8. $Na^+$ mediated anionic lipid pairs.** **a** Mole fraction of POPS or POPG lipids participating in $Na^+$ mediated pairs in three nanodiscs: MSP1D1ΔH5, MSP1D1, and MSP1E3D1. **b** Mole fraction of POPS or POPG lipids participating in $Na^+$ mediated pairs in MSP1D1 and MSP1E3D1 nanodiscs with $A_{2A}AR$ embedded. Simulations consisted of either POPC:POPS at a molar ratio of 80:20 or POPC:POPG at a molar ratio of 80:20.

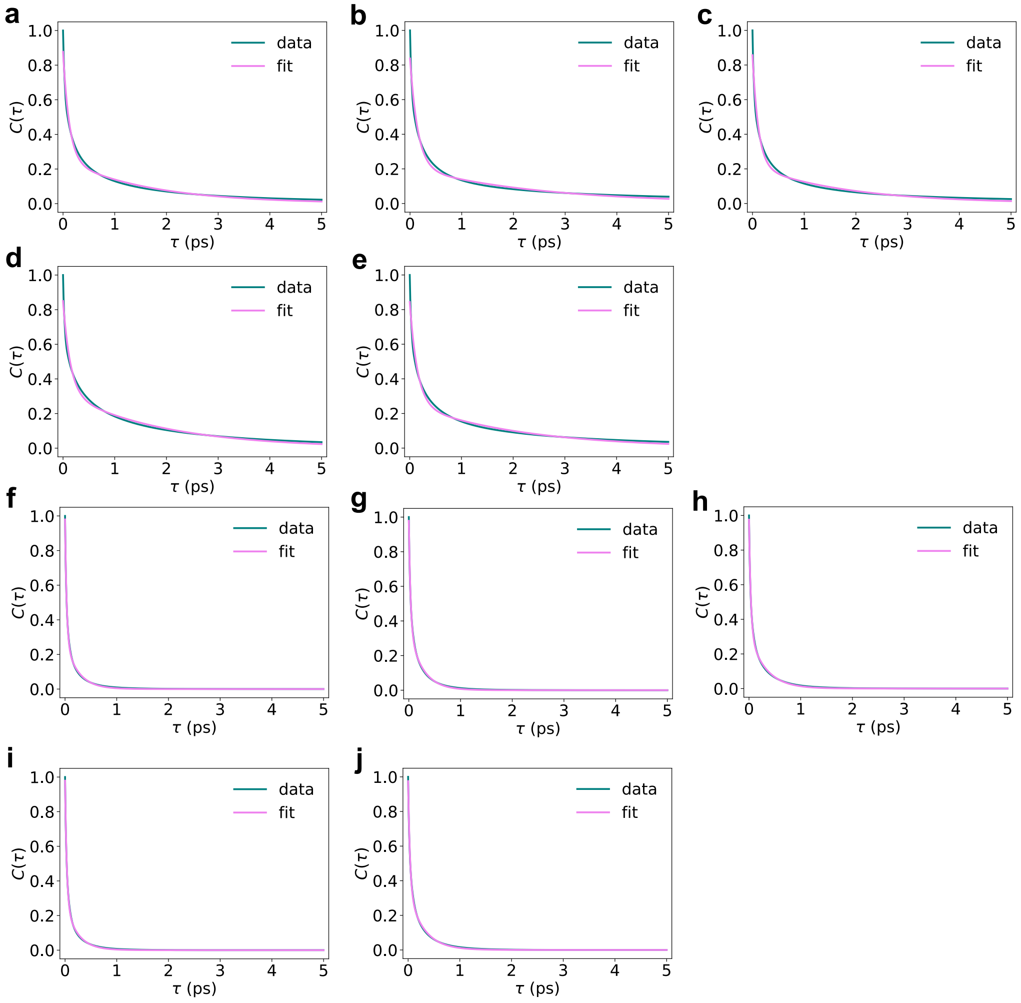


**Supplementary Fig. 9. Hydrogen-bond lifetime C(τ) curves. a-c** Lifetime curves calculated for a POPC:POPS molar ratio of 80:20 in nanodiscs formed from **a** MSP1D1ΔH5, **b** MSP1D1, and **c** MSP1E3D1. **d** and **e** Lifetime curves calculated for a POPC:POPS molar ratio of 80:20 in nanodiscs formed from **d** MSP1D1 and **e** MSP1E3D1, each containing $A_{2A}$AR embedded. **f-h** Lifetime curves calculated for a POPC:POPG molar ratio of 80:20 in nanodiscs formed from **f** MSP1D1ΔH5, **g** MSP1D1, and **h** MSP1E3D1. **i** and **j** Lifetime curves calculated for a POPC:POPG molar ratio of 80:20 in nanodiscs formed from **i** MSP1D1 and **j** MSP1E3D1, each containing $A_{2A}$AR embedded.

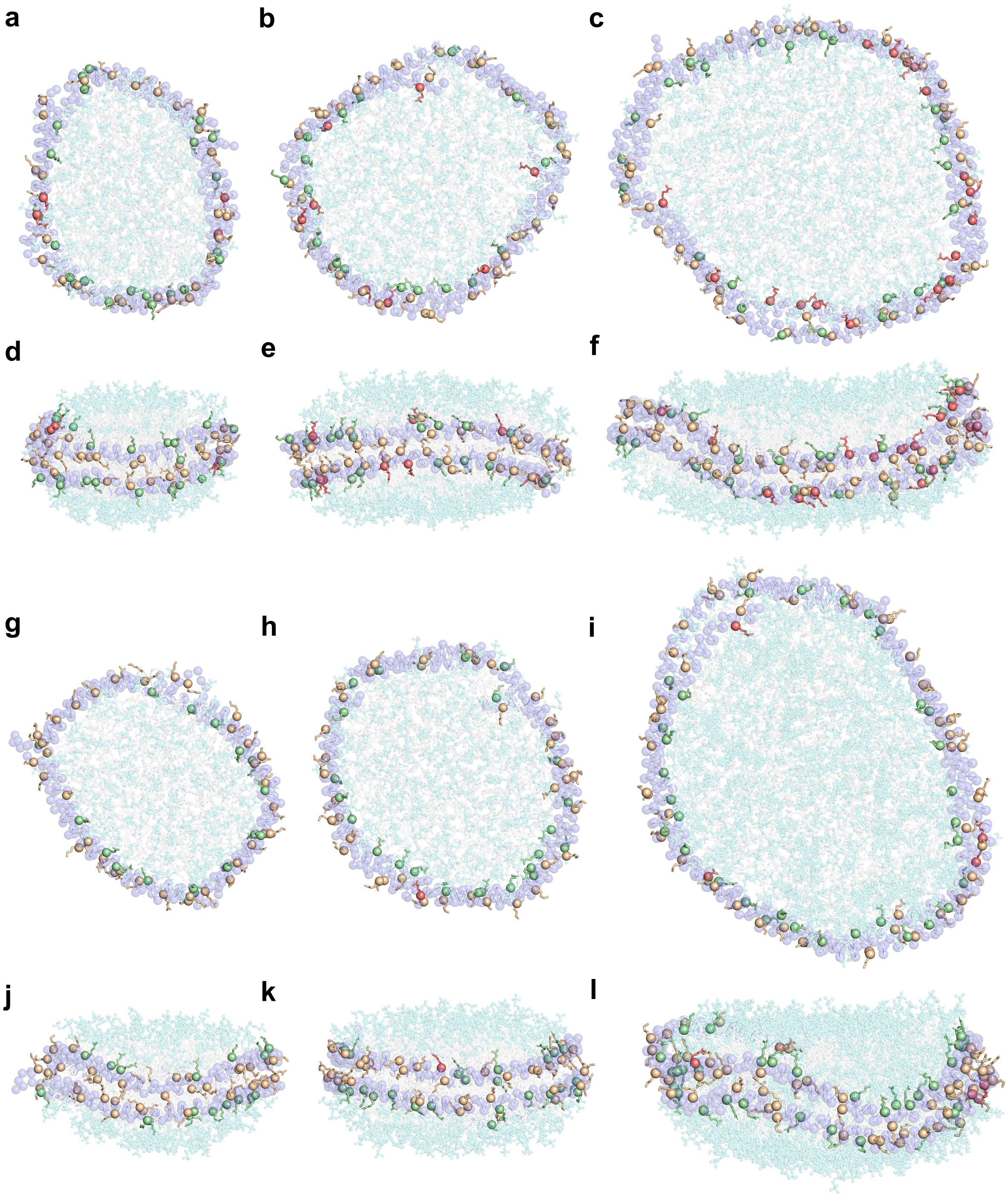

**Supplementary Fig. 10. Visualization of MSP Arg and Lys residues interacting with POPS or POPG in nanodiscs. a-c** Top-down views of MSP1D1ΔH5, MSP1D1, and MSP1E3D1 nanodiscs containing POPC:POPS membranes at a molar ratio of 80:20. **d-f** Corresponding side-on views of panels a-c. **g-i** Top-down views of MSP1D1ΔH5,

MSP1D1, and MSP1E3D1 nanodiscs containing POPC:POPG membranes at a molar ratio of 80:20. **j-l** Corresponding side-on views of panels g-i. The C$\alpha$ atoms of MSP residues are shown as spheres: red indicates residues exhibiting strong interactions with POPS or POPG lipids (≥40% of the simulation time), green indicates medium interactions (≥5%), and wheat indicates weak interactions (<5%), including zero interactions for Arg/Lys residues that do not contact lipids. Definitions of strong, medium, and weak interactions are provided in the Methods section. Non-positively charged residues are shown ase semi-transparent purple spheres. Lipid headgroups are shown as aquamarine transparent sticks, while lipid tails are shown as gray transparent sticks.

a

c

b

d

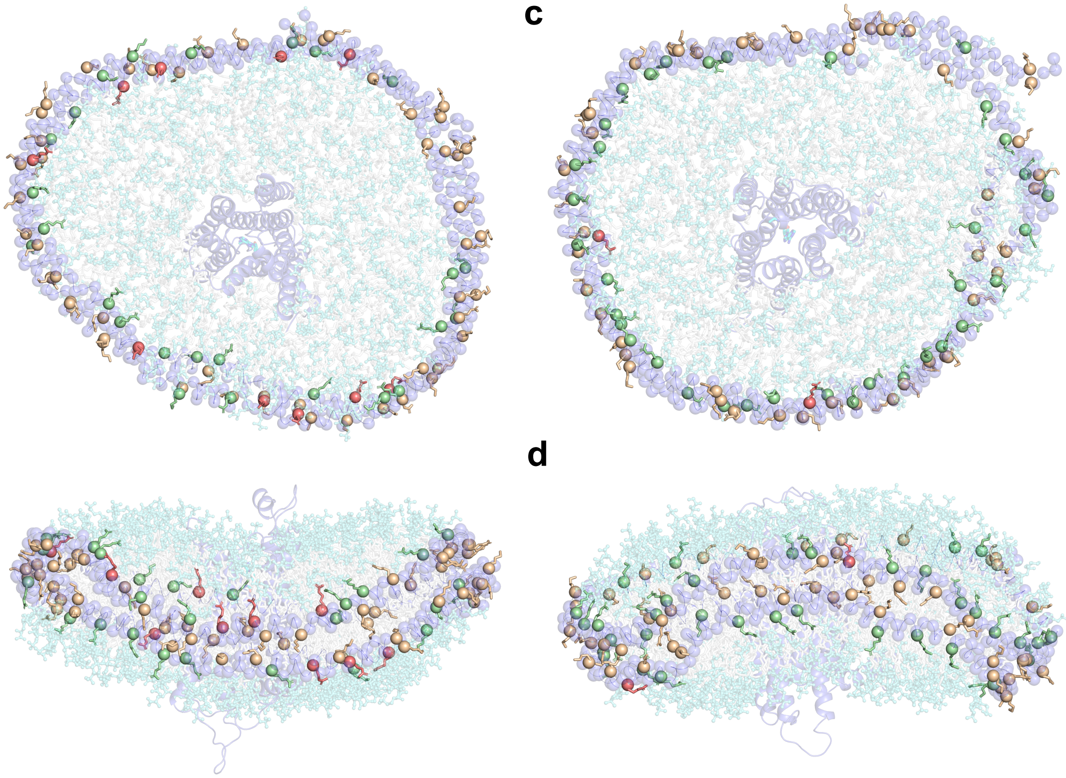

**Supplementary Fig. 11. Visualization of MSP Arg and Lys residues interacting with POPS or POPG in nanodiscs containing A$_{2A}$AR.** **a** and **b** Top-down and side-on views of MSP1E3D1 containing A$_{2A}$AR embedded in a membrane containing POPC:POPS at an 80:20 molar ratio. **c** and **d** Top-down and side-on views of MSP1E3D1 containing A$_{2A}$AR embedded in a membrane containing POPC:POPG at an 80:20 molar ratio. The C$\alpha$ atoms of MSP residues are shown as spheres: red indicates residues exhibiting strong interactions with POPS/POPG lipids (≥40% of the simulation time), green indicates medium interactions (≥5%), and wheat indicates weak interactions (<5%), including zero interactions for Arg/Lys residues that do not contact lipids. Definitions of strong, medium, and weak interactions are provided in the Methods section. Non-positively charged residues are shown as semi-transparent purple spheres. Lipid headgroups are shown as aquamarine transparent sticks, while lipid tails are shown as gray transparent sticks.

$A_{2A}$AR is shown as a transparent purple cartoon representation, and the ligand is shown as transparent sticks.

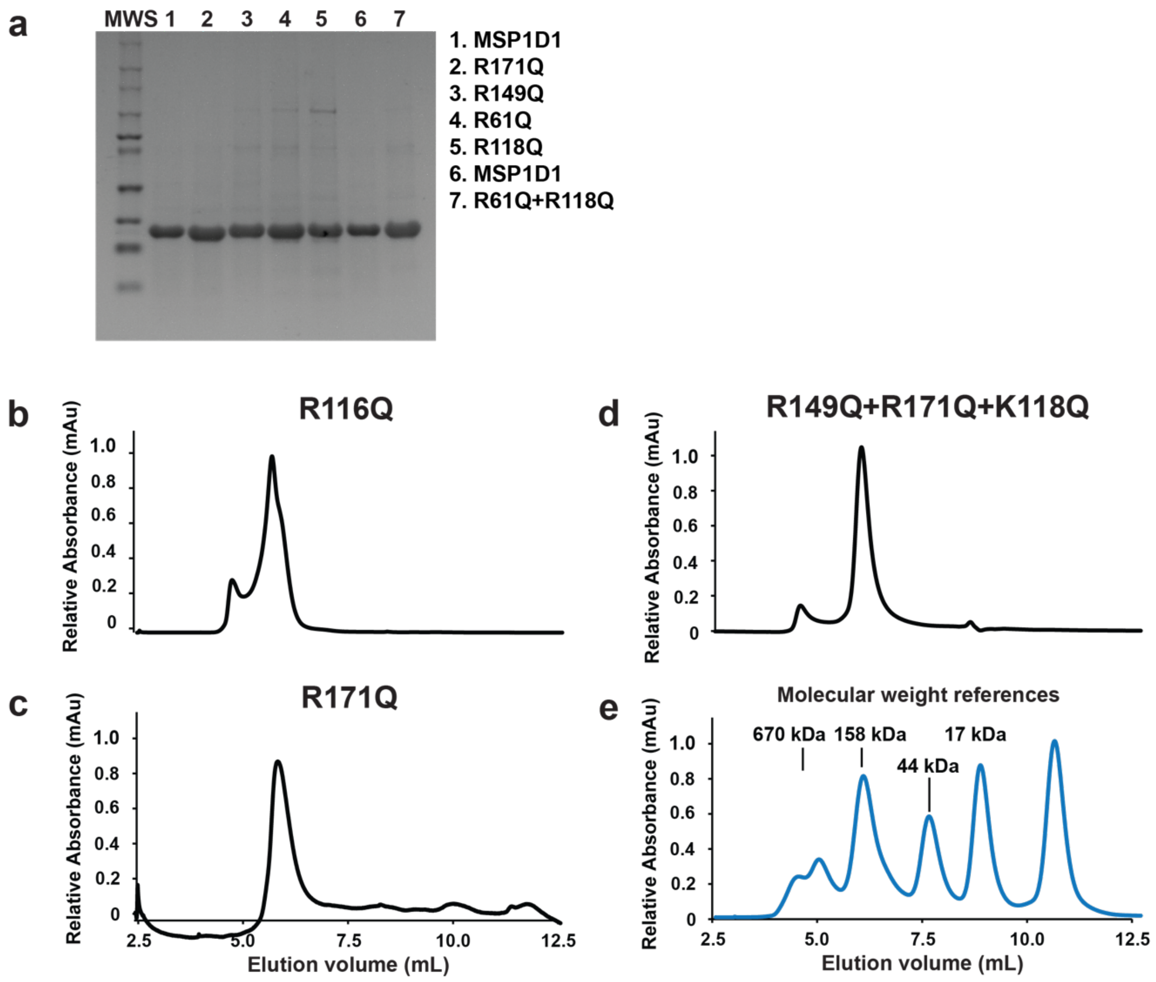


**Supplementary Fig. 12. Biochemical characterization of lipid nanodiscs formed from variant MSP1D1 proteins.** **a** SDS-PAGE gel of purified MSP1D1 and MSP1D1 variants, as annotated. The lane labeled "MWS" corresponds to molecular weight standards. **b-d** Analytical size exclusion (aSEC) chromatograms of lipid nanodiscs formed from variant MSP1D1 proteins. **e** Chromatogram of the molecular standards used to estimate the molecular weight of the nanodisc systems with annotated molecular weights shown.

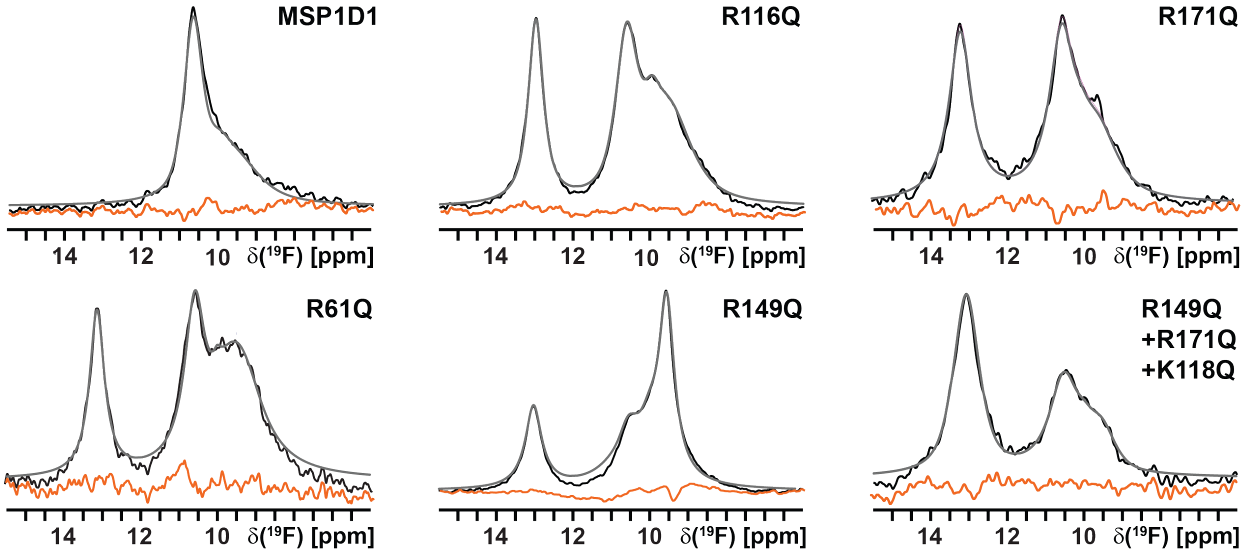


**Supplementary Fig. 13. Residual differences between the raw NMR data and summation of the individually fit components from Figure 6.** Experimental NMR data are shown in black, grey lines superimposed on the spectra are the total sums of the individual deconvolutions, and the orange lines are the calculated differences between the grey and black lines for $^{19}$F-NMR spectra with $A_{2A}$AR[A289C$^{TET}$] in nanodiscs containing POPC:POPS at a molar ratio 85:15 and formed from MSP1D1 and MSP1D1 variants.

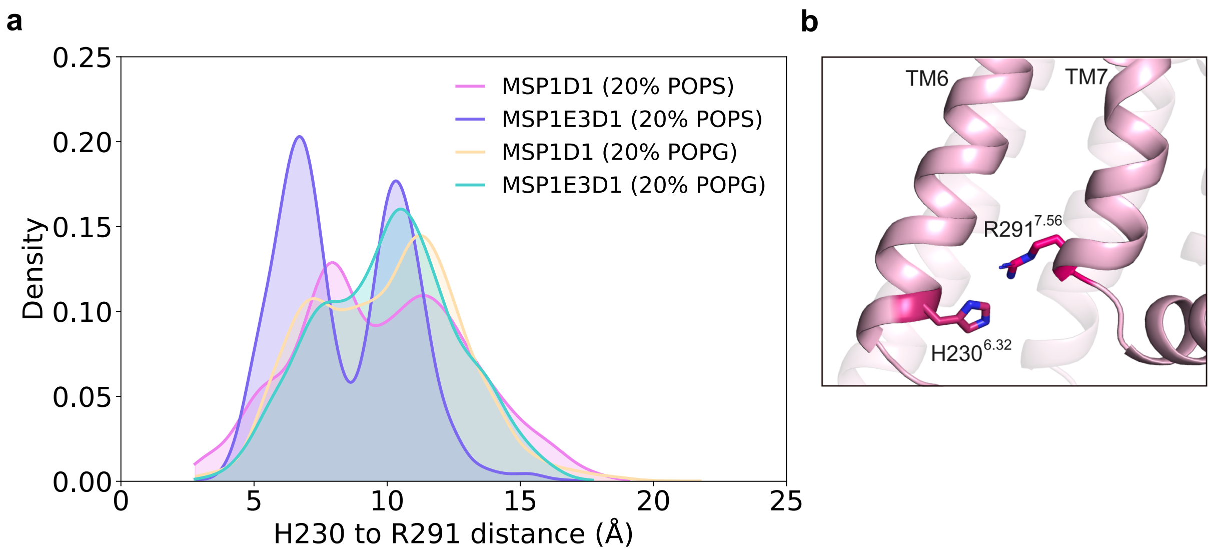


**Supplementary Fig. 14. Distance distributions between $A_{2A}$AR residues H230$^{6.32}$-R291$^{7.56}$ in different simulated nanodisc systems. a** Distance distributions between H230$^{6.32}$ and R291$^{7.56}$ of $A_{2A}$AR embedded in MSP1D1 and MSP1E3D1 nanodiscs containing either POPC:POPS at a molar ratio of 80:20 or POPC:POPG at a molar ratio of 80:20, as indicated. **b** Representative snapshot showing the positions of H230$^{6.32}$ and R291$^{7.56}$.

**Supplementary Table 1.** Simulated systems and conditions. D is the initial diameter of each MSP.

| Lipid composition | D/nm | MSP type | $A_{2A}AR$ embedded | Duration/μs |
|---|---|---|---|---|
| POPS:POPC (1:4) | 8.1 | MSP1D1ΔH5 | no | 5 |
| | 9.7 | MSP1D1 | no | 5 |
| | 12.1 | MSP1E3D1 | no | 5 |
| | 9.7 | MSP1D1 | yes | 5 |
| | 12.1 | MSP1E3D1 | yes | 5 |
| POPG:POPC (1:4) | 8.1 | MSP1D1ΔH5 | no | 5 |
| | 9.7 | MSP1D1 | no | 5 |
| | 12.1 | MSP1E3D1 | no | 5 |
| | 9.7 | MSP1D1 | yes | 5 |
| | 12.1 | MSP1E3D1 | yes | 5 |

**Supplementary Table 2. Hydrogen-bond lifetimes of POPS and POPG lipids.** The autocorrelation is extracted from the fluctuation of the hydrogen bond number and fitted by a bi-exponential decay. *Lifetimes were calculated using $A\tau_1 + B\tau_2$ after fitting. **Lifetimes were also estimated using the trapezoidal rule.

| Lipid composition | MSP type | $A_{2A}AR$ embedded | A | $\tau_1$ /ps | B | $\tau_2$ /ps | A+B | *Lifetime /ps | **Lifetime /ps |
|---|---|---|---|---|---|---|---|---|---|
| POPS:POPC (1:4) | MSP1D1ΔH5 | No | 0.62 | 0.12 | 0.25 | 1.66 | 0.88 | 0.50 | 0.48 |
| | MSP1D1 | No | 0.63 | 0.15 | 0.21 | 2.45 | 0.84 | 0.60 | 0.54 |
| | MSP1E3D1 | No | 0.64 | 0.11 | 0.22 | 1.82 | 0.86 | 0.47 | 0.45 |
| | MSP1D1 | Yes | 0.53 | 0.14 | 0.32 | 1.92 | 0.85 | 0.69 | 0.65 |
| | MSP1E3D1 | Yes | 0.59 | 0.15 | 0.25 | 2.13 | 0.84 | 0.63 | 0.58 |
| POPG:POPC (1:4) | MSP1D1ΔH5 | No | 0.70 | 0.03 | 0.28 | 0.25 | 0.98 | 0.09 | 0.10 |
| | MSP1D1 | No | 0.63 | 0.03 | 0.35 | 0.26 | 0.98 | 0.11 | 0.12 |
| | MSP1E3D1 | No | 0.32 | 0.32 | 0.65 | 0.03 | 0.98 | 0.12 | 0.13 |
| | MSP1D1 | Yes | 0.71 | 0.03 | 0.27 | 0.23 | 0.98 | 0.09 | 0.09 |
| | MSP1E3D1 | Yes | 0.63 | 0.04 | 0.34 | 0.30 | 0.98 | 0.12 | 0.13 |

**Supplementary Table 3. Interaction frequencies of Arg and Lys residues in the two MSP1D1ΔH5 chains with anionic lipids in a membrane containing POPC:POPS at a molar ratio of 80:20.**

| Chain A | | | Chain B | | |
|---|---|---|---|---|---|
| Resname | Resid_in_PDB | Interactions (%) | Resname | Resid_in_PDB | Interactions (%) |
| ARG | 177 | 69.92 | ARG | 188 | 72.76 |
| ARG | 173 | 53.34 | ARG | 177 | 70.63 |
| LYS | 239 | 28.77 | ARG | 173 | 25.26 |
| LYS | 107 | 21.64 | LYS | 195 | 23.31 |
| LYS | 59 | 19.48 | LYS | 206 | 21.31 |
| ARG | 188 | 16.20 | LYS | 96 | 18.60 |
| LYS | 195 | 11.93 | LYS | 118 | 18.42 |
| LYS | 206 | 10.89 | LYS | 107 | 17.00 |
| LYS | 88 | 9.78 | LYS | 239 | 12.55 |
| ARG | 151 | 9.64 | LYS | 88 | 12.20 |
| LYS | 118 | 9.51 | ARG | 151 | 10.31 |
| LYS | 96 | 7.89 | LYS | 238 | 9.89 |
| LYS | 106 | 5.33 | LYS | 106 | 6.95 |
| LYS | 238 | 1.31 | ARG | 171 | 3.89 |
| LYS | 77 | 0.96 | LYS | 77 | 2.31 |
| ARG | 61 | 0 | LYS | 59 | 0.09 |
| ARG | 83 | 0 | ARG | 61 | 0 |
| LYS | 94 | 0 | ARG | 83 | 0 |
| ARG | 116 | 0 | LYS | 94 | 0 |
| ARG | 149 | 0 | ARG | 116 | 0 |
| ARG | 153 | 0 | ARG | 149 | 0 |
| ARG | 160 | 0 | ARG | 153 | 0 |
| ARG | 171 | 0 | ARG | 160 | 0 |
| LYS | 182 | 0 | LYS | 182 | 0 |
| LYS | 208 | 0 | LYS | 208 | 0 |
| ARG | 215 | 0 | ARG | 215 | 0 |
| LYS | 226 | 0 | LYS | 226 | 0 |

**Supplementary Table 4. Interaction frequencies of Arg and Lys residues in the two MSP1D1 chains with anionic lipids in membranes containing POPC:POPS at a molar ratio of 80:20.**

| Chain A | | | Chain B | | |
|---|---|---|---|---|---|
| Resname | Resid_in_PDB | Interactions (%) | Resname | Resid_in_PDB | Interactions (%) |
| ARG | 116 | 65.52 | ARG | 149 | 60.28 |
| ARG | 61 | 48.79 | ARG | 61 | 52.83 |
| ARG | 171 | 41.50 | ARG | 123 | 50.41 |
| ARG | 160 | 36.95 | ARG | 171 | 48.03 |
| ARG | 123 | 32.77 | ARG | 116 | 46.72 |
| LYS | 94 | 29.99 | ARG | 153 | 36.93 |
| ARG | 149 | 26.62 | ARG | 160 | 30.44 |
| ARG | 153 | 19.24 | ARG | 215 | 23.66 |
| ARG | 131 | 16.84 | LYS | 208 | 23.15 |
| ARG | 83 | 12.40 | ARG | 131 | 19.97 |
| LYS | 106 | 9.31 | LYS | 59 | 14.24 |
| LYS | 226 | 5.44 | ARG | 83 | 11.09 |
| ARG | 215 | 4.47 | LYS | 182 | 9.51 |
| LYS | 77 | 4.44 | LYS | 94 | 7.80 |
| LYS | 208 | 3.75 | LYS | 77 | 4.91 |
| LYS | 88 | 2.20 | LYS | 106 | 2.24 |
| LYS | 182 | 1.24 | LYS | 226 | 0.89 |
| LYS | 59 | 0 | LYS | 88 | 0 |
| LYS | 96 | 0 | LYS | 96 | 0 |
| LYS | 107 | 0 | LYS | 107 | 0 |
| LYS | 118 | 0 | LYS | 118 | 0 |
| LYS | 133 | 0 | LYS | 133 | 0 |
| LYS | 140 | 0 | LYS | 140 | 0 |
| ARG | 151 | 0 | ARG | 151 | 0 |
| ARG | 173 | 0 | ARG | 173 | 0 |
| ARG | 177 | 0 | ARG | 177 | 0 |
| ARG | 188 | 0 | ARG | 188 | 0 |
| LYS | 195 | 0 | LYS | 195 | 0 |
| LYS | 206 | 0 | LYS | 206 | 0 |
| LYS | 238 | 0 | LYS | 238 | 0 |
| LYS | 239 | 0 | LYS | 239 | 0 |

**Supplementary Table 5. Interaction frequencies of Arg and Lys residues in the two MSP1D1 chains with anionic lipids in membrane containing POPC:POPS at a molar ratio of 80:20 and with $A_{2A}AR$ embedded.**

| Chain A | | | Chain B | | |
|---|---|---|---|---|---|
| Resname | Resid_in_PDB | Interactions (%) | Resname | Resid_in_PDB | Interactions (%) |
| ARG | 171 | 58.56 | ARG | 160 | 82.40 |
| ARG | 215 | 34.88 | ARG | 153 | 71.23 |
| ARG | 61 | 32.26 | ARG | 215 | 61.99 |
| ARG | 131 | 30.02 | ARG | 171 | 47.90 |
| LYS | 208 | 23.42 | ARG | 149 | 46.90 |
| ARG | 153 | 21.24 | LYS | 208 | 36.55 |
| ARG | 160 | 20.53 | LYS | 226 | 15.69 |
| LYS | 182 | 19.33 | ARG | 83 | 13.91 |
| LYS | 226 | 14.26 | ARG | 131 | 13.86 |
| ARG | 123 | 10.35 | LYS | 94 | 13.77 |
| ARG | 149 | 7.64 | LYS | 182 | 11.11 |
| LYS | 94 | 6.87 | ARG | 116 | 8.93 |
| ARG | 116 | 6.80 | LYS | 88 | 2.93 |
| LYS | 106 | 3.98 | LYS | 106 | 2.11 |
| ARG | 83 | 2.27 | ARG | 123 | 1.16 |
| LYS | 59 | 0.09 | LYS | 59 | 0 |
| LYS | 77 | 0 | ARG | 61 | 0 |
| LYS | 88 | 0 | LYS | 77 | 0 |
| LYS | 96 | 0 | LYS | 96 | 0 |
| LYS | 107 | 0 | LYS | 107 | 0 |
| LYS | 118 | 0 | LYS | 118 | 0 |
| LYS | 133 | 0 | LYS | 133 | 0 |
| LYS | 140 | 0 | LYS | 140 | 0 |
| ARG | 151 | 0 | ARG | 151 | 0 |
| ARG | 173 | 0 | ARG | 173 | 0 |
| ARG | 177 | 0 | ARG | 177 | 0 |
| ARG | 188 | 0 | ARG | 188 | 0 |
| LYS | 195 | 0 | LYS | 195 | 0 |
| LYS | 206 | 0 | LYS | 206 | 0 |
| LYS | 238 | 0 | LYS | 238 | 0 |
| LYS | 239 | 0 | LYS | 239 | 0 |

**Supplementary Table 6. Interaction frequencies of Arg and Lys residues in the two MSP1E3D1 chains with anionic lipids in membranes containing POPC and POPS at a molar ratio of 80:20.**

| Chain A | | | Chain B | | |
|---|---|---|---|---|---|
| Resname | Resid_in_PDB | Interactions (%) | Resname | Resid_in_PDB | Interactions (%) |
| ARG | 123 | 89.54 | ARG | 61 | 91.40 |
| ARG | 153 | 65.50 | ARG | 215 | 74.67 |
| ARG | 160 | 55.52 | ARG | 149 | 71.83 |
| ARG | 116 | 54.97 | ARG | 153 | 67.56 |
| ARG | 226 | 51.88 | ARG | 219 | 64.03 |
| ARG | 219 | 46.77 | ARG | 160 | 53.59 |
| ARG | 215 | 44.72 | ARG | 83 | 53.52 |
| ARG | 149 | 44.32 | ARG | 123 | 47.83 |
| ARG | 197 | 38.15 | ARG | 226 | 43.23 |
| ARG | 61 | 35.59 | ARG | 116 | 38.97 |
| ARG | 83 | 34.53 | ARG | 237 | 20.17 |
| ARG | 237 | 32.35 | ARG | 182 | 14.73 |
| LYS | 274 | 22.93 | LYS | 94 | 14.31 |
| ARG | 189 | 22.44 | LYS | 274 | 12.15 |
| ARG | 182 | 13.51 | ARG | 197 | 12.13 |
| LYS | 77 | 11.71 | ARG | 189 | 10.49 |
| ARG | 281 | 11.11 | ARG | 281 | 8.42 |
| ARG | 131 | 10.35 | ARG | 131 | 8.00 |
| LYS | 94 | 9.20 | LYS | 172 | 7.40 |
| LYS | 292 | 8.86 | LYS | 106 | 5.73 |
| LYS | 248 | 5.64 | LYS | 292 | 4.22 |
| LYS | 59 | 3.58 | LYS | 248 | 3.58 |
| LYS | 172 | 2.09 | LYS | 77 | 3.42 |
| LYS | 106 | 2.02 | LYS | 88 | 1.47 |
| LYS | 304 | 0.38 | LYS | 272 | 0.13 |
| LYS | 88 | 0 | LYS | 59 | 0 |
| LYS | 96 | 0 | LYS | 96 | 0 |
| LYS | 107 | 0 | LYS | 107 | 0 |
| LYS | 118 | 0 | LYS | 118 | 0 |
| LYS | 133 | 0 | LYS | 133 | 0 |
| LYS | 140 | 0 | LYS | 140 | 0 |
| ARG | 151 | 0 | ARG | 151 | 0 |
| LYS | 173 | 0 | LYS | 173 | 0 |

| | | | | | |
|---|---|---|---|---|---|
| LYS | 184 | 0 | LYS | 184 | 0 |
| LYS | 199 | 0 | LYS | 199 | 0 |
| LYS | 206 | 0 | LYS | 206 | 0 |
| ARG | 217 | 0 | ARG | 217 | 0 |
| ARG | 239 | 0 | ARG | 239 | 0 |
| ARG | 243 | 0 | ARG | 243 | 0 |
| ARG | 254 | 0 | ARG | 254 | 0 |
| LYS | 261 | 0 | LYS | 261 | 0 |
| LYS | 272 | 0 | LYS | 304 | 0 |
| LYS | 305 | 0 | LYS | 305 | 0 |

**Supplementary Table 7. Interaction frequencies of Arg and Lys residues in the two MSP1E3D1 chains with anionic lipids in membranes containing POPC:POPS at a molar ratio of 80:20 with $A_{2A}AR$ embedded.**

| Chain A | | | Chain B | | |
|---|---|---|---|---|---|
| Resname | Resid_in_PDB | Interactions (%) | Resname | Resid_in_PDB | Interactions (%) |
| ARG | 123 | 63.96 | ARG | 61 | 64.41 |
| ARG | 116 | 41.92 | ARG | 116 | 53.88 |
| ARG | 281 | 40.57 | ARG | 197 | 52.23 |
| ARG | 237 | 40.17 | ARG | 160 | 49.90 |
| ARG | 153 | 37.04 | ARG | 189 | 49.50 |
| ARG | 160 | 35.08 | ARG | 83 | 44.52 |
| LYS | 94 | 33.66 | ARG | 123 | 36.79 |
| ARG | 131 | 24.46 | ARG | 153 | 34.75 |
| ARG | 83 | 20.73 | LYS | 94 | 32.55 |
| LYS | 274 | 19.64 | ARG | 215 | 31.66 |
| ARG | 197 | 18.60 | LYS | 59 | 24.91 |
| ARG | 149 | 12.97 | ARG | 131 | 20.91 |
| ARG | 219 | 11.13 | LYS | 248 | 18.62 |
| LYS | 292 | 10.38 | ARG | 182 | 17.60 |
| ARG | 226 | 9.26 | ARG | 237 | 7.91 |
| LYS | 248 | 4.67 | LYS | 172 | 7.66 |
| LYS | 88 | 4.22 | ARG | 219 | 7.55 |
| ARG | 215 | 2.58 | LYS | 77 | 5.73 |
| LYS | 106 | 2.53 | LYS | 88 | 5.49 |
| LYS | 77 | 2.47 | ARG | 149 | 5.29 |
| ARG | 189 | 1.58 | LYS | 292 | 4.67 |
| ARG | 182 | 1.38 | LYS | 274 | 1.64 |
| ARG | 61 | 0.40 | LYS | 106 | 1.49 |
| LYS | 172 | 0.29 | ARG | 226 | 1.36 |
| LYS | 304 | 0.04 | ARG | 281 | 0.29 |
| LYS | 59 | 0 | LYS | 96 | 0 |
| LYS | 96 | 0 | LYS | 107 | 0 |
| LYS | 107 | 0 | LYS | 118 | 0 |
| LYS | 118 | 0 | LYS | 133 | 0 |
| LYS | 133 | 0 | LYS | 140 | 0 |
| LYS | 140 | 0 | ARG | 151 | 0 |
| ARG | 151 | 0 | LYS | 173 | 0 |
| LYS | 173 | 0 | LYS | 184 | 0 |
| LYS | 184 | 0 | LYS | 199 | 0 |

| | | | | | |
|---|---|---|---|---|---|
| LYS | 199 | 0 | LYS | 206 | 0 |
| LYS | 206 | 0 | ARG | 217 | 0 |
| ARG | 217 | 0 | ARG | 239 | 0 |
| ARG | 239 | 0 | ARG | 243 | 0 |
| ARG | 243 | 0 | ARG | 254 | 0 |
| ARG | 254 | 0 | LYS | 261 | 0 |
| LYS | 261 | 0 | LYS | 272 | 0 |
| LYS | 272 | 0 | LYS | 304 | 0 |
| LYS | 305 | 0 | LYS | 305 | 0 |

**Supplementary Table 8. Interaction frequencies of Arg and Lys residues in the two MSP1D1ΔH5 chains with anionic lipids in membranes containing POPC:POPG at a molar ratio of 80:20.**

| Chain A | | | Chain B | | |
|---|---|---|---|---|---|
| Resname | Resid_in_PDB | Interactions (%) | Resname | Resid_in_PDB | Interactions (%) |
| ARG | 188 | 38.55 | ARG | 177 | 24.35 |
| ARG | 177 | 34.64 | ARG | 173 | 18.04 |
| ARG | 173 | 28.70 | LYS | 118 | 17.33 |
| ARG | 151 | 22.24 | ARG | 188 | 15.11 |
| LYS | 118 | 14.66 | ARG | 151 | 14.09 |
| LYS | 107 | 11.64 | LYS | 107 | 12.55 |
| LYS | 96 | 8.91 | LYS | 195 | 9.69 |
| LYS | 239 | 4.38 | LYS | 96 | 7.00 |
| LYS | 206 | 3.33 | LYS | 239 | 6.55 |
| LYS | 195 | 2.20 | LYS | 206 | 6.44 |
| ARG | 61 | 1.64 | ARG | 61 | 2.49 |
| ARG | 215 | 0.33 | LYS | 77 | 0.13 |
| LYS | 77 | 0.22 | LYS | 106 | 0.04 |
| LYS | 88 | 0.18 | LYS | 59 | 0.02 |
| LYS | 59 | 0.07 | ARG | 83 | 0 |
| LYS | 106 | 0.04 | LYS | 88 | 0 |
| ARG | 83 | 0 | LYS | 94 | 0 |
| LYS | 94 | 0 | ARG | 116 | 0 |
| ARG | 116 | 0 | ARG | 149 | 0 |
| ARG | 149 | 0 | ARG | 153 | 0 |
| ARG | 153 | 0 | ARG | 160 | 0 |
| ARG | 160 | 0 | ARG | 171 | 0 |
| ARG | 171 | 0 | LYS | 182 | 0 |
| LYS | 182 | 0 | LYS | 208 | 0 |
| LYS | 208 | 0 | ARG | 215 | 0 |
| LYS | 226 | 0 | LYS | 226 | 0 |
| LYS | 238 | 0 | LYS | 238 | 0 |

**Supplementary Table 9. Interaction frequencies of Arg and Lys residues in the two MSP1D1 chains with anionic lipids in membranes containing POPC:POPG at a molar ratio of 80:20.**

| Chain A | | | Chain B | | |
|---|---|---|---|---|---|
| Resname | Resid_in_PDB | Interactions (%) | Resname | Resid_in_PDB | Interactions (%) |
| ARG | 116 | 44.95 | ARG | 131 | 37.70 |
| ARG | 153 | 18.42 | ARG | 153 | 36.30 |
| ARG | 83 | 17.53 | ARG | 171 | 30.13 |
| ARG | 131 | 14.66 | ARG | 149 | 29.50 |
| ARG | 123 | 14.13 | ARG | 83 | 26.48 |
| ARG | 160 | 12.64 | ARG | 123 | 25.59 |
| ARG | 215 | 10.22 | ARG | 160 | 21.66 |
| LYS | 206 | 8.93 | LYS | 94 | 21.33 |
| ARG | 61 | 8.46 | ARG | 215 | 15.29 |
| ARG | 149 | 7.58 | LYS | 208 | 13.82 |
| LYS | 226 | 4.42 | LYS | 106 | 13.49 |
| ARG | 171 | 3.71 | LYS | 59 | 13.00 |
| LYS | 94 | 2.38 | LYS | 182 | 9.26 |
| LYS | 182 | 2.38 | ARG | 116 | 7.80 |
| LYS | 208 | 1.89 | LYS | 226 | 3.42 |
| LYS | 195 | 0.98 | ARG | 61 | 1.58 |
| LYS | 59 | 0.89 | LYS | 206 | 0.33 |
| ARG | 188 | 0.53 | LYS | 239 | 0.09 |
| LYS | 239 | 0.36 | LYS | 77 | 0.02 |
| LYS | 77 | 0 | LYS | 88 | 0.02 |
| LYS | 88 | 0 | LYS | 96 | 0 |
| LYS | 96 | 0 | LYS | 107 | 0 |
| LYS | 106 | 0 | LYS | 118 | 0 |
| LYS | 107 | 0 | LYS | 133 | 0 |
| LYS | 118 | 0 | LYS | 140 | 0 |
| LYS | 133 | 0 | ARG | 151 | 0 |
| LYS | 140 | 0 | ARG | 173 | 0 |
| ARG | 151 | 0 | ARG | 177 | 0 |
| ARG | 173 | 0 | ARG | 188 | 0 |
| ARG | 177 | 0 | LYS | 195 | 0 |
| LYS | 238 | 0 | LYS | 238 | 0 |

**Supplementary Table 10. Interaction frequencies of Arg and Lys residues in the two MSP1D1 chains with anionic lipids in membranes containing POPC:POPG at a molar ratio of 80:20 and with $A_{2A}AR$ embedded.**

| Chain A | | | Chain B | | |
|---|---|---|---|---|---|
| Resname | Resid_in_PDB | Interactions (%) | Resname | Resid_in_PDB | Interactions (%) |
| ARG | 131 | 64.50 | ARG | 160 | 73.32 |
| ARG | 153 | 42.55 | ARG | 171 | 36.41 |
| ARG | 83 | 39.41 | ARG | 153 | 14.73 |
| ARG | 149 | 28.59 | LYS | 182 | 13.40 |
| LYS | 226 | 16.71 | ARG | 131 | 11.31 |
| LYS | 94 | 12.55 | LYS | 208 | 7.22 |
| ARG | 171 | 7.89 | LYS | 94 | 6.60 |
| ARG | 215 | 6.78 | ARG | 83 | 5.40 |
| ARG | 123 | 6.24 | ARG | 123 | 5.18 |
| ARG | 61 | 4.18 | ARG | 116 | 4.62 |
| ARG | 160 | 3.04 | ARG | 215 | 3.18 |
| LYS | 208 | 2.36 | ARG | 61 | 3.04 |
| LYS | 182 | 1.69 | ARG | 149 | 1.29 |
| LYS | 106 | 1.38 | LYS | 226 | 0.42 |
| ARG | 116 | 0.62 | LYS | 106 | 0.02 |
| LYS | 88 | 0.02 | LYS | 59 | 0 |
| LYS | 59 | 0 | LYS | 77 | 0 |
| LYS | 77 | 0 | LYS | 88 | 0 |
| LYS | 96 | 0 | LYS | 96 | 0 |
| LYS | 107 | 0 | LYS | 107 | 0 |
| LYS | 118 | 0 | LYS | 118 | 0 |
| LYS | 133 | 0 | LYS | 133 | 0 |
| LYS | 140 | 0 | LYS | 140 | 0 |
| ARG | 151 | 0 | ARG | 151 | 0 |
| ARG | 173 | 0 | ARG | 173 | 0 |
| ARG | 177 | 0 | ARG | 177 | 0 |
| ARG | 188 | 0 | ARG | 188 | 0 |
| LYS | 195 | 0 | LYS | 195 | 0 |
| LYS | 206 | 0 | LYS | 206 | 0 |
| LYS | 238 | 0 | LYS | 238 | 0 |
| LYS | 239 | 0 | LYS | 239 | 0 |

**Supplementary Table 11. Interaction frequencies of Arg and Lys residues in the two MSP1E3D1 chains with anionic lipids in membranes containing POPC:POPG at a molar ratio of 80:20.**

| belt_1 | | | belt_2 | | |
|---|---|---|---|---|---|
| Resname | Resid_in_PDB | interactions (%) | Resname | Resid_in_PDB | interactions (%) |
| ARG | 131 | 53.32 | ARG | 123 | 73.29 |
| ARG | 61 | 50.26 | ARG | 116 | 44.23 |
| ARG | 160 | 36.08 | ARG | 131 | 37.35 |
| ARG | 83 | 32.06 | ARG | 219 | 31.28 |
| ARG | 182 | 27.15 | ARG | 83 | 27.79 |
| ARG | 281 | 25.91 | ARG | 189 | 27.13 |
| ARG | 226 | 24.59 | ARG | 61 | 24.04 |
| ARG | 219 | 22.91 | ARG | 237 | 23.13 |
| ARG | 189 | 20.97 | ARG | 182 | 23.08 |
| LYS | 274 | 19.20 | ARG | 215 | 20.44 |
| ARG | 123 | 16.51 | ARG | 226 | 17.86 |
| ARG | 149 | 13.11 | ARG | 197 | 15.66 |
| LYS | 292 | 12.73 | ARG | 160 | 12.55 |
| ARG | 215 | 11.95 | ARG | 153 | 10.13 |
| ARG | 153 | 10.89 | ARG | 281 | 8.20 |
| LYS | 248 | 8.04 | LYS | 248 | 7.62 |
| ARG | 197 | 7.93 | ARG | 149 | 4.78 |
| ARG | 116 | 7.60 | LYS | 292 | 4.47 |
| ARG | 237 | 7.04 | LYS | 274 | 2.31 |
| LYS | 94 | 5.49 | LYS | 94 | 1.36 |
| LYS | 59 | 0.62 | LYS | 106 | 0.22 |
| LYS | 172 | 0.18 | LYS | 172 | 0.09 |
| LYS | 88 | 0.02 | LYS | 77 | 0.02 |
| LYS | 106 | 0.02 | LYS | 59 | 0 |
| LYS | 77 | 0 | LYS | 88 | 0 |
| LYS | 96 | 0 | LYS | 96 | 0 |
| LYS | 107 | 0 | LYS | 107 | 0 |
| LYS | 118 | 0 | LYS | 118 | 0 |
| LYS | 133 | 0 | LYS | 133 | 0 |
| LYS | 140 | 0 | LYS | 140 | 0 |
| ARG | 151 | 0 | ARG | 151 | 0 |
| LYS | 173 | 0 | LYS | 173 | 0 |
| LYS | 184 | 0 | LYS | 184 | 0 |
| LYS | 199 | 0 | LYS | 199 | 0 |

| | | | | | |
|---|---|---|---|---|---|
| LYS | 206 | 0 | LYS | 206 | 0 |
| ARG | 217 | 0 | ARG | 217 | 0 |
| ARG | 239 | 0 | ARG | 239 | 0 |
| ARG | 243 | 0 | ARG | 243 | 0 |
| ARG | 254 | 0 | ARG | 254 | 0 |
| LYS | 261 | 0 | LYS | 261 | 0 |
| LYS | 272 | 0 | LYS | 272 | 0 |
| LYS | 304 | 0 | LYS | 304 | 0 |
| LYS | 305 | 0 | LYS | 305 | 0 |

**Supplementary Table 12. Interaction frequencies of Arg and Lys residues in the two MSP1E3D1 chains with anionic lipids in membranes containing POPC:POPG at a molar ratio of 80:20 and with $A_{2A}AR$ embedded.**

| Chain A | | | Chain B | | |
|---|---|---|---|---|---|
| Resname | Resid_in_PDB | Interactions (%) | Resname | Resid_in_PDB | Interactions (%) |
| ARG | 197 | 55.72 | ARG | 189 | 42.50 |
| ARG | 61 | 39.92 | ARG | 237 | 31.39 |
| ARG | 189 | 26.91 | ARG | 149 | 29.13 |
| ARG | 83 | 26.77 | ARG | 182 | 26.37 |
| ARG | 116 | 24.08 | ARG | 153 | 21.13 |
| ARG | 219 | 24.02 | ARG | 131 | 19.44 |
| ARG | 160 | 19.26 | ARG | 281 | 19.33 |
| ARG | 182 | 18.95 | ARG | 254 | 18.84 |
| ARG | 153 | 18.24 | ARG | 61 | 18.55 |
| LYS | 94 | 14.77 | ARG | 116 | 18.31 |
| ARG | 149 | 14.53 | ARG | 160 | 16.57 |
| ARG | 226 | 13.44 | ARG | 219 | 16.53 |
| ARG | 237 | 10.53 | ARG | 226 | 14.57 |
| LYS | 292 | 7.98 | ARG | 123 | 14.06 |
| ARG | 131 | 7.24 | ARG | 83 | 13.26 |
| LYS | 248 | 7.24 | LYS | 248 | 12.35 |
| ARG | 123 | 7.09 | ARG | 197 | 8.00 |
| ARG | 215 | 6.13 | ARG | 215 | 6.78 |
| ARG | 281 | 3.60 | LYS | 274 | 6.78 |
| LYS | 274 | 3.20 | LYS | 94 | 5.55 |
| LYS | 88 | 0.18 | LYS | 292 | 4.49 |
| LYS | 172 | 0.07 | LYS | 261 | 3.62 |
| LYS | 304 | 0.02 | LYS | 172 | 3.04 |
| LYS | 59 | 0 | LYS | 272 | 1.44 |
| LYS | 77 | 0 | LYS | 106 | 0.56 |
| LYS | 96 | 0 | LYS | 304 | 0.27 |
| LYS | 106 | 0 | LYS | 59 | 0.09 |
| LYS | 107 | 0 | LYS | 88 | 0.02 |
| LYS | 118 | 0 | LYS | 77 | 0 |
| LYS | 133 | 0 | LYS | 96 | 0 |
| LYS | 140 | 0 | LYS | 107 | 0 |
| ARG | 151 | 0 | LYS | 118 | 0 |
| LYS | 173 | 0 | LYS | 133 | 0 |
| LYS | 184 | 0 | LYS | 140 | 0 |

| | | | | | |
|---|---|---|---|---|---|
| LYS | 199 | 0 | ARG | 151 | 0 |
| LYS | 206 | 0 | LYS | 173 | 0 |
| ARG | 217 | 0 | LYS | 184 | 0 |
| ARG | 239 | 0 | LYS | 199 | 0 |
| ARG | 243 | 0 | LYS | 206 | 0 |
| ARG | 254 | 0 | ARG | 217 | 0 |
| LYS | 261 | 0 | ARG | 239 | 0 |
| LYS | 272 | 0 | ARG | 243 | 0 |
| LYS | 305 | 0 | LYS | 305 | 0 |

**Supplementary Table 13. Interaction frequencies of Arg and Lys residues of $A_{2A}AR$ with anionic lipids in MSP1D1 nanodiscs containing POPC:POPS at a molar ratio of 80:20.**

| Resname | Resid | Interactions (%) |
|---|---|---|
| ARG | 296 | 98.76 |
| ARG | 300 | 98.22 |
| ARG | 199 | 96.18 |
| ARG | 304 | 90.22 |
| ARG | 293 | 89.56 |
| ARG | 206 | 78.76 |
| ARG | 291 | 77.63 |
| ARG | 205 | 57.81 |
| ARG | 120 | 40.55 |
| ARG | 107 | 35.48 |
| ARG | 102 | 25.42 |
| ARG | 220 | 11.66 |
| ARG | 222 | 10.53 |
| LYS | 122 | 8.40 |
| LYS | 233 | 3.95 |
| LYS | 227 | 1.09 |
| LYS | 209 | 0.13 |
| LYS | 301 | 0.07 |
| ARG | 111 | 0.00 |
| LYS | 150 | 0.00 |
| LYS | 153 | 0.00 |

**Supplementary Table 14. Interaction frequencies of Arg and Lys residues of $A_{2A}$AR with anionic lipids in MSP1E3D1 nanodiscs containing POPC:POPS lipids at a molar ratio of 80:20.**

| Resname | Resid | Interactions (%) |
|---|---|---|
| ARG | 206 | 86.80 |
| ARG | 199 | 86.71 |
| ARG | 300 | 83.74 |
| ARG | 296 | 80.32 |
| ARG | 107 | 73.45 |
| ARG | 293 | 67.74 |
| ARG | 205 | 62.08 |
| ARG | 220 | 54.30 |
| LYS | 233 | 53.45 |
| ARG | 120 | 44.48 |
| ARG | 111 | 41.70 |
| ARG | 222 | 35.84 |
| LYS | 122 | 15.84 |
| ARG | 304 | 12.93 |
| LYS | 209 | 7.98 |
| LYS | 227 | 6.71 |
| LYS | 150 | 2.18 |
| LYS | 301 | 0.20 |
| LYS | 153 | 0.07 |
| ARG | 102 | 0.00 |
| ARG | 291 | 0.00 |

**Supplementary Table 15. Interaction frequencies of Arg and Lys residues of $A_{2A}AR$ with anionic lipids in MSP1D1 nanodiscs containing POPC:POPG lipids at a molar ratio of 80:20.**

| Resname | Resid | Interactions (%) |
|---|---|---|
| ARG | 199 | 81.72 |
| ARG | 107 | 74.29 |
| ARG | 304 | 64.16 |
| LYS | 233 | 47.01 |
| ARG | 206 | 32.90 |
| ARG | 291 | 21.97 |
| ARG | 120 | 20.20 |
| ARG | 205 | 18.48 |
| LYS | 122 | 12.69 |
| ARG | 300 | 12.29 |
| ARG | 111 | 8.80 |
| ARG | 296 | 7.55 |
| ARG | 222 | 6.55 |
| ARG | 293 | 4.04 |
| LYS | 209 | 1.49 |
| LYS | 227 | 1.33 |
| ARG | 102 | 0.73 |
| LYS | 150 | 0.60 |
| LYS | 301 | 0.42 |
| ARG | 220 | 0.36 |
| LYS | 153 | 0.04 |

**Supplementary Table 16. Interaction frequencies of Arg and Lys residues of $A_{2A}$AR with anionic lipids in MSP1E3D1 nanodiscs containing POPC:POPG lipids at a molar ratio of 80:20.**

| Resname | Resid | Interactions (%) |
|---|---|---|
| ARG | 199 | 85.78 |
| ARG | 107 | 63.19 |
| ARG | 206 | 61.19 |
| ARG | 291 | 47.32 |
| ARG | 296 | 39.01 |
| LYS | 122 | 34.99 |
| LYS | 233 | 33.08 |
| ARG | 293 | 12.64 |
| ARG | 205 | 10.73 |
| ARG | 300 | 10.24 |
| ARG | 222 | 7.33 |
| ARG | 120 | 7.18 |
| ARG | 111 | 6.47 |
| ARG | 304 | 3.33 |
| ARG | 220 | 1.07 |
| LYS | 150 | 0.67 |
| ARG | 102 | 0.56 |
| LYS | 227 | 0.56 |
| LYS | 209 | 0.53 |
| LYS | 153 | 0.00 |
| LYS | 301 | 0.00 |

**Supplementary Table 17. Mean principal axis lengths of the simulated MSP nanodiscs along the major, minor (second), and third ("thickness") axes.** The aspect ratio was computed as the ratio of the major to the minor axis. Values in parentheses are the standard deviation. Values are in nm.

| Lipid composition | MSP type | $A_{2A}AR$ embedded | Major axis | Minor axis | Third axis | Aspect ratio |
|---|---|---|---|---|---|---|
| POPS:POPC (1:4) | MSP1D1ΔH5 | no | 9.29 (0.42) | 8.13 (0.42) | 2.99 (0.28) | 1.15 (0.09) |
| | MSP1D1 | no | 10.36 (0.35) | 9.09 (0.32) | 3.09 (0.38) | 1.14 (0.06) |
| | MSP1E3D1 | no | 14.44 (1.32) | 10.94 (0.48) | 4.36 (0.44) | 1.32 (0.15) |
| | MSP1D1 | yes | 10.69 (1.89) | 9.74 (0.48) | 2.77 (0.38) | 1.10 (0.19) |
| | MSP1E3D1 | yes | 13.62 (0.62) | 11.44 (0.60) | 4.14 (0.30) | 1.20 (0.11) |
| POPG:POPC (1:4) | MSP1D1ΔH5 | no | 9.75 (1.66) | 8.01 (0.48) | 3.01 (0.29) | 1.22 (0.20) |
| | MSP1D1 | no | 10.51 (0.35) | 9.11 (0.32) | 3.08 (0.50) | 1.16 (0.07) |
| | MSP1E3D1 | no | 13.80 (0.50) | 11.50 (0.41) | 3.77 (0.34) | 1.20 (0.08) |
| | MSP1D1 | yes | 10.49 (1.67) | 9.40 (0.35) | 2.74 (0.30) | 1.12 (0.17) |
| | MSP1E3D1 | yes | 13.94 (0.44) | 10.59 (0.42) | 4.85 (0.50) | 1.32 (0.08) |

**Supplementary Table 18. Relative populations of the $A_{2A}$AR conformational states observed in $^{19}$F-NMR spectra.** The relative populations of the $A_{2A}$AR conformational states as observed in the $^{19}$F-NMR spectra are tabulated below. The value reported for each conformation is a ratio of the integrated area of the specific deconvoluted peak to the total integral of all signals from 6 ppm to 15.5 ppm, excluding any signals from free TET.

| Ligand | Ligand efficacy | MSP | Composition | Relative populations | | | | |
|---|---|---|---|---|---|---|---|---|
| | | | | P4 | P3 | P2 | P1 | P5 |
| NECA | Full agonist | MSP1D1ΔH5 | POPC(100) | 0.32 | | 0.46 | 0.14 | 0.05 |
| NECA | Full agonist | MSP1D1ΔH5 | POPC:POPS (85:15) | 0.32 | | 0.35 | 0.26 | 0.06 |
| NECA | Full agonist | MSP1D1ΔH5 | POPC:POPS (78:22) | 0.33 | | 0.30 | 0.54 | 0.009 |
| NECA | Full agonist | MSP1D1ΔH5 | POPC:POPS (70:30) | 0.48 | | 0.33 | 0.13 | 0.04 |
| NECA | Full agonist | MSP1D1ΔH5 | POPC:POPG (85:15) | 0.37 | | 0.26 | 0.30 | 0.05 |
| NECA | Full agonist | MSP1D1ΔH5 | POPC:POPG (70:30) | 0.45 | | 0.23 | 0.28 | 0.02 |
| ZM241385 | Inverse agonist | MSP1D1ΔH5 | POPC:POPS (70:30) | | 0.78 | | 0.21 | |
| NECA | Full agonist | MSP1D1 | POPC(100) | | 0.80 | | 0.12 | 0.07 |
| NECA | Full agonist | MSP1D1 | POPC:POPS (85:15) | | 0.52 | | 0.31 | 0.15 |
| NECA | Full agonist | MSP1D1 | POPC:POPS (78:22) | 0.44 | | 0.31 | 0.15 | 0.07 |
| NECA | Full agonist | MSP1D1 | POPC:POPS (70:30) | 0.43 | | 0.41 | 0.08 | 0.07 |
| NECA | Full agonist | MSP1D1 | POPC:POPG (85:15) | 0.37 | | 0.35 | 0.19 | 0.07 |
| NECA | Full agonist | MSP1D1 | POPC:POPG (70:30) | 0.44 | | 0.40 | 0.12 | 0.03 |
| ZM241385 | Inverse agonist | MSP1D1 | POPC:POPS (70:30) | | 0.68 | | 0.31 | |
| NECA | Full agonist | MSP1E3D1 | POPC(100) | | | | | |
| NECA | Full agonist | MSP1E3D1 | POPC:POPS (85:15) | | | 0.51 | 0.39 | 0.09 |
| NECA | Full agonist | MSP1E3D1 | POPC:POPS (78:22) | | 0.63 | | 0.21 | 0.14 |
| NECA | Full agonist | MSP1E3D1 | POPC:POPS (70:30) | 0.43 | | 0.34 | 0.19 | 0.03 |
| NECA | Full agonist | MSP1E3D1 | POPC:POPG (85:15) | | 0.64 | | 0.16 | 0.18 |

| | | | | | | | | |
|---|---|---|---|---|---|---|---|---|
| NECA | Full agonist | MSP1E3D1 | POPC:POPG (70:30) | **0.48** | | **0.18** | **0.26** | **0.06** |
| ZM241385 | Inverse agonist | MSP1E3D1 | POPC:POPS (70:30) | | **0.79** | | **0.20** | |

**Supplementary Table 19. Relative populations of the $A_{2A}$AR conformational states observed in $^{19}$F-NMR spectra with nanodiscs formed from MSP1D1 variants.** The relative populations of the $A_{2A}$AR conformational states as observed in the $^{19}$F-NMR spectra are tabulated below. The value reported for each conformation is a ratio of the integrated area of the specific deconvoluted peak to the total integral of all signals from 6 ppm to 15.5 ppm, excluding any signals from free TET.

| Ligand | Ligand efficacy | MSP variant | Lipid composition | Relative populations | | | | |
|---|---|---|---|---|---|---|---|---|
| | | | | P4 | P3 | P2 | P1 | P5 |
| NECA | Full agonist | MSP1D1 | POPC:POPS (85:15) | | 0.52 | | 0.31 | 0.15 |
| NECA | Full agonist | R85Q | POPC:POPS (85:15) | 0.26 | | 0.30 | 0.35 | 0.07 |
| NECA | Full agonist | R140Q | POPC:POPS (85:15) | 0.37 | | 0.43 | 0.15 | 0.03 |
| NECA | Full agonist | R30Q | POPC:POPS (85:15) | 0.21 | | 0.26 | 0.50 | 0.009 |
| NECA | Full agonist | R118Q | POPC:POPS (85:15) | 0.20 | | 0.27 | 0.39 | 0.11 |
| NECA | Full agonist | R118Q+R140Q+ K87Q | POPC:POPS (85:15) | 0.49 | | 0.39 | 0.07 | 0.03 |
